# An atlas of Arabidopsis protein S-Acylation reveals its widespread role in plant cell organisation of and function

**DOI:** 10.1101/2020.05.12.090415

**Authors:** Manoj Kumar, Paul Carr, Simon Turner

## Abstract

S-acylation is the addition of a fatty acid to a cysteine residue of a protein. While this modification may profoundly alter protein behaviour, its effects on the function of plant proteins remains poorly characterised, largely as a result to the lack of basic information regarding which proteins are S-acylated and where in the proteins the modification occurs. In order to address this gap in our knowledge, we have performed a comprehensive analysis of plant protein S-acylation from 6 separate tissues. In our highest confidence group, we identified 5185 cysteines modified by S-acylation, which were located in 4891 unique peptides from 2643 different proteins. This represents around 9% of the entire Arabidopsis proteome and suggests an important role for S-acylation in many essential cellular functions including trafficking, signalling and metabolism. To illustrate the potential of this dataset, we focus on cellulose synthesis and confirm for the first time the S-acylation of all proteins known to be involved in cellulose synthesis and trafficking of the cellulose synthase complex. In the secondary cell walls, cellulose synthesis requires three different catalytic subunits (CESA4, CESA7 and CESA8) that all exhibit striking sequence similarity. While all three proteins have been widely predicted to possess a RING-type zinc finger at their N-terminus, for CESA4 and CESA8, we find evidence for S-acylation of cysteines in this region that is incompatible with any role in coordinating metal ions. We show that while CESA7 may possess a RING type domain, the same region of CESA4 and CESA8 appear to have evolved a very different structure. Together, the data suggests this study represents an atlas of S-acylation in Arabidopsis that will facilitate the broader study of this elusive post-translational modification in plants as well as demonstrates the importance of undertaking further work in this area.

## Introduction

S-Acylation, also known as palmitoylation, is a reversible post-translational modification involving the transfer of a fatty acid group, frequently stearate or palmitate, to a cysteine residue in target proteins. The very hydrophobic nature of the acyl group can dramatically alter the protein properties. In proteins that lack transmembrane helices, the addition of an acyl group will confer membrane localisation, while for proteins that already possess transmembrane helices, addition of an acyl group has a variety of effects including altering their subcellular localisation, partitioning into membrane microdomains and enabling the formation of multi-protein complexes ^1,2^. The extent of S-acylation in eukaryotic cells has been revealed by studies on yeast and mammalian cells. An estimate based on the proteins in SwissPalm, the recently established database of protein palmitoylation, suggests that 25% of mammalian proteins are likely to be modified by S-acylation and even this is likely to be an underestimate ^3^.

The acyl group is added by a family of protein acyl transferases (PATs) that are characterised by a DHHC motif within their catalytic domain. The structure of a human PAT, DHHC17, has recently been solved ^4^, but despite this and the identification of a large number of S-acylation sites, there are no recognisable motifs around the modified cysteines of the substrate proteins. A recent study that monitored S-acylation directly using mass spectrometry (MS) suggested that cysteines were modified independently of any sequence motif around the site, in a stochastic process that depends upon the accessibility of any given cysteine to the action of a PAT ^5^. Consequently, making accurate bioinformatics prediction of S-acylation sites remains very challenging and knowledge of the extent of S-acylation is dependent on empirical identification of S-acylation sites.

The plant “acylome” has been partially described by two proteomic studies from Arabidopsis seedlings ^6^ and poplar suspension cultures ^7^ that identified a total of 694 and 449 proteins respectively with various levels of confidence. Neither study identified the sites of the acyl group attachment and as a consequence acyl-sites have been identified for only a relatively small number of individual plant proteins. The lack of information about acyl-sites of plant proteins is particularly acute in comparison to the information available from mammalian systems ^8-11^ though the importance of S-acylation has been demonstrated for a small number of proteins ^2,12-14^. These proteins include heterotrimeric G proteins, small G proteins, proteins involved in Ca signalling, proteins involved in pathogenesis, transcription factors and CESA proteins. Functional analysis on the role of S-acylation in the absence of information about the sites of modification is challenging. A study of the catalytic subunit of the plant cellulose synthase complex, AtCESA7, identified 6 S-acylated cysteines organised in two clusters^15^. This work required the use of systematic mutagenesis, which for a protein with 26 cysteines, is a laborious approach and does not guarantee the identification of S-acylation sites. While the limited knowledge of acylation sites has hampered studies on the role of S-acylation in the function of plant proteins, analysis of the individual Arabidopsis PATs has implicated S-acylation in a variety of processes including root hair formation, cell death, ROS production and branching, cell expansion and division, gametogenesis and salt tolerance ^2,12-14,16^. So while our understanding of both the extent and function of S-acylation of individual plant proteins remains limited, the available data indicates S-acylation is important for many aspects of plant cell function.

To meet the need for more information on both the true extent of S-acylation of plant proteins and the sites at which the modifications occur, we undertook a comprehensive analysis of S-acylation from 6 Arabidopsis tissues. Within our high confidence group, we identify evidence for a total of 5185 cysteines modified by S-acylation which were located in 4891 unique peptides from 2643 different proteins. We verify the accuracy of the data and demonstrate its utility by focussing on cellulose biosynthesis and identify that S-acylation appears to be a general feature of proteins involved in cellulose biosynthesis. Furthermore, we demonstrate how functional analysis of this data provides new insights into protein structure. In particular, the function of the “RING finger” domain has diverged between different CESA isoforms, a finding that alters our understanding of how this domain functions during cellulose synthesis. Together, this study represents an important milestone in the study of S-acylation of plant proteins, demonstrates the extent of S-acylation of plant proteins and provides a unique dataset that may be used as a resource for the functional analysis of S-acylation of plant proteins.

## Results and discussion

### S-Acylated proteins from 6 Arabidopsis tissues

The identification of S-acylated proteins was performed using Acyl-RAC assay ^17^ that relies on capture of S-acylated peptides on beads in a hydroxylamine dependent manner. We optimised the protocol by incorporating chemical scavenging of NEM with 2-3 dimethyl-1-3 butadiene ^18^, thereby avoiding the need for protein precipitation steps. Furthermore, we have previously shown that during membrane preparation and solubilisation with non-ionic detergents, a large fraction of CESA proteins remains with the cell wall debris ^19^ suggesting that this approach may bias the composition of the starting material. Consequently, we solubilised the tissue powder directly in a buffer containing 2.5% SDS. After blocking free cysteines, we captured S-acylated proteins by hydroxylamine (HA) dependent binding to thiol reactive beads. After stringent washes, we performed on-bead trypsin digestion of the S-Acylated proteins. The peptides containing the acyl cysteines remained bound to the beads and were released in a DTT dependent manner (Figure S1). We analysed 3 biological replicates each from Arabidopsis material collected from: 7 day old whole seedlings, and 5 tissues collected from 5-week old plants: mature stems, hypocotyl, siliques, rosette leaves and inflorescence meristem.

MS data was analysed using Proteome Discoverer and Microsoft excel (see methods for details). The entire data from all tissues consisted of 176373 peptide spectral matches (PSMs) with associated intensity, peptide sequence and peptide modifications. This data was converted into normalised and log2 transformed intensity data for a total of 11219 peptide forms. A PCA plot using this data (Figure 1A) indicated a good clustering of the biological replicates and a good separation between samples treated with HA and controls that were untreated. Any peptides not containing a carbamidomethyl cysteine were removed and the remaining 10705 peptides were then matched to Araport11 proteins. 9186 peptides had a single unambiguous database match (exclusive peptides), representing 4240 proteins. A further 1519 peptides matched to more than one protein (ambiguous peptides) that mapped up to 1816 proteins. We then calculated the protein intensities by taking the sum of intensities of peptides matching to each protein. This was done for exclusive and ambiguous peptides separately.

**Figure 1.**
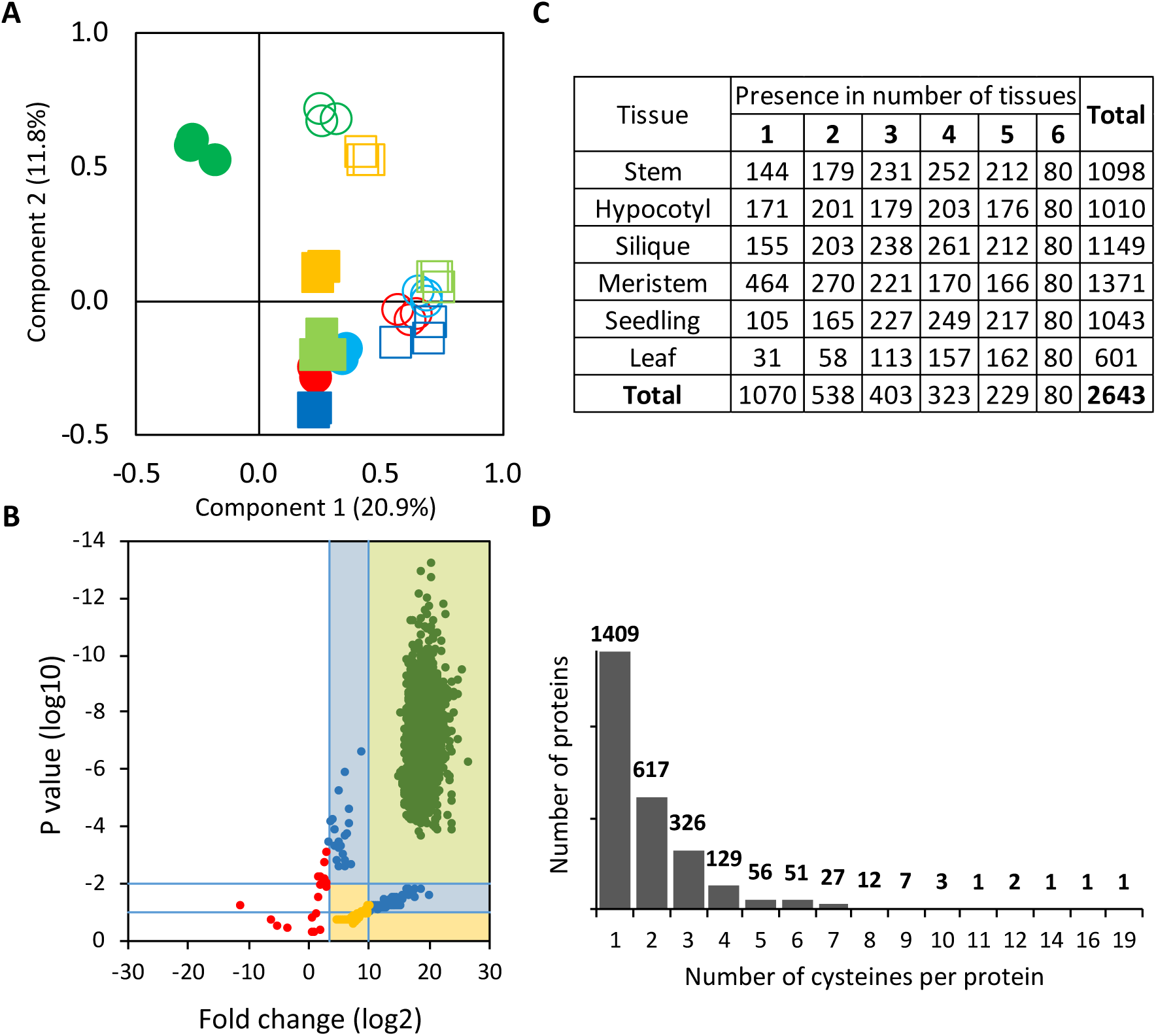
Overview of the Arabidopsis Acylome. A) PCA analysis of peptide data from extracts of stem (red circles), hypocotyl (green circles), silique (pale blue circle), meristem (yellow square), seedlings (pale green squares) and leaf (dark blue squares). Extracts were treated with (solid) or without (open) hydroxylamine. (B) Fold change (log2) and P value (log10) derived from peptide intensities of stem sample treated with HA compared to untreated controls. The shaded areas of the plot indicate the way in which the peptides were classified into different groups with 1 (light green shading), 2 (light blue shading) and 3 (light yellow shading) referring to the high, medium and low confidence groups. Peptides in the unshaded area (red dots) were not considered further (see text for details). (C) Distribution of acylated proteins across tissues. Out of 2643 high confidence proteins, 1070 were detected in only one tissue while 80were detected in all 6 tissues. (D) Distribution of the number of S-acylated cysteines per protein among those proteins from the high confidence class.

We calculated the fold change and p value for individual peptides by comparing the intensities in the HA treated samples with those of control samples in each tissue. When the fold change was plotted against P value, the bulk of the peptides formed a distinctive group that exhibited both a high fold change and low P value (Figure 1B, Figure S2A). We defined this high confidence group as peptides that exhibited a fold change of greater than 1000 fold (log2 = 9.97) and a P value of less than 0.01 (Figure 1B, Table 1). We also defined a medium confidence group as those peptides that either exhibited a fold change of more than 1000, but a P value between 0.1 and 0.01 or those a P value of less than 0.01, but a fold change of between 10 and 1000. While the low confidence group comprised peptides with a fold change of between 10 and 1000, but a P value of less than 0.10 (Figure 1B, Table 1, Figure S2A). We were able to plot a similar distribution for protein intensities (Figure S2B) and in most cases, the protein score matched to the score of the best peptide mapped to that protein. The final protein grouping was assigned by comparing the protein score with the score of the best peptide and taking the lower of the two. Therefore, a protein in the high confidence group must also possess a peptide from the high confidence group. The peptides and proteins in the high confidence group are detected in all three biological replicates of the HA treated sample but were undetectable in the untreated control. At this stage, we also filtered out 43 proteins that are known to contain thio-ester bonds unrelated to S-acylation ^6^.

**Table 1.** Summary of proteins, peptides and cysteines in within confidence groups. The data is based on exclusive or ambiguous peptides. In the peptide table, each ambiguous peptide is counted only once and hence reflects the actual number of peptides detected. For the protein and cysteine table, on the other hand, the numbers in ambiguous categories are likely to be overestimates as all proteins represented by the ambiguous peptides are counted. The data is shown for each of the 6 tissues analysed. The last column “best” indicates the best confidence for a peptide/protein/cysteine across all 6 tissues.

|  | Peptide Type | Confidence Group | Tissue |  |  |  |  |  |  |
| --- | --- | --- | --- | --- | --- | --- | --- | --- | --- |
|  |  |  | Stem | Hypocotyl | Silique | Meristem | Seedling | Leaf | Best tissue |
| Proteins | Exclusive | High | 1098 | 1010 | 1149 | 1371 | 1043 | 601 | 2643 |
|  | Exclusive | Medium | 461 | 510 | 574 | 867 | 605 | 405 | 796 |
|  | Exclusive | Low | 486 | 393 | 479 | 676 | 613 | 384 | 743 |
|  | Ambiguous | High | 378 | 379 | 401 | 394 | 369 | 264 | 716 |
|  | Ambiguous | Medium | 178 | 244 | 220 | 306 | 191 | 156 | 214 |
|  | Ambiguous | Low | 154 | 160 | 142 | 151 | 247 | 145 | 150 |
| Peptides | Exclusive | High | 1744 | 1618 | 1893 | 2290 | 1529 | 892 | 4972 |
|  | Exclusive | Medium | 1004 | 851 | 1134 | 1441 | 1135 | 711 | 1953 |
|  | Exclusive | Low | 1091 | 840 | 1179 | 1320 | 1366 | 814 | 2043 |
|  | Ambiguous | High | 383 | 344 | 455 | 397 | 369 | 243 | 940 |
|  | Ambiguous | Medium | 203 | 199 | 238 | 245 | 206 | 174 | 295 |
|  | Ambiguous | Low | 168 | 160 | 178 | 174 | 251 | 145 | 247 |
| Cysteines | Exclusive | High | 2066 | 1789 | 2047 | 2485 | 1682 | 1012 | 5279 |
|  | Exclusive | Medium | 1114 | 851 | 1171 | 1465 | 1197 | 707 | 1935 |
|  | Exclusive | Low | 1152 | 867 | 1217 | 1312 | 1422 | 852 | 2007 |
|  | Ambiguous | High | 1000 | 889 | 1111 | 997 | 932 | 601 | 2088 |
|  | Ambiguous | Medium | 436 | 423 | 464 | 550 | 421 | 389 | 600 |
|  | Ambiguous | Low | 356 | 346 | 340 | 328 | 520 | 317 | 470 |

Unless otherwise stated, we have focussed on proteins and peptides belonging to the high confidence group with a P-value of less than 0.01 and fold change of >1000. The data for the lower confidence groups is included in Table S1 and may contain important information regarding acylation sites that is useful for data mining, but it is likely that the rate of false positives in these lower confidence groups may be higher.

### Comparison with previous studies of the plant acylome

In order to establish the accuracy of our acylome data set, we compared it with the only previous proteomics study of S-Acylated proteins in Arabidopsis ^6^ that identified a total of 582 acylated proteins from seedling tissue that were classified as either high, (122) medium (94) or low confidence (336). We identified the majority of these proteins in our study, with more than 65% of their high and medium confidence groups also identified in our study.

The most comprehensive acylomes available in SwissPalm are for human and mouse ^3^. We identified putative homologs of our 2643 S-acylated Arabidopsis proteins in these two species and then checked whether or not there was evidence for acylation of those human and mouse proteins. We identified human homologs for 1556 proteins (using an e value of -10), 878 (56%) of which were known to be acylated in one of the 17 studies available ^3^. The corresponding figures for mouse were 1570 and 819 (52%) respectively. While this number was higher than expected, it remains to be established whether this is the result of convergent evolution.

One of the major advances of this study is to identify sites of acylation. To confirm the accuracy of our data, we compared our dataset to the relatively small number of S-acylation sites experimentally verified for plant proteins ^20^. In addition to CESA proteins (see below), we also confirmed previously published acylation sites on several Rho GTPases. These proteins contain no transmembrane spanning helices, but association with membranes occurs via lipid modifications that are essential for signalling. In Arabidopsis there are 11 Rho of plants (ROPs) that fall in 2 classes ^21^. The Type-I ROPs are reversibly S-acylated at 2 internal cysteines, C21 and C158. ^22,23^, while Type-II ROPs are S-acylated at the C-terminal cysteines ^24^. Our data confirmed the S-acylation of the C terminal cysteines in the type II ROPS (10 and 11) as well as identifying the cysteine corresponding to C158 in 6 type I (ROP1 to ROP6) and 1 type II (ROP11) (Figure S3A). Additionally, we identified an ambiguous peptide which maps to all 11 ROPs and contains a cysteine corresponding to C21 for ROP8 (Figure S3A). RPM1 interacting protein 4 (RIN4) is a small protein which forms a central component of plant defence that associates with the membrane as a result of S-acylation of three C-terminal cysteines ^25^. In our acylome, we identified a single RIN peptide containing these 3 cysteines indicating that for both the relatively well-characterised ROPs and RIN4 we have successfully identified their known S-acylation sites. Together these comparisons illustrate that our acylome dataset contains most of the previously identified plant S-acylated proteins and has captured most of the known S-acylation sites in those proteins.

### Experimental verification of the acylome

We have previously demonstrated that CESA proteins, the catalytic subunits of the cellulose synthase complex, are extensively modified by S-acylation ^15^. The CESA proteins were well represented in our dataset (see below); however, we also identified several other proteins that are known to be involved in cellulose synthesis. This included the endoglucanase KORRIGAN (KOR1), CELLULOSE SYNTHASE INTERACTING (CSI) and CELLULOSE MICROTUBULE UNCOUPLING (CMU) that all interact with the cellulose synthesis complex and/or microtubules ^26,27^. Furthermore within the acylome we also identified a number of proteins known to be involved in CSC trafficking (DET3, PATROL1), exocytosis (EXO70A1, EXO84B, SEC6 and SHOU4) and endocytosis (AP2M/µ2, TML, TPLATE and TWD40-2) (reviewed in ^28^). 10 of these proteins were identified with high confidence in at least one tissue (Table S2). In order to independently verify S-acylation of these proteins, we performed ACYL-RAC assays followed by immunoblotting using proteins transiently expressed in Tobacco and/or in stably transformed lines. These assays verified that KOR, CSI, CMU1, CMU2, SHOU4 and SHOUL4 were clearly modified by S-acylation, while our negative controls were not (Figure 2A-B). Since the data suggests all cellulose synthesis related proteins might be modified by S-acylation, we hypothesised that another protein that is known to be associated with CSC but not identified in our data, known as COMPANION OF CELLULOSE SYNTASE (CC) would also be S-acylated. Acyl-RAC assays followed by immunoblots revealed that both CC1 and CC2 are indeed S-acylated (Figure 2C).

**Figure 2.**
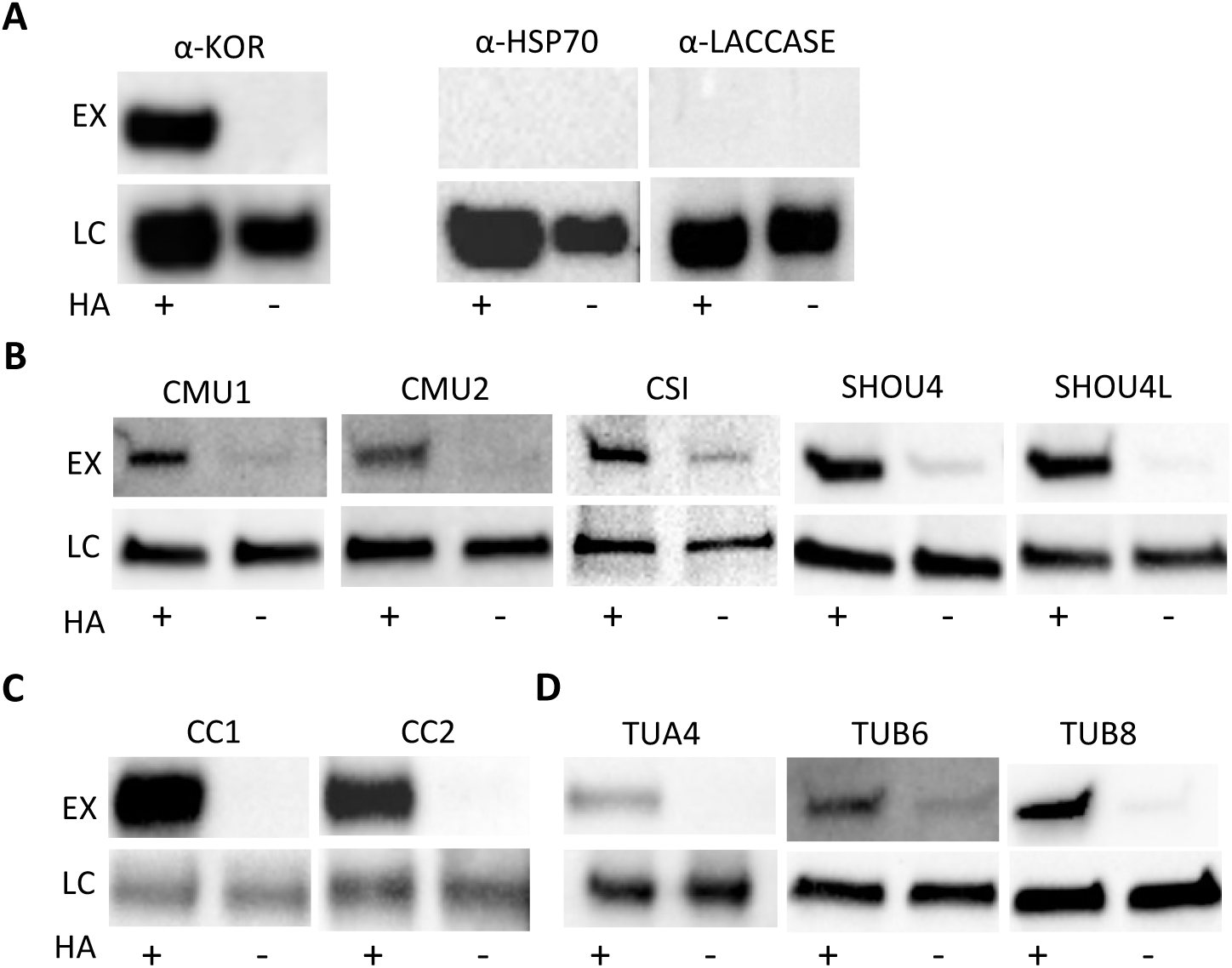
Validation of S-acylated proteins using Acyl-RAC assay and immunoblotting. For each assay, the experimental sample (EX) was compared with the loading control (LC) with or without (+/-) hydroxylamine (HA) for hydroxylamine-dependent capture of S-acylated proteins. (A) Individual proteins extracted from stems were detected with specific antibodies. (B-D) Immunoblots of extracts containing GFP-fusion proteins expressed either transiently in tobacco using 35S promoter (SHOU4, SHOU4L and TUA4) or stably in Arabidopsis under the control of native promoters (CMU1, CMU2, CSI, CC1 and CC2) probed with an anti-GFP antibody.

CSI, CC and CMU all associate with microtubules as well as the cellulose synthase complex. In common with previous reports ^29^, many tubulin subunits were found in our acylome dataset. However, the conserved nature of these proteins, especially alpha tubulins, makes it hard to distinguish individual family members by MS. We selected two alpha tubulins, TUA1 and TUA4, and two beta tubulins, TUB6 and TUB8, based on their expression in secondary wall forming tissues. We expressed GFP fusions of all 4 proteins transiently in tobacco leaves and/or in stable transgenic plants. Acyl-RAC assays confirmed that TUB6, TUB8, TUA1 and TUA4 were all S-acylated (Figure 2D).

KOR1 is known to associate with the CESA proteins in the cellulose synthase complex ^30^. KOR1 was found in our acylome data in all 6 tissues, but only a single S-acylated site (C64) was detected. The presence of a single S-acylation site was examined using a variation of the Acyl RAC assay, known as PEGylation ^31^, in which the acyl groups is exchanged for PEG of sufficiently large size to cause a noticeable mobility shift on SDS-PAGE. Using this assay, the KOR1 antibody detected a dimer that is likely to represent the unmodified form and the modified form with a single PEG addition (Figure 3A) suggesting that KOR does only possess a single site for S-acylation. In order to verify the acyl site, we mutated the C64 to a serine and generated stably transformed lines. Acyl-RAC assays clearly demonstrated the mutation abolished S-acylation (Figure 3B). We measured root length to assess the *kor* phenotype, but we were unable to identify a significant defect associated with the *KOR1*^*C64S*^ mutant (Figure 3C, Figure S4) suggesting that the phenotype is likely to be subtle or only apparent under certain conditions. C64 is located close to the TM domain of KOR and previous studies have also failed to identify a phenotype associated with mutations in these juxta-membrane sites ^18^. Nevertheless, the correct identification of sites of S-acylation that together with the identification of previously described S-acylated proteins provides independent verification that the dataset presented here represents an accurate picture of S-acylation of Arabidopsis proteins.

**Figure 3.**
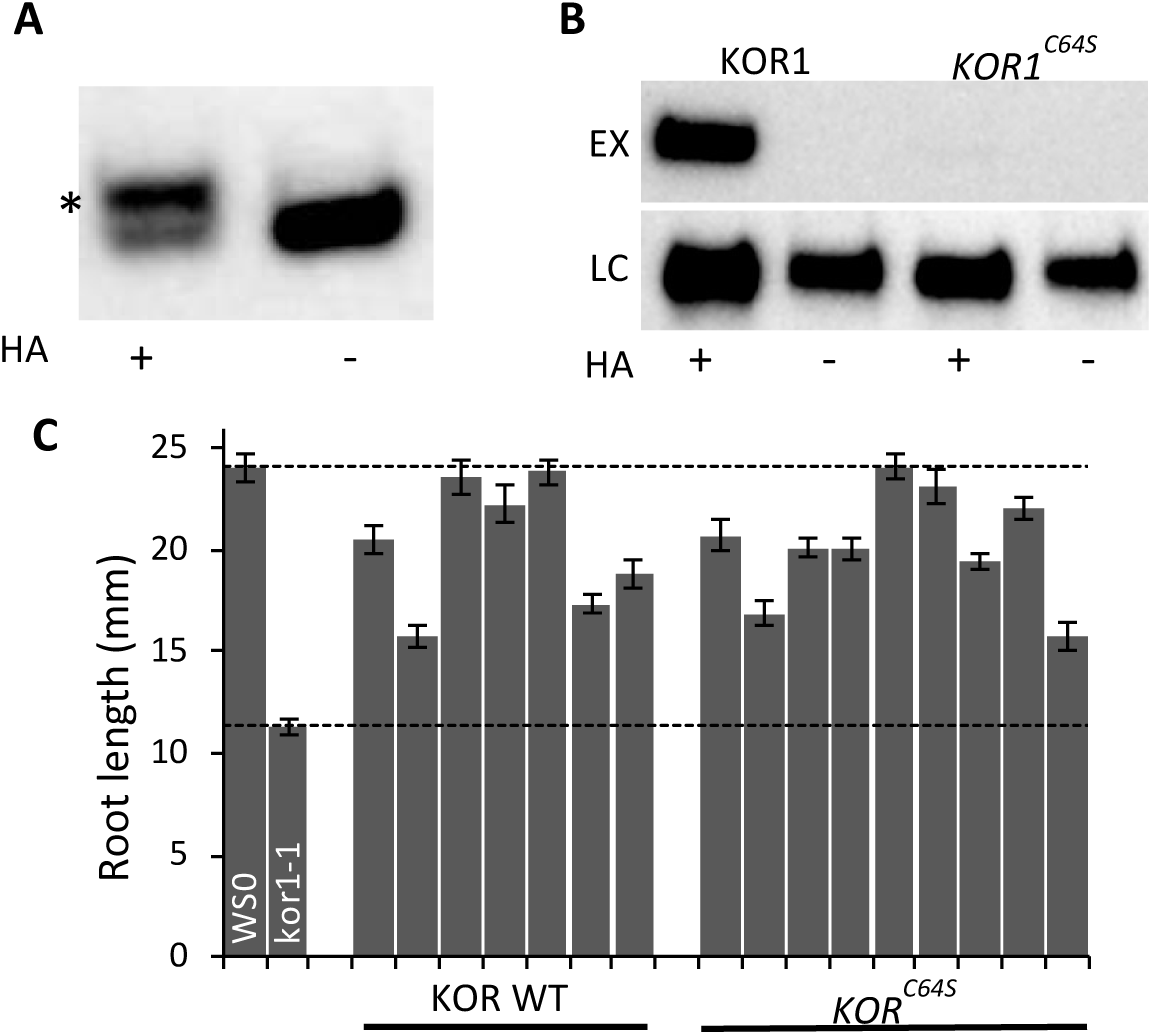
S-Acylation of KOR1. (A) Acyl-PEG exchange (APE) assays were performed on wild type stem material using 5 kD mPEG-maleimide and analysed on immunoblots probed with anti-KOR1 antibody. Assay was performed with or without hydroxylamine (HA) for hydroxylamine-dependent PEG addition. The band exhibiting decreased mobility on the gel as a result of PEG addition is indicated (*). (B). Acyl-RAC assays were performed on 7 day old *kor1-1* seedling mutants transformed with GFP fusions of either wild type KOR1 or the *KOR1*^*C64S*^ mutant expressed under control of the native promoter. The experimental sample (EX) was compared with the loading control (LC) with or without (+/-) hydroxylamine (HA) for hydroxylamine-dependent capture of S-acylated proteins. Proteins were detected using an anti-GFP antibody. (C) The root length of wild type WS0, *kor1-1* and independent T2 lines of *kor1-1* transformed with either the wild type KOR or the *KOR1*^*C64S*^ mutant. At least 30 roots were measured for each line using ImageJ. Error bars shown are standard error of mean. The dashed horizontal lines indicate root lengths of wild type WS0 and *kor1-1*.

### Overview of Acylated proteins

In order to get a crude estimate of the proportion of the proteome modified by S-acylation, we compared our data to a recent draft of the Arabidopsis proteome ^32^. They identified a total of 18175 proteins from 30 different tissues. Analysis of this data suggests that most proteins are present in multiple tissues, so we compared our acylome from all tissues to the entire proteome. Our high confidence group (2643) represents 14% of the entire proteome. This is lower than estimates of the human acylome that suggests up to 25% of proteins may be acylated; however, our acylome data is unlikely to be exhaustive. If we include proteins from our high medium and low confidence class (4182 proteins), we have a figure of 22% much closer to that suggested for mammalian cells.

Within the high confidence group, a total of 2643 proteins were identified in one or more tissues (Table 1, Figure 1C). A comparison of the overlap between tissues shows that out of 2643, 1070 proteins were identified in only one tissue, 464 of which were contributed by meristem reflecting a unique acylome in this tissue (Figure 1C). The remaining 1573 proteins were detected in more than one tissue, with 80 proteins being identified in all 6 tissues.

In order to understand how S-acylation might affect protein localisation, we analysed each protein for presence or absence of transmembrane helices (TMHs) using Phobius and predicted subcellular localisation using SUBA4 (Table 2). One of the striking features of the data was the very high proportion of proteins that lack TMHs. Of the 2643 proteins, 2193 (83%) lack TMHs suggesting that S-acylation will have a large influence on the cellular distribution of the plant proteome. Of the 245 proteins that were predicted to be localised to plasma membrane in SUBA4, 131 (53%) lack any transmembrane helices and consequently S-acylation is likely to play an important, if not essential, role in anchoring many of them to the plasma membrane.

**Table 2.** Prediction of sub-cellular localisation of high confidence proteins. Proteins from high confidence class were analysed for subcellular localisation using SUBA4 and presence of trans-membrane helices (TMHs) using Phobius.

| SubaCon localisation prediction | All | TMH | No_TMh | No_TMh (%) |
| --- | --- | --- | --- | --- |
| cytosol | 605 | 21 | 584 | 97 |
| plastid | 514 | 59 | 455 | 89 |
| nucleus | 347 | 19 | 328 | 95 |
| plasma membrane | 245 | 114 | 131 | 53 |
| extracellular | 209 | 20 | 189 | 90 |
| mitochondrion | 197 | 24 | 173 | 88 |
| golgi | 167 | 71 | 96 | 57 |
| endoplasmic reticulum | 93 | 53 | 40 | 43 |
| peroxisome | 77 | 10 | 67 | 87 |
| vacuole | 57 | 35 | 22 | 39 |
| nucleus, cytosol | 51 | 3 | 48 | 94 |
| mitochondrion, plastid | 11 | 0 | 11 | 100 |
| plasma membrane, cytosol | 11 | 1 | 10 | 91 |
| endoplasmic reticulum, golgi | 7 | 2 | 5 | 71 |
| endoplasmic reticulum, plasma membrane | 7 | 4 | 3 | 43 |
| golgi, cytosol | 5 | 0 | 5 | 100 |
| Other | 40 | 14 | 26 | 65 |
| **Total** | **2643** | **450** | **2193** | **83** |

We next looked at functional classification of acylated proteins using Mapman bins ^33^. We performed enrichment analysis in various protein lists including all S-acylated proteins (2643) and all S-acylated proteins within each tissue (Figure 4). Functional classes associated with metabolism of all major biomolecules were among the significantly enriched, indicating the diversity of processes involving S-acylation. There appears to be distinct specificity within the bins. For example, within the large group enriched for proteins involved in vesicle trafficking (Figure 4.), we find sub-groups containing machinery for clathrin coated vesicle and Coat protein II (COPII) coatomer machinery are enriched while those for coat protein I coatomer machinery are not (Table S3).

**Figure 4.**
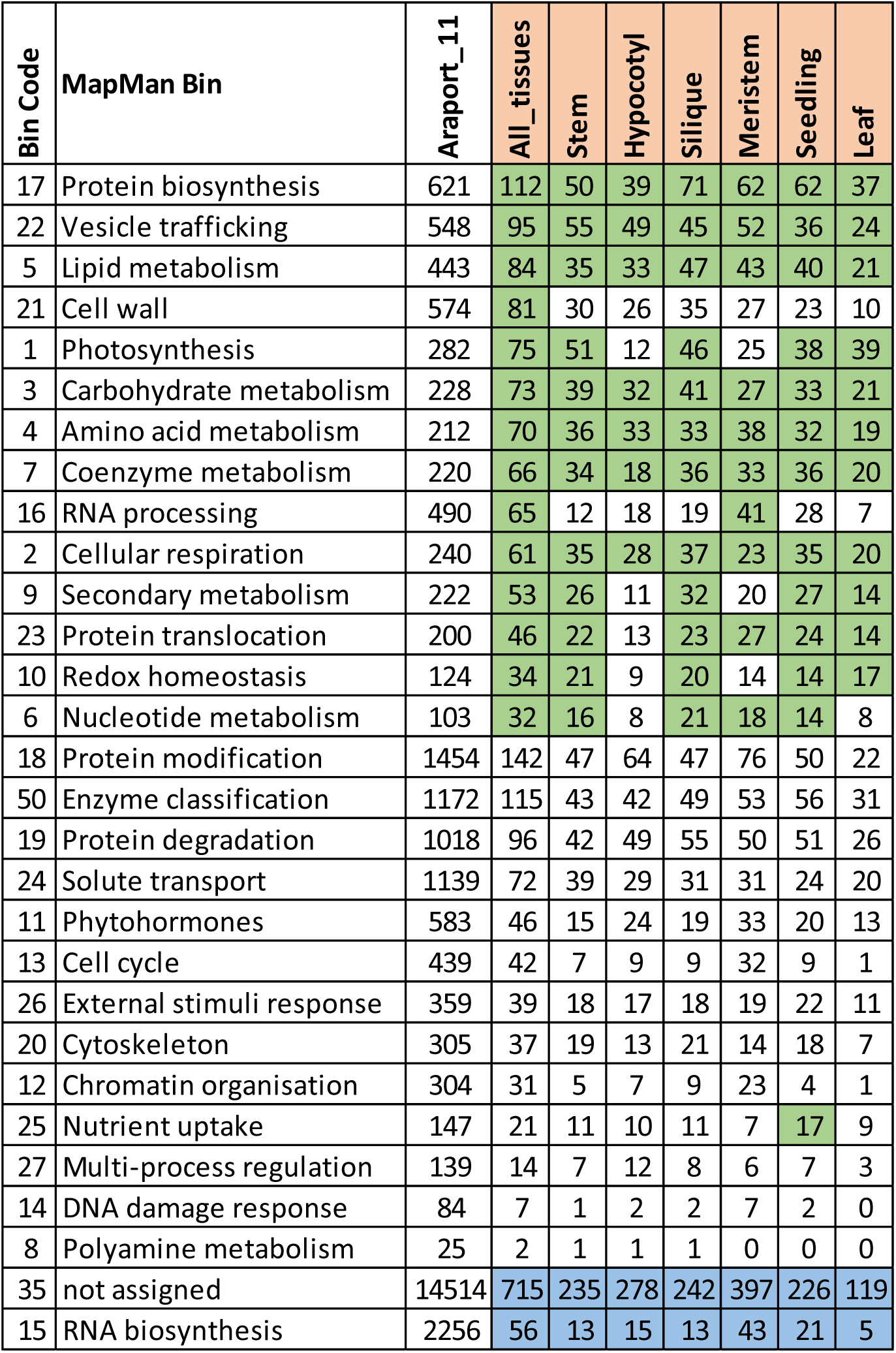
Functional classification of S-Acylated proteins. Proteins identified with high confidence in at least one of the 6 tissues (All_tissues) or each individual tissue were subjected to over-representation analysis using Mapman categories. The total number of proteins for each category in whole proteome (Araport11) and the acylomes from all or individual tissues are shown. Green shading indicates over representation with a P value less than 0.05 while blue shading indicates under-representation.

### Overview of Acyl-Sites

To be classed into the high confidence group, by definition a protein must have at least one peptide and by extension one cysteine with a p value of <0.01 and fold change >1000. The 2643 proteins in this group contained 5185 high confidence cysteines and while many of the proteins (1409) contained only a single high confidence cysteine, the remainder (1234) contained multiple cysteines (Figure 1D).

There were 28 proteins for which we identified 8 or more of the high confidence cysteines. These heavily acylated proteins included many proteins involved in cellulose synthesis like CESA4, CESA8 and CESA3 (see below) and CSI1 adding further support to the idea that S-acylation is particularly important during cellulose synthesis.

A number of algorithms have been developed for prediction of S-acylation sites. We used CSS-PALM 4.0 ^34^ to predict S-acylation sites in our high confidence protein set (2643 proteins) and then we compared the overlap between the predicted sites and sites identified in this data. Out of 5185 high confidence sites experimentally identified in our data, only 582 were predicted by CSS-PALM using the high threshold settings. A further 444 and 261 sites were predicted using medium and low settings respectively making a total of 1287 sites out of 5185 (25%) being predicted by CSS-PALM (Table S4). This result demonstrates that while useful, the training sets used for these programs are not adequate, especially for plant proteins where the sites are known only for a small number of proteins.

To analyse whether any consensus sequence may exist around S-acylation sites, we divided the proteins in two groups based on whether they contain a transmembrane helix or not. In neither of these classes did we identify any clear sequence consensus, but consistent with previous studies ^8,35^ we did identify very simple sequence motifs enriched in a limited subset of proteins (Figure S5).

### S-acylation of protein kinases

A recent detailed study of S-acylation of LRR protein kinases analysed 7 well characterised proteins and found them all to be S-acylated, leading to the suggestion it may be a feature of all receptor kinases ^18^. Hence, we performed a systematic search of LRR and other classes of protein kinases in our data. In general, kinases were not over-represented in our data (Table S5). Our inability to detect S-acylation for many protein kinases may mean that they are not modified in this way, but it may also be a consequence of their relatively low abundance in the tissues we examined. It may also be a result of many kinases exhibiting dual modifications ^36^, most of which are not identified in our analysis (see below). There are, however, several protein kinase families that are well represented in our data. The most abundant kinases represented are members of the raf family of kinase, but there is little information on the function or structural characterisation of these kinases with regard to any lipidation. Calcium dependent protein kinases (CDPKs or CPKs) are a family composed of 34 members in Arabidopsis ^37^ that are myristoylated at the N-terminal glycine, an amide (N-) linked acyl modification that is not detected with the Acyl-RAC assay. While many CPKs are also predicted to be S-acylated at the N-terminus, to date this has only been verified biochemically for AtCPK6 ^36^. S-acylation sites adjacent to myristoylation sites are poorly represented in our data, however we do find that all 10 CPKs found in our high confidence group were modified at the beginning of the kinase domain (Figure S3B). For some members additional sites were also identified in the middle (5 CPKs) and end (2 CPKs) of kinase domain suggesting a more complex pattern of S-acylation.

Brassinosteroid signalling kinases (BSKs) are a family of proteins that lack transmembrane helices but facilitate brassinosteroid signalling by binding to the receptor (BRI) ^38^. It was recently reported that BSKs are membrane bound as a result of myristoylation at the N-terminus ^39,40^. 6 of the 11 BSK family members are represented in our data and all share a common site of S-acylation in the kinase domain (Figure S3B). For membrane-associated proteins, myristoylation frequently precedes S-acylation and it is probable that dual myristoylation and S-acylation are required for proper function on BSKs. In addition to BSKs, we found evidence for S-acylation of a number of other proteins required for brassinosteroid signalling and perception. The high confidence group contains BSL1 and there are multiple ambiguous peptides matching to BSL1/BSL2/BSL3. BSL proteins are related to BRI1 Suppressor1 (BSU1), a protein that acts in the BR signalling pathway downstream of BSK. Additionally, within our low confidence group, we also found a single acylated peptide located in kinase domain on the brassinosteroid receptor BRI1, and two highly conserved ambiguous peptides that map to a group of 10 related shaggy-like kinases. Seven of these kinases, which include BIN2, are known to play an essential role in BR signalling ^41^. Together, this suggests that S-acylation may be a common characteristic of proteins involved in BR signalling; a hypothesis that is now relatively easy to test using the data provided in this study.

### Functional analysis reveals diversification of RING domain function between CESA classes

To demonstrate how these data may require us to re-think the structure and function of some protein domains, we looked more closely at how the CESA proteins are S-acylated. We previously identified 4 cysteines in the variable region 2 (VR2) for AtCESA7 as major site of S-acylation ^15^. The acylome data presented here confirmed that the VR2 cysteines of CESA7, CESA4, and CESA8 are all acylated. Furthermore, we identified several additional novel acylation sites for CESA proteins (Figure S3D). In particular, we identified a number of highly conserved cysteines close to the N-terminus. These sites are part of a motif containing 8 cysteines that is highly conserved in all CESA proteins, and also in the related CSLD group of glycosyltransferases. This motif is frequently described as a RING-type zinc finger ^42^. A structure for this region of CESA7 has been solved by NMR and demonstrates it to be a zinc binding RING domain (PDB 1WEO) (Figure 5A). Even though these proteins exhibit high overall similarity in this region and were previously considered to be orthologous domains ^42,43^ (Figure 5C), we identified no acylation sites in the RING domain of CESA7, but we identified 7 cysteines of AtCESA8 and 3 cysteines of AtCESA4 in either the high or medium confidence dataset (Figure S3D). Acylation of these cysteines is not compatible with these regions forming a Zn-binding RING domain and suggests a very clear divergence in the structure, and probably the function, of this domain between CESA7 and CESA4/8. We also identified S-acylated cysteines in the RING domain of CESA3, but not CESA1 or CESA6 suggesting that primary wall CESA proteins may also have diverged in a similar way. There are a large number of RING-type zinc fingers in Arabidopsis. Out of 487 RING domain proteins, apart from 3 CESA proteins, we only detected modification of the RING domain cysteines in 5 other proteins suggesting this is a rare modification of such structures.

**Figure 5.**
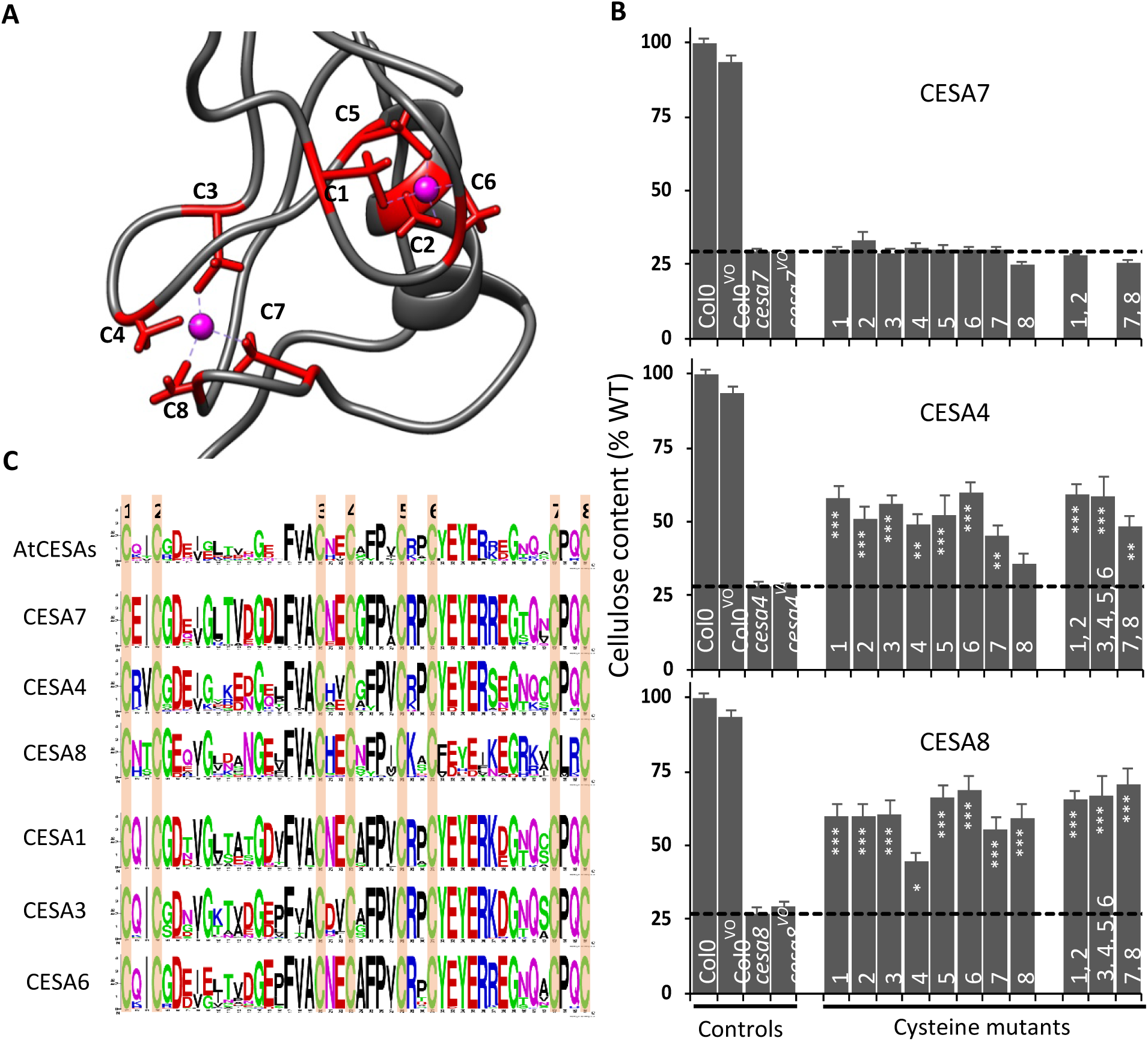
Analysis of RING finger cysteines in CESA7, CESA4 and CESA8. Part of the structure of RING-finger of CESA7 (1WEO) indicating the 8 cysteines (red sticks) which coordinate two Zn atoms (magenta spheres). The cysteine numbering is the same as that used in parts B and C. (B) Cellulose content of *cesa7*^*irx3-7*^, *cesa4*^*irx5-4*^ and *cesa8*^*irx1-7*^ mutants transformed with the corresponding genes in which one or more cysteines have been mutated to serines. For each genotype at least 9 plants were analysed. The suffix VO indicates plants transformed with empty vector only. The numbers at the base of each bar refer to which cysteines are mutated and correspond to those shown in part A. Horizontal dashed line indicates cellulose content of background mutants. Error bars shown are standard error of mean. Significance levels at 0.001 (***), 0.01 (**) and 0.05 (*) are shown for comparison of cysteine mutants with the respective T-DNA mutants and were generated using univariate ANOVA. (C) Sequence analysis of zinc finger domain of CESA proteins. Weblogos were generated using either all 10 Arabidopsis CESAs (top row) or for each CESA protein’s class using CESA sequences identified in 42 fully sequenced plant genomes. The 8 conserved cysteines that are proposed to coordinate the zinc ion are indicated.

To test any differences in the function of this N -terminal domain between different isoforms, we performed a comprehensive mutagenesis of these cysteines in CESA4, CESA7 and CESA8. Initially, each individual cysteine was mutated to serine and the constructs transformed into their respective null mutants. Consistent with the known structure (Figure 5A) and the essential nature of the RING domain in CESA7, ^15^, mutating any single cysteine is sufficient to completely abolish any complementation of either the plant height (Figure S6) or cellulose content phenotype (Figure 5B). In contrast, when we mutate the same cysteines of CESA4, we observe partial but significant complementation of cellulose content and plant height. We obtain the same results when we analyse these mutants in CESA8; all constructs exhibit significant complementation of the *cesa8* null mutant. To confirm that our mutant proteins will no longer be able to function as a metal binding RING domain, we constructed a series of additional constructs in which we mutated pairs of cysteines from each putative Zn binding site (Figure 5A) and a mutant in which we mutated 4 cysteines, two from each putative Zn binding site. As expected none of the CESA7 constructs were able to complement the *cesa7*^*irx3-7*^ mutant. In contrast, the multiple cysteine mutant constructs of CESA4 and CESA8 were still able to complement the mutant in a manner comparable to the single cysteine mutants even though we designed the mutations to ensure the sequence is no longer able to function as a metal bind RING domain (Figure 5B). Together, the data suggests a very different function for the RING domain between AtCESA7 and AtCESA4/8. This conclusion is supported by previous domain swap studies of chimeric proteins in which the RING domain of AtCESA7 was swapped with that of AtCESA4 or AtCESA8 and neither were able complement the *cesa7*^*irx3-7*^ mutant ^43^. While our data demonstrates that the cysteine residues in this region or CESA4 and CESA8 are not compatible with them forming a metal binding RING domain, the results are consistent with our previous data on the S-acylation of CESA proteins. We have previously demonstrated that when clusters of cysteines are modified by S-acylation, the essential nature of this modification is only revealed when all acylated cysteines in the region are mutated ^15^. To demonstrate the different patterns of S-acylation of CESA4 and CESA8 compared to CESA7 more directly, we examined levels of S-acylation in mutants in which known S-acylation sites are mutated (Figure 6). We have previously demonstrated that when all cysteines within the variable region 2 (4 cysteines) and C-terminal region (2 cysteines) of CESA7 are mutated, the level of CESA7 S-acylation was significantly reduced (Figure 6A) ^15^. In contrast when we mutated the cysteines in the same regions of CESA4 (8 cysteines) and CESA8 (5 cysteines), we observed little or no reduction in the level of protein S-acylation (Figure 6). This result is consistent with CESA4 and CESA8 still be extensively S-acylated at sites other than the VR2 and CT sites, while CESA7 is not. These other sites would include N-terminal cysteines previously considered part of the metal binding RING domain.

**Figure 6.**
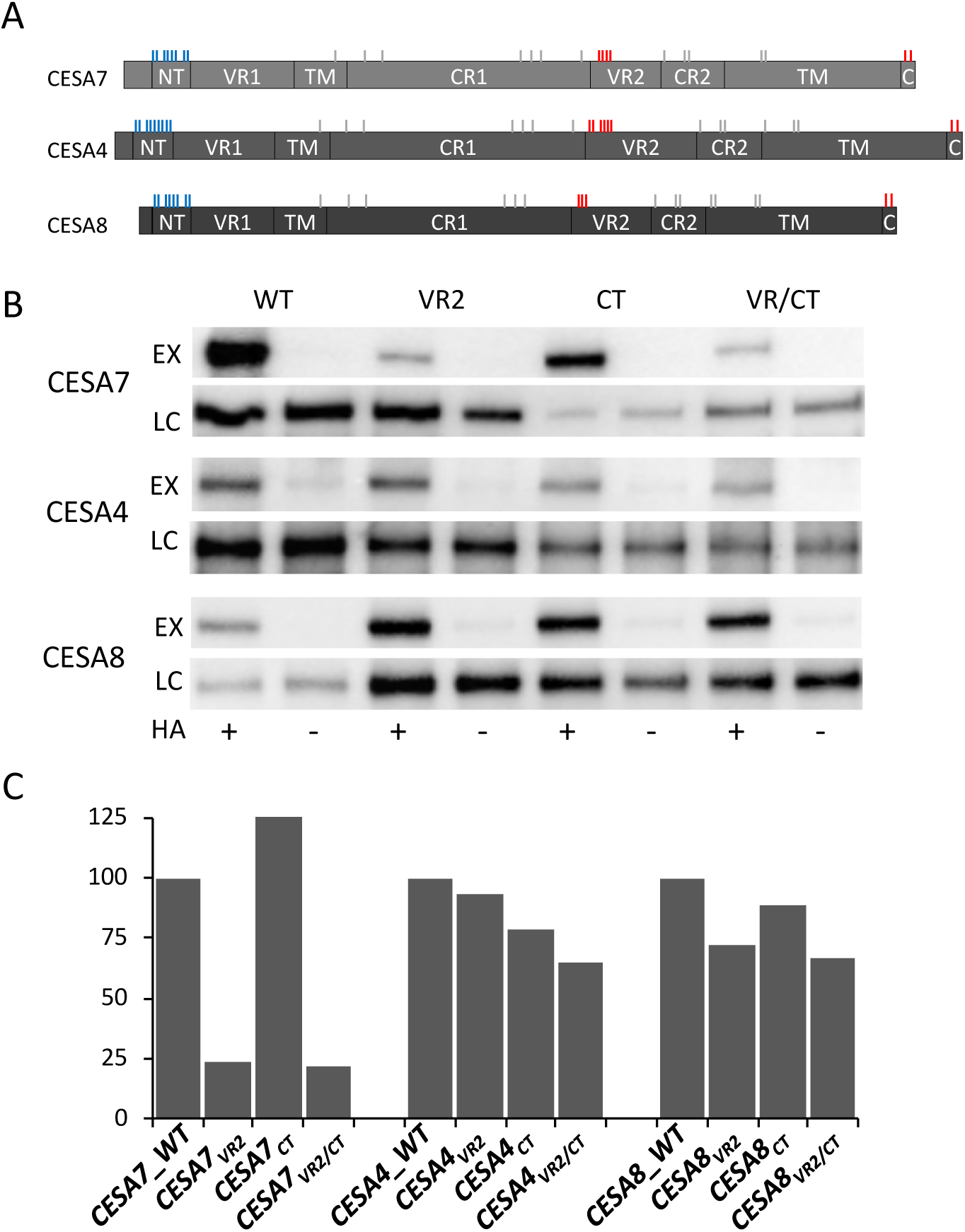
Differential S-acylation of CESA isoforms. (A) Diagrammatic representation of CESA protein structure indicating the position of cysteine residues (vertical bars). Cluster of mutated cysteines in the RING domain are coloured blue, while cysteines mutated in the variable regions 2 (VR2) and C-terminus (CT) are coloured red. The transmembrane (TM), conserved (CR1, CR2) and variable 1 (VR1) regions are also indicated. (B) EYFP tagged WT, VR2, CT or VR2/CT version of CESA7, CESA4 and CESA8 transformed into *cesa7*^*irx3-7*^, *cesa4*^*irx5-4*^ and *cesa8*^*irx1-7*^ respectively. Acyl-RAC assays were performed on mature stem material. For each assay, the experimental sample (EX) was compared with the loading control (LC) with or without (+/-) hydroxylamine (HA) for hydroxylamine-dependent capture of S-acylated proteins. Proteins were detected using protein specific antibodies. (C) Quantification of the immunoblots shown in B generated by normalising the signal relative to the loading control and expressed as percentage of WT.

To study the sequence variation within the RING domain, we aligned the RING domain sequences (46 amino acid long stretch) of all CESA proteins from 43 fully sequenced genomes and generated a consensus sequence for each CESA class (Figure 5C, Figure S7). The RING domain of CESA7 was most highly conserved; excluding the 8 conserved cysteine residues, 30 of the remaining 38 amino acids exhibited more than 90% conservation across species. In contrast, the consensus for CESA8 exhibited much greater variation with only 11 residues showing more than 90% conservation across the class, while the RING domain of CESA4 also appeared less well conserved than that of CESA7 with 16 invariant residues. We interpret this analysis as the higher conservation of CESA7 retaining its function as RING domain while the selective pressure on CESA4 and CESA8 functions largely to retain the cysteines only.

Based largely on studies performed following expression of the RING domain in yeast, it was suggested that the RING domain cysteines of a CESA from cotton act to reversibly dimerise CESA proteins by forming disulphide bonds ^44^. While our data do not support such models, they do suggest that in CESA4 and CESA8 this domain has diversified and is able to form structures other than a canonical zinc binding RING domain. RING-type zinc fingers are classified into a number of groups by PFAM. The RING domain of CESA proteins has been assigned the PFAM id zf-UDP (PF14569) and contains cysteines at all 8 metal ligand positions. Only two other RING domains contain cysteines at all 8 metal binding ligand positions: zf-RING_4 (PF1570), which is present in 8 proteins including 4 CSLD proteins, and FYVE (PF0136), composed of 13 proteins in Arabidopsis. Interestingly, the FYVE domain functions in lipid binding ^45^. It is plausible that the zf-UDP of CESA4 and CESA8 initially evolved as lipid binding domains and this function has been replaced by S-acylation as a means of binding the domain to a membrane.

### Concluding Remarks

We present a large-scale atlas of protein S-acylation across six tissues of Arabidopsis, thus enabling the broader study of this elusive modification and its biological roles in plants. Many of the proteins we identified in our acylome are predicted to be membrane localised even though they lack any transmembrane helices. S-acylation is likely to have an important, if not essential, role in anchoring these proteins to the membrane and consequently is important to the overall organisation of plant cells. We have demonstrated the robustness of the data by identifying many of the plant proteins that have been identified as S-acylated in previous studies. While we have focussed on proteins that fall into our high confidence group, this is likely to be biased in favour of more abundant proteins that are preferentially detected by MS. Lower abundance proteins are likely to have lower confidence scores. However, many of these proteins would be truly acylated and should be taken into consideration when examining one’s favourite group of proteins. Our high confidence group contains nearly 2600 proteins that represent around 9% of all Arabidopsis proteins. While this number initially appears high, it is comparable with the number of proteins identified in humans and mice. This suggests that S-acylation is a very important modification and likely to be as essential for the proper functioning of many plant proteins as it is in metazoans. However, it is important to add a note of caution that analysis of acylation sites either by indirect methods, such as the Acyl-RAC used here, or directly by the addition of labelled fatty acids, has the potential to generate false positives. The most stringent categorisation only includes proteins identified by both of these methods ^1^. Similarly, when a protein is identified as S-acylated, there is no measure of the relative amount of the modified vs unmodified form. Assays such as PEGylation provide a measure of the relative proportion of different protein forms, but are currently low throughput. PEGylation of KOR suggests that a large proportion of the protein is acylated (Figure 3A), but it may be much lower for other proteins and may also vary dramatically in different tissues and/or under different environmental conditions. We consider that first and foremost, the acylome data presented may be mined as a starting point for identifying how this modification affects protein function. We demonstrate the utility of the dataset by demonstrating the important differences in the structure and function of a conserved domain at the N-terminus of all CESA proteins that was previously assumed to perform similar function in various CESA classes.

Overall, our data indicates that S-acylation is far more widespread in plants than anticipated. Its role in cell wall biosynthesis and dynamics is likely to be even broader than previously shown. Novel functions in cellular traffic, signalling, defence and membrane-cytoskeleton interactions are very likely. We have shown how the identification of S-acylation sites in CESA proteins has forced a re-appraisal of structure-function relationships. Our acylome data, resolved to more than 5000 individual cysteine residues, makes it possible to address these and other important biological questions.

## Materials and methods

### Plant material

For proteomics, Arabidopsis Col0 seed was grown for 7 days on plates followed by a further 4 weeks on soil as previously described ^46^. The material collected was composed of inflorescence stem (after removing flowers, siliques and leaves and the basal 1cm), siliques (at various developmental stages), meristem (apical 1 cm of stem tip including unopened flowers) and rosette leaves. Seedlings were grown on plates for 7 days.

The loss of function mutants used in the study, *cesa8*^*irx1-7*^, *cesa7*^*irx3-7* 47^, *cesa4*^*irx5-4* 48^, *cc1, cc2, cc1 cc2* ^49^, *cmu1, cmu2, cmu1 cmu2* ^26^, *csi1* ^50^ and *kor1-1* ^51^ have been previously described. *tub6-2* (SALKseq_126440) and *tub8-1* (SALK_116231) were obtained from NASC and homozygous plants were isolated and confirmed by PCR based genotyping and crossed to create *tub6 tub8* double mutant. All plant transformation and complementation analysis was as performed as described previously. Cellulose content was determined using a medium throughput adaptation of Updegraff’s method ^47^. For KOR1 lines, complementation assay was performed using plate based assays.

Transient expression in tobacco has been described in detail previously ^52^. Agrobacterium solutions were infiltrated into lower side of 4 weeks old *Nicotiana benthamiana* plants using a 1-ml syringe. The infiltrated leaves were harvested 4 days after infiltration, frozen in liquid nitrogen and stored at -80°C until further use.

### Acyl-RAC assay, trypsin digestion and mass spectrometry

The optimised ACYL-RAC assay was based on previously described methods ^17,53^. About 800 mg of frozen tissue powder was solubilised in 20 mL of lysis buffer (100mM Tris pH 7.2, 150mM NaCl, 5mM EDTA, 2.5% SDS, 1 mM PMSF and Sigma plant protease inhibitor cocktail) and heated at 70°C for 20 minutes in a water bath. Samples were then centrifuged at 4500g and supernatant filtered through miracloth. Protein concentration was measured using Millipore BCA assay kit according to manufacturer’s instructions and typically revealed a protein concentration of 1-2 mg/mL.

The extract was incubated with 300 µl of Immobilized TCEP Disulfide Reducing Gel (Thermo Scientific, Catalog number 77712) for 1 hour and the TCEP gel was then removed by passing the sample through a plastic column fitted with a porous disc (Thermo Scientific, Cat number 89898). NEM was then added to a final concentration 20 mM followed by incubation at room temperature in the dark for 2 hours. Free NEM was then removed by adding 100 mM 2-3 dimethyl-1-3 butadiene (DMB) and incubating for 1 hour ^53^. DMB was removed by extraction twice with 0.1 volume of chloroform.

9 mL of blocked protein sample was combined with 4.5 mL of either 1M hydroxylamine (final concentration 333 mM) or water and 80 mg (dry weight equivalent) thiopropyl sepharose beads (GE Healthcare Catalog number 17042001) and incubated for 1 hour at room temperature in a column fitted with a porous disc. Samples were washed for 5min in 5ml: 8M urea (2x), 2M NaCl (2x), 80% acetonitrile + 0.1% TFA (2x) and 50 mM ammonium bicarbonate (AmBic) (3x) and the beads dried completely by centrifugation at 300g for 2 minutes. The columns were capped at the bottom and 200 µl AmBic containing 4 ng/µl of Trypsin + LysC (Promega Catalog number V5073) was added to each sample and incubated in a 37°C shaker for 16 hours. After digestion, unbound peptides were collected by centrifugation at 300g for 2 minutes and the beads were then washed again as described above. Bound peptides were then eluted by adding 200 µl AmBic containing 50 mM DTT and incubating in a 37°C shaker for 30 minutes followed by centrifugation at 300g for 2 minutes. Peptides were alkylated by adding iodoacetamide (IAM) to a final concentration of 100 mM and incubating in dark at room temperature with gentle agitation for 2 hours. Further DTT was added to bring final concentration to 100 mM to quench the unreacted IAM and samples stored at -20°C until further use.

Reduced and alkylated peptides were acidified by adding equal volume of 0.1% formic acid. Acidified samples were bound to OLIGO R3 reverse phase resin (Thermo Scientific, catalogue number 1133903), washed twice with 0.1% formic acid and eluted with 200 µl of 30% acetonitrile + 0.1% formic acid. Peptides were then quantified using Pierce Quantitative Fluorometric Peptide Assay (Thermo Scientific, catalogue number 23290) according to manufacturer’s instructions. Known quantities (500 – 3000 ng) of peptides were then aliquoted into mass spec vials and dried completely using a speed vac.

Digested samples were analysed by LC-MS/MS using an UltiMate® 3000 Rapid Separation LC (RSLC, Dionex Corporation, Sunnyvale, CA) coupled to a Q Exactive HF (Thermo Fisher Scientific, Waltham, MA) mass spectrometer. Peptide mixtures were separated using a multistep gradient from 95% A (0.1% FA in water) and 5% B (0.1% FA in acetonitrile) to 7% B at 1 min, 18% B at 35 min, 27% B in 43 min and 60% B at 44 min at 300 nL min-1, using a 75 mm x 250 μm i.d. 1.7 µM CSH C18, analytical column (Waters). Peptides were selected for fragmentation automatically by data dependant analysis.

### Database searching

Database searching and PSM (peptide spectrum matches) processing was performed in Proteome Discoverer 2.2 (Thermo Fisher Scientific, UK). Two complementary search engines Sequest™ HT (Thermo Fisher Scientific, UK) and Mascot (Matrix Science, UK) were used for searching the data. These engines were set up to search the Arabidopsis thaliana Araport11 protein database (48359 entries) assuming the digestion enzyme trypsin. A Precursor Mass Tolerance of 10 ppm and Fragment Mass Tolerance of 0.02 Da was used for Sequest while Mascot was searched with a Precursor Mass Tolerance of 20 ppm and Fragment Mass Tolerance of 0.02 Da. Oxidation of methionine, carbamidomethyl of cysteine and N-ethylmaleimide of cysteine were specified as variable modifications for both search engines. All other program settings were kept as default. Modifications and precursor ion intensity were exported out of Proteome Discoverer and majority of further calculations and data processing was performed in Microsoft Excel.

### Peptide and Protein confidence scores and classes

The data was first normalised using assay total signal normalisation. Each biological replicate constitutes one assay and includes 2 sample types, HA-plus and HA-minus. So there are 18 assays in the data, 3 for each tissue. Average assay total intensity across 18 assays was 1.4E10. For simplicity, the intensities were normalised for total signal intensity for each assay to be 1E10. The normalisation factor thus calculated was used to compute normalised intensity for each PSM.

The data was then reduced to peptide forms (defined as a combination of peptide sequence and modification) and intensities were computed for each peptide form by taking the sum of intensities of all PSMs matching to a particular peptide form. Three modifications were used in the searches: oxidation of methionine, carbamidomethyl of cysteine and n-ethylmaleimide of cysteine. At this stage, PSM intensities for all peptide forms were summed together leading to a total of 11188 peptide forms.

In order to get meaningful fold change (FC) ratios, missing values were replaced with 2 (a small number relative to average peptide intensity of ∼590000) and data was log2 transformed. T-test P values and fold changes were then calculated on the transformed data.

The peptides sequences were then mapped to Araport11 proteome. When multiple splice variants matched to a peptide, only one splice variant (with lowest isoform number) was kept for each locus. For each protein, total intensity was computed by taking sum of peptides matching to that protein and computed for exclusive and ambiguous peptides separately. For proteins where an exclusive peptide was available, its protein score was based only on exclusive peptide intensities. Ambiguous peptides contributed to protein score calculation only when exclusive peptides were not present.

Next, a protein class was computed by comparing the protein score and the peptide score of the best peptide matching to that protein. The lower of those two scores was considered to be the protein class.

### DNA Cloning

Gateway technology (ThermoFisher Scientific, UK) was used for construction of most of the plasmids. Cloning of wild type CESA8, 7 and 4 CDS fragments in a Gateway entry vector, pDONOR/pZEO, has been described previously ^43^. All other CDS fragments including KOR, CC1, CC2, CMU1, CMU2, CMU3, CSI1, TUA1, TUA4, TUB6, TUB8, SHOU4 and SHOU4L were amplified using primers listed in Table S6 and entry clones were made using the same procedure. The vector for expressing CESA7 has been described previously. New destination vectors were constructed using pCambia1300 backbone. The component fragments of these vectors (promoters, tags, Gateway cassette and the terminator) were PCR amplified and cloned into a pJET vector using a CloneJET PCR Cloning Kit (ThermoFisher Scientific, UK). All primers (Table S6) included appropriate restriction sites to allow concatenation of the fragments. The inserts in the pJET vector were fully sequenced before assembling to create the final destination vectors. Cysteine mutants of CESA4, CESA7, CESA8 and KOR1 were made using PCR-based site directed mutagenesis ^15,54^. Sequences of different GSF Primers used are listed in Table S6. Plasmids were sequenced fully to verify that no mutations had occurred.

### Acyl-RAC immunoblots and Acyl PEG Exchange Assay

The Acyl-RAC assay protocol was the same as described above in the proteomics section. After washing the TPS beads with the wash buffer, instead of trypsin digestion, proteins were eluted with DTT. The proteins were separated on 7.5% SDS-PAGE gels, blotted onto PVDF membranes and probed with various antibodies. Primary antibodies used in this study included: anti KOR1 ^55^, Mouse anti HSP73 (Enzo life sciences, Cat# ADI-SPA-818, used at 1: 20000 dilution), anti-Laccase (peptide antibody raised in rabbit against Apple laccase MDP0000242301; peptide sequence RDIHSEKWKGPRSTN, used at 1: 500 dilution) and Mouse anti GFP (Invitrogen, Cat# 14-6674-82, used at 1: 1000 dilution). The secondary antibody used were anti-Rabbit HRP (Dako, Cat# 0448, used at 1:1000 dilution) and anti-Mouse HRP (Dako, Cat# 0448, used at 1:1000 dilution).

The acyl PEG exchange (APE) assay was performed using a modification of previously described protocol ^31^. The initial part of the assay including crude protein extraction, disulphide bond reduction, NEM alkylation and NEM removal was the same as described above for the Acyl-RAC assay. After NEM treatment and NEM removal, the protein sample was divided into 2 portions. HA (final concentration 333 mM) was added to the HA+ sample while water was added to the HA-sample. After incubating the samples for 1 hour with gentle rotation, the protein samples were precipitated using chloroform methanol precipitation method. After washing the pellet with methanol, the pellet was dried and resuspended in 1 mL of 100 mM Tris HCl, pH7.4, 4 mM EDTA, 4 % SDS. mPEG maleimide 5 kD (Sigma) was then added to a final concentration of 1 mM and samples incubated at room temperature with gentle agitation. The samples were then precipitated again and resuspended in 100 µl of 2x Laemmli buffer without DTT and analysed on non-reducing SDS-PAGE followed by botting and probing with anti-KOR antibody.

## Acknowledgements

This work as supported by a grant from the BBSRC reference BB/P01013X/1. We acknowledge help provided by Stacey Warwood, Julian Selley, Ronan O’cualain, David Knight and Emma-jayne Keevill at the Bio-MS core facility, University of Manchester for the proteomic analysis. We would like to thank Staffan Persson for kindly providing *cmu1 cmu2, cc1 cc2* and csi1 mutants, Samantha Vernhettes for providing *kor1-1* mutants and Thomas Nuhse for his comments on the manuscript.

## Supplementary data

Supplementary data contains supplementary figures (Figure 1-7), Supplementary data file (Table S1) and supplementary tables (Table 2-6).

