## Supplementary data for "An atlas of Arabidopsis protein S-Acylation reveals its widespread role in plant cell organisation of and function"

This file contains supplementary figures 1 to 7 and supplementary tables 2 to 6.

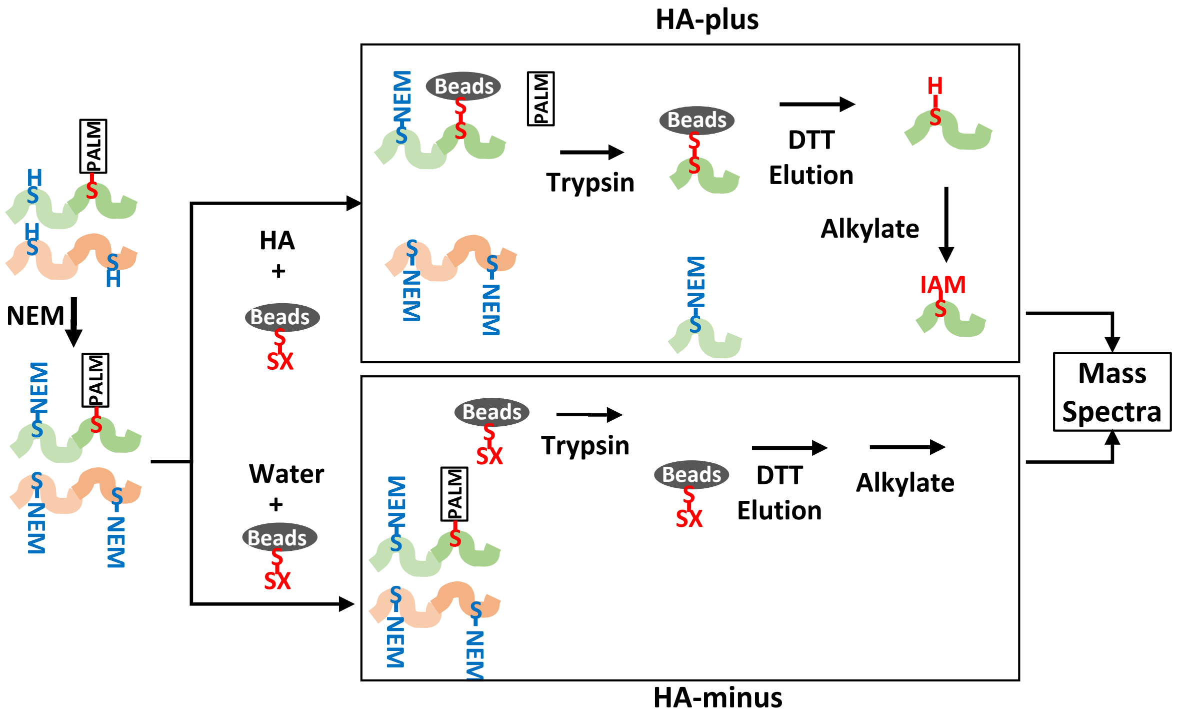

**Figure S1. Overview of the Acyl-RAC and on-bead digestion protocol.**
Extracted proteins were treated with TCEP to reduce the disulphide bonded cysteines. Free cysteines were then capped with NEM and proteins were captured by binding to thiopropyl sepharose beads in the presence or absence of hydroxylamine (HA). Following on-bead trypsin digestion bound peptides that contain acyl cysteines were eluted with DTT, alkylated with iodoacemtamide, and analysed by mass spectrometry.

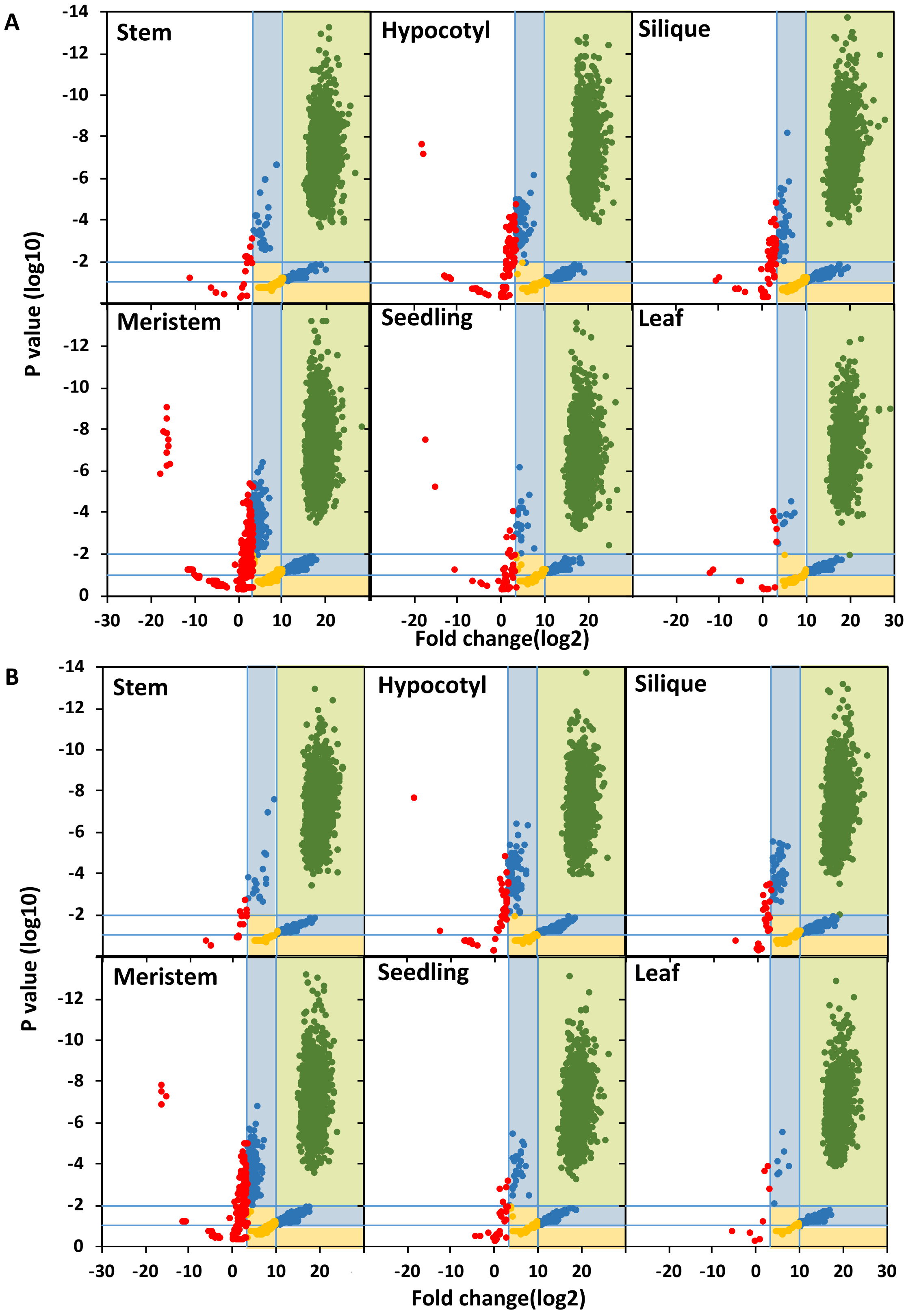

**Figure S2. Confidence score assignment**

Fold change (log2) and P value (log10) derived from peptide (A) and protein (B) intensities of each tissue treated with HA compared to untreated controls. The shaded areas of the plot indicate the way in which the peptides were classified into different groups with 1 (light green shading), 2 (light blue shading) and 3 (light yellow shading) referring to the high, medium and low confidence groups. Peptides in the unshaded area (red dots) were not considered further (see text for details). For each tissue the total number of peptides and proteins falling within each group is shown in Table 1.

A

```
                   10        20        30        40        50        60        70        80        90
AtROP01 ------------MSASRFVKCVTVGDGAVGKTCLLISYTSNTFPTDYVPTVFDNFSANVVVNGSTVNLGLWDTAGQEDYNRLRPLSYRGADVFILAFSL  87
AtROP02 -------------MASRFIKCVTVGDGAVGKTCMLISYTSNTFPTDYVPTVFDNFSANVVVDGNTVNLGLWDTAGQEDYNRLRPLSYRGADVFILAFSL  86
AtROP03 ------------MSASRFIKCVTVGDGAVGKTCLLISYTSNTFPTDYVPTVFDNFSANVVVNGATVNLGLWDTAGQEDYNRLRPLSYRGADVFILAFSL  87
AtROP04 ------------MSASRFIKCVTVGDGAVGKTCMLISYTSNTFPTDYVPTVFDNFSANVVVDGNTVNLGLWDTAGQEDYNRLRPLSYRGADVFILAFSL  87
AtROP05 ------------MSASRFIKCVTVGDGAVGKTCLLISYTSNTFPTDYVPTVFDNFSANVVVNGATVNLGLWDTAGQEDYNRLRPLSYRGADVFILAFSL  87
AtROP06 ------------MSASRFIKCVTVGDGAVGKTCLLISYTSNTFPTDYVPTVFDNFSANVIVDGNTINLGLWDTAGQEDYNRLRPLSYRGADVFLLAFSL  87
AtROP07 ------------MSTARFIKCVTVGDGAVGKTCMLISYTSNTFPTDYVPTVFDNFSANVVVDGSTVNLGLWDTAGQEDYNRLRPLSYRGADVFLLAFSL  87
AtROP08 MSASMAATSTSSATATTFIKCVTVGDGAVGKTCLLISYTSNTFPTDYVPTVFDNFNANVLVDGKTVNLGLWDTAGQEDYNRVRPLSYRGADVFILAFSL  99
AtROP09 ------------MSASKFIKCVTVGDGAVGKTCMLICYTSNKFPTDYIPTVFDNFSANVAVDGQIVNLGLWDTAGQEDYSRLRPLSYRGADIFVLAFSL  87
AtROP10 ----------MASSASKFIKCVTVGDGAVGKTCMLICYTSNKFPTDYIPTVFDNFSVNVVVEGITVNLGLWDTAGQEDYNRLRPLSYRGADVFVLAFSL  89
AtROP11 ----------MASSASKFIKCVTVGDGAVGKTCMLICYTSNKFPTDYIPTVFDNFSANVVVEGTTVNLGLWDTAGQEDYNRLRPLSYRGADVFVLSFSL  89

        100       110       120       130       140       150       160       170       180       190
AtROP01 ISKASYENVSKKWIPELKHYAPGVPIVLVGTKLDLRDDKQFFIDHPGAVPITTAQGEELRKQIGAPTYIECSSKTQENVKAVFDAAIRVVLQPPKQ--- 183
AtROP02 ISKASYENIAKKWIPELRHYAPGVPIILVGTKLDLRDDKQFFIDHPGAVPITTNQGEELKKLIGSAVYIECSSKTQQNVKAVFDAAIKVVLQPPKQ--- 182
AtROP03 ISKASYENVSKKWIPELKHYAPGVPIVLVGTKLDLRDDKQFFIDHPGAVPITTAQGEELKKLIGAPAYIECSSKTQENVKGVFDAAIRVVLQPPKQ--- 183
AtROP04 ISKASYENVAKKWIPELRHYAPGVPIILVGTKLDLRDDKQFFIDHPGAVPITTNQGEELKKLIGSPIYIECSSKTQQNVKAVFDAAIKVVLQPPKQ--- 183
AtROP05 ISKASYENVSKKWIPELKHYAPGVPIVLVGTKLDLRDDKQFFIDHPGAVPITTVQGEELKKLIGAPAYIECSSKSQENVKGVFDAAIRVVLQPPKQ--- 183
AtROP06 VSKASYENVSKKWVPELRHYAPGVPIILVGTKLDLRDDKQFFAEHPGAVPISTAQGEELKKLIGAPAYIECSAKTQQNVKAVFDAAIKVVLQPPKNK-- 184
AtROP07 ISKASYENIHKKWLPELKHYAPGIPIVLVGTKLDLRDDKQFLKDHPGAASITTAQGEELRKMIGAVRYLECSSKTQQNVKAVFDTAIRVALRPPKA--- 183
AtROP08 ISRPSFENIAKKWVPELRHYAPTVPIVLVGTKSDLRDNMQFPKNYPGACTIFPEQGQELRKEIGALAYIECSSKAQMNVKAVFDEAIKVVLHPPSK--- 195
AtROP09 ISKASYENVLKKWMPELRRFAPNVPIVLVGTKLDLRDDKGYLADHTNV--ITSTQGEELRKQIGAAAYIECSSKTQQNVKAVFDTAIKVVLQPPRRKEV 184
AtROP10 ISRASYENVFKKWIPELQHFAPGVPIVLVGTKMDLREDRHYLSDHPGLSPVTTSQGEELRKHIGATYYIECSSKTQQNVKAVFDAAIKVVIKPAVKQKE 188
AtROP11 VSRASYENVFKKWIPELQHFAPGVPLVLVGTKLDLREDKHYLADHPGLSPVTTAQGEELRKLIGATYYIECSSKTQQNVKAVFDSAIKEVIKPLVKQKE 188

         200          210          220
AtROP01 ---KKK----KSKAQKACSIL---------- 197
AtROP02 ---KKK----K-KNKNRCAFL---------- 195
AtROP03 ---KKK----KSKAQKACSIL---------- 197
AtROP04 ---KKK----K-KNKNRCVFL---------- 196
AtROP05 ---KKK----KNKAQKACSIL---------- 197
AtROP06 ---KKK----KRKSQKGCSIL---------- 198
AtROP07 ---KKKIKPLKTKRSRICFFL---------- 201
AtROP08 ---TKK----RKRKIGLCHVL---------- 209
AtROP09 P--RRR----KNHRRSGCSIASIVCGGCTAA 209
AtROP10 ---KKK----KQKPRSGCLSNILCGKN---- 208
AtROP11 KTKKKK----KQKSNHGCLSNVLCGRIVTRH 215
```

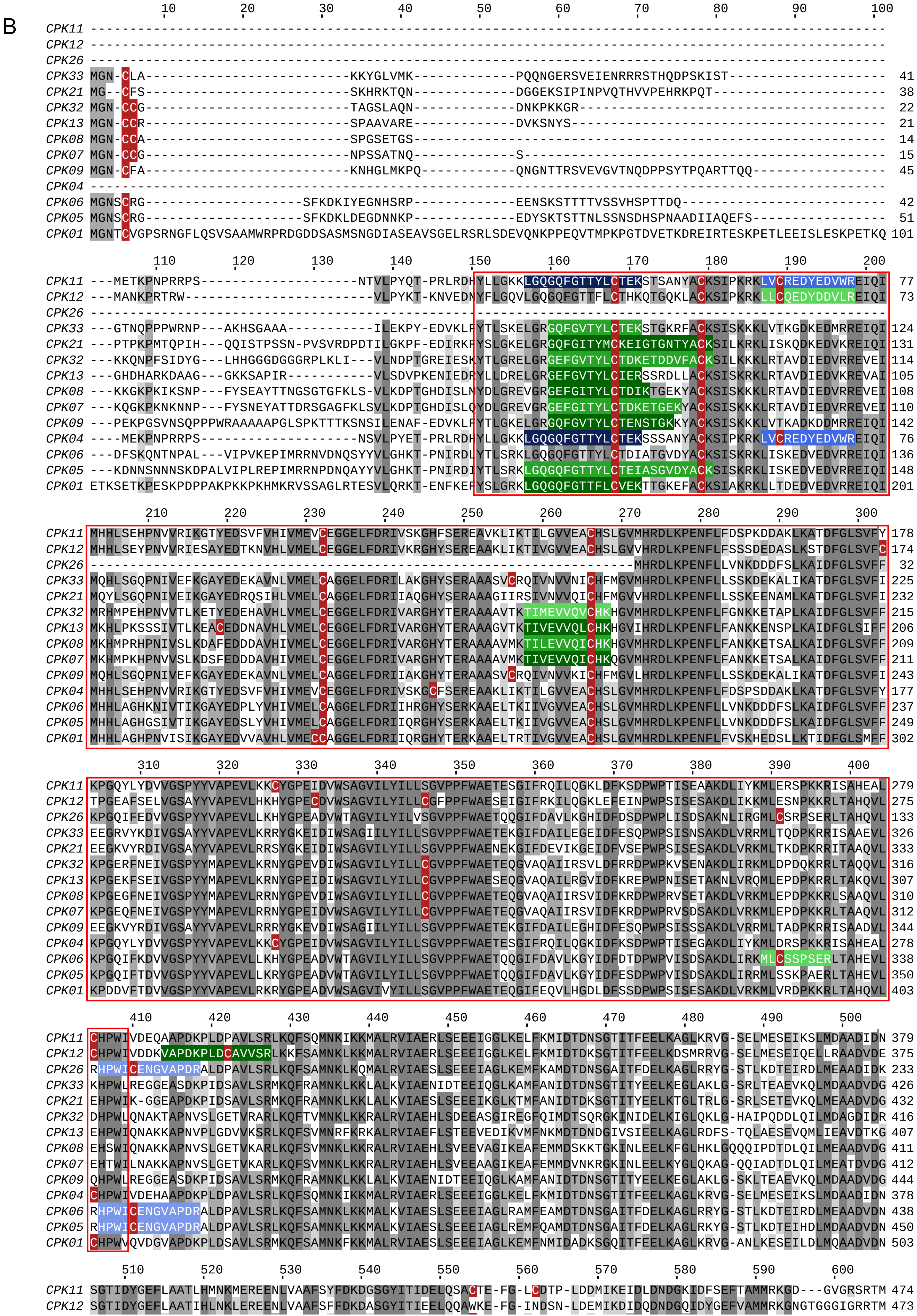
B
10 20 30 40 50 60 70 80 90 100
CPK11 ----------------------------------------------------------------------------------------------------
CPK12 ----------------------------------------------------------------------------------------------------
CPK26 ----------------------------------------------------------------------------------------------------
CPK33 MGN-CLA-------------------------KKYGLVMK-------------PQQNGERSVEIENRRRSTHQDPSKIST-------------------- 41
CPK21 MG--CFS-------------------------SKHRKTQN-------------DGGEKSIPINPVQTHVVPEHRKPQT---------------------- 38
CPK32 MGN-CCG-------------------------TAGSLAQN-------------DNKPKKGR--------------------------------------- 22
CPK13 MGN-CCR-------------------------SPAAVARE-------------DVKSNYS---------------------------------------- 21
CPK08 MGN-CCA-------------------------SPGSETGS------------------------------------------------------------ 14
CPK07 MGN-CCG-------------------------NPSSATNQ-------------S---------------------------------------------- 15
CPK09 MGN-CFA-------------------------KNHGLMKPQ------------QNGNTTRSVEVGVTNQDPPSYTPQARTTQQ----------------- 45
CPK04 ----------------------------------------------------------------------------------------------------
CPK06 MGNSCRG-------------------SFKDKIYEGNHSRP-------------EENSKSTTTTVSSVHSPTTDQ-------------------------- 42
CPK05 MGNSCRG-------------------SFKDKLDEGDNNKP-------------EDYSKTSTTNLSSNSDHSPNAADIIAQEFS----------------- 51
CPK01 MGNTCVGPSRNGFLQSVSAAMWRPRDGDDSASMSNGDIASEAVSGELRSRLSDEVQNKPPEQVTMPKPGTDVETKDREIRTESKPETLEEISLESKPETKQ 101
110 120 130 140 150 160 170 180 190 200
CPK11 ---METKPNPRRPS-------------------NTVLPYQT-PRLRDHYLLGKKLGQGQFGTTYLCTEKSTSANYACKSIPKRKLVCREDYEDVWREIQI 77
CPK12 ---MANKPRTRW-----------------------VLPYKT-KNVEDNYFLGQVLGQGQFGTTFLCTHKQTGQKLACKSIPKRKLLCQEDYDDVLREIQI 73
CPK26 ----------------------------------------------------------------------------------------------------
CPK33 ---GTNQPPPWRNP---AKHSGAAA----------ILEKPY-EDVKLFYTLSKELGRGQFGVTYLCTEKSTGKFRACKSISKKKLVTKGDKEDMRREIQI 124
CPK21 ---PTPKPMTQPIH---QQISTPSSN-PVSVRDPDTILGKPF-EDIRKFYSLGKELGRGQFGITYMCKEIGTGNTYACKSILKRKLISKQDKEDVKREIQI 131
CPK32 ---KKQNPFSIDYG---LHHGGGDGGGRPLKLI---VLNDPTGREIESKYTLGRELGRGEFGVTYLCTDKETDDVFACKSILKKKLRTAVDIEDVRREVEI 114
CPK13 ---GHDHARKDAAG---GKKSAPIR----------VLSDVPKENIEDRYLLDRELGRGEFGVTYLCIERSSRDLLACKSISKRKLRTAVDIEDVKREVAI 105
CPK08 ---KKGKPKIKSNP---FYSEAYTTNGSGTGFKLS-VLKDPTGHDISLMYDLGREVGRGEFGITYLCTDIKTGEKYACKSISKKKLRTAVDIEDVRREVEI 108
CPK07 ---KQGKPKNKNNP---FYSNEYATTDRSGAGFKLSVLKDPTGHDISLQYDLGREVGRGEFGITYLCTDKETGEKYACKSISKKKLRTAVDIEDVRREVEI 110
CPK09 ---PEKPGSVNSQPPPWRAAAAAPGLSPKTTTKSNSILENAF-EDVKLFYTLGKELGRGQFGVTYLCTENSTGKKYACKSISKKKLVTKADKDDMRREIQI 142
CPK04 ----MEKPNPRRPS-------------------NSVLPYET-PRLRDHYLLGKKLGQGQFGTTYLCTEKSSSANYACKSIPKRKLVCREDYEDVWREIQI 76
CPK06 ---DFSKQNTNPAL---VIPVKEPIMRRNVDNQSYYVLGHKT-PNIRDLYTLSRKLGQGQFGTTYLCTDIATGVDYACKSISKRKLISKEDVEDVRREIQI 136
CPK05 ---KDNNSNNNSKDPALVIPLREPIMRRNPDNQAYYVLGHKT-PNIRDIYTLSRKLGQGQFGTTYLCTEIASGVDYACKSISKRKLISKEDVEDVRREIQI 148
CPK01 ETKSETKPESKPDPPAKPKKPKHMKRVSSAGLRTESVLQRKT-ENFKEFYSLGRKLGQGQFGTTFLCVEKTTGKEFACKSIAKRKLLTDEDVEDVRREIQI 201
210 220 230 240 250 260 270 280 290 300
CPK11 MHHLSEHPNVVRIKGTYEDSVFVHIVMEVCEGGELFDRIVSKGHFSEREAVKLIKTILGVVEACHSLGVMHRDLKPENFLFDSPKDDAKLKATDFGLSVFY 178
CPK12 MHHLSEYPNVVRIESAYEDTKNVHLVMELCEGGELFDRIVKRGHYSEREAAKLIKTIVGVVEACHSLGVVHRDLKPENFLFSSSDEDASLKSTDFGLSVFC 174
CPK26 --------------------------------------------------------------------MHRDLKPENFLLVNKDDDFSLKAIDFGLSVFF 32
CPK33 MQHLSGQPNIVEFKGAYEDEKAVNLVMELCAGGELFDRILAKGHYSERAAASVCRQIVNVVNICHFMGVMHRDLKPENFLLSSKDEKALIKATDFGLSVFI 225
CPK21 MQYLSGQPNIVEIKGAYEDRQSIHLVMELCAGGELFDRIIAQGHYSERAAAGIIRSIVNVVQICHFMGVVHRDLKPENFLLSSKEENAMLKATDFGLSVFI 232
CPK32 MRHMPEHPNVVTLKETYEDEHAVHLVMELCEGGELFDRIVARGHYTERAAAAVTKTIMEVVQVCHKHGVMHRDLKPENFLFGNKKETAPLKAIDFGLSVFF 215
CPK13 MKHLPKSSSIVTLKEACEDDNAVHLVMELCEGGELFDRIVARGHYTERAAAGVTKTIVEVVQLCHKHGVIHRDLKPENFLFANKKENSPLKAIDFGLSIFF 206
CPK08 MKHMPRHPNIVSLKDAFEDDDAVHIVMELCEGGELFDRIVARGHYTERAAAAVMKTILEVVQICHKHGVMHRDLKPENFLFANKKETSALKAIDFGLSVFF 209
CPK07 MKHMPKHPNVVSLKDSFEDDDAVHIVMELCEGGELFDRIVARGHYTERAAAAVMKTIVEVVQICHKQGVMHRDLKPENFLFANKKETSALKAIDFGLSVFF 211
CPK09 MQHLSGQPNIVEFKGAYEDEKAVNLVMELCAGGELFDRIIAKGHYTERAAASVCRQIVNVVKICHFMGVLHRDLKPENFLLSSKDEKALIKATDFGLSVFI 243
CPK04 MHHLSEHPNVVRIKGTYEDSVFVHIVMEVCEGGELFDRIVSKGCFSEREAAKLIKTILGVVEACHSLGVMHRDLKPENFLFDSPSDDAKLKATDFGLSVFY 177
CPK06 MHHLAGHKNIVTIKGAYEDPLYVHIVMELCAGGELFDRIIHRGHYSERKAAELTKIIVGVVEACHSLGVMHRDLKPENFLLVNKDDDFSLKAIDFGLSVFF 237
CPK05 MHHLAGHGSIVTIKGAYEDSLYVHIVMELCAGGELFDRIIQRGHYSERKAAELTKIIVGVVEACHSLGVMHRDLKPENFLLVNKDDDFSLKAIDFGLSVFF 249
CPK01 MHHLAGHPNVISIKGAYEDVVAVHLVMECCAGGELFDRIIQRGHYTERKAAELTRTIVGVVEACHSLGVMHRDLKPENFLFVSKHEDSLLKTIDFGLSMFF 302
310 320 330 340 350 360 370 380 390 400
CPK11 KPGQYLYDVVGSPYYVAPEVLKKCYGPEIDVWSAGVILYILLSGVPPFWAETESGIFRQILQGKLDFKSDPWPTISEAAKDLIYKMLERSPKKRISAHEAL 279
CPK12 TPGEAFSELVGSAYYVAPEVLHKHYGPECDVWSAGVILYILLCGFPPFWAESEIGIFRKILQGKLEFEINPWPSISESAKDLIKKMLESNPKKRLTAHQVL 275
CPK26 KPGQIFEDVVGSPYYVAPEVLLKHYGPEADVWTAGVILYILVSGVPPFWAETQQGIFDAVLKGHIDFDSDPWPLISDSAKNLIRGMLCSRPSERLTAHQVL 133
CPK33 EEGRVYKDIVGSAYYVAPEVLRRRYGKEIDIWSAGIILYILLSGVPPFWAETEKGIFDAILEGEIDFESQPWPSISNSAKDLVRRMLTQDPKRRISAAEVL 326
CPK21 EEGKVYRDIVGSAYYVAPEVLRRSYGKEIDIWSAGVILYILLSGVPPFWAENEKGIFDEVIKGEIDFVSEPWPSISESAKDLVRKMLTKDPKRRITAAQVL 333
CPK32 KPGERFNEIVGSPYYMAPEVLKRNYGPEVDIWSAGVILYILLCGVPPFWAETEQGVAQAIIRSVLDFRRDPWPKVSENAKDLIRKMLDPDQKRRLTAQQVL 316
CPK13 KPGEKFSEIVGSPYYMAPEVLKRNYGPEIDIWSAGVILYILLCGVPPFWAESEQGVAQAILRGVIDFKREPWPNISETAKNLVRQMLEPDPKRRLTAKQVL 307
CPK08 KPGEGFNEIVGSPYYMAPEVLRRNYGPEVDIWSAGVILYILLCGVPPFWAETEQGVAQAIIRSVIDFKRDPWPRVSETAKDLVRKMLEPDPKKRLSAAQVL 310
CPK07 KPGEQFNEIVGSPYYMAPEVLRRNYGPEIDVWSAGVILYILLCGVPPFWAETEQGVAQAIIRSVIDFKRDPWPRVSDSAKDLVRKMLEPDPKKRLTAAQVL 312
CPK09 EEGKVYRDIVGSAYYVAPEVLRRRYGKEVDIWSAGIILYILLSGVPPFWAETEKGIFDAILEGHIDFESQPWPSISSSAKDLVRRMLTADPKRRISAADVL 344
CPK04 KPGQYLYDVVGSPYYVAPEVLKKCYGPEIDVWSAGVILYILLSGVPPFWAETESGIFRQILQGKIDFKSDPWPTISEGAKDLIYKMLDRSPKKRISAHEAL 278
CPK06 KPGQIFKDVVGSPYYVAPEVLLKHYGPEADVWTAGVILYILLSGVPPFWAETQQGIFDAVLKGYIDFDTDPWPVISDSAKDLIRKMLCSSPSERLTAHEVL 338
CPK05 KPGQIFTDVVGSPYYVAPEVLLKRYGPEADVWTAGVILYILLSGVPPFWAETQQGIFDAVLKGYIDFESDPWPVISDSAKDLIRRMLSSKPAERLTAHEVL 350
CPK01 KPDDVFTDVVGSPYYVAPEVLRKRYGPEADVWSAGVIVYILLSGVPPFWAETEQGIFEQVLHGDLDFSSDPWPSISESAKDLVRKMLVRDPKKRLTAHQVL 403
410 420 430 440 450 460 470 480 490 500
CPK11 CHPWIVDEQAAPDKPLDPAVLSRLKQFSQMNKIKKMALRVIAERLSEEEIGGLKELFKMIDTDNSGTITFEELKAGLKRVG-SELMESEIKSLMDAADIDN 379
CPK12 CHPWIVDDKVAPDKPLDCAVVSRLKKFSAMNKLKKMALRVIAERLSEEEIGGLKELFKMIDTDKSGTITFEELKDSMRRVG-SELMESEIQELLRAADVDE 375
CPK26 RHPWICENGVAPDRALDPAVLSRLKQFSAMNKLKQMALRVIAESLSEEEIAGLKEMFKAMDTDNSGAITFDELKAGLRRYG-STLKDTEIRDLMEAADIDK 233
CPK33 KHPWLREGGEASDKPIDSAVLSRMKQFRAMNKLKKLALKVIAENIDTEEIQGLKAMFANIDTDNSGTITYEELKEGLAKLG-SRLTEAEVKQLMDAADVDG 426
CPK21 EHPWIK-GGEAPDKPIDSAVLSRMKQFRAMNKLKKLALKVIAESLSEEEIKGLKTMFANIDTDKSGTITYEELKTGLTRLG-SRLSETEVKQLMEAADVDG 432
CPK32 DHPWLQNAKTAPNVSLGETVRARLKQFTVMNKLKKRALRVIAEHLSDEEASGIREGFQIMDTSQRGKINIDELKIGLQKLG-HAIPQDDLQILMDAGDIDR 416
CPK13 EHPWIQNAKKAPNVPLGDVVKSRLKQFSVMNRFKRKALRVIAEFLSTEEVEDIKVMFNKMDTDNDGIVSIEELKAGLRDFS-TQLAESEVQMLIEAVDTKG 407
CPK08 EHSWIQNAKKAPNVSLGETVKARLKQFSVMNKLKKRALRVIAEHLSVEEVAGIKEAFEMMDSKKTGKINLEELKFGLHKLGQQQIPDTDLQILMEAADVDG 411
CPK07 EHTWILNAKKAPNVSLGETVKARLKQFSVMNKLKKRALRVIAEHLSVEEAAGIKEAFEMMDVNKRGKINLEELKYGLQKAG-QQIADTDLQILMEATDVDG 412
CPK09 QHPWLREGGEASDKPIDSAVLSRMKQFRAMNKLKKLALKVIAENIDTEEIQGLKAMFANIDTDNSGTITYEELKEGLAKLG-SKLTEAEVKQLMDAADVDG 444
CPK04 CHPWIVDEHAAPDKPLDPAVLSRLKQFSQMNKIKKMALRVIAERLSEEEIGGLKELFKMIDTDNSGTITFEELKAGLKRVG-SELMESEIKSLMDAADIDN 378
CPK06 RHPWICENGVAPDRALDPAVLSRLKQFSAMNKLKKMALKVIAESLSEEEIAGLRAMFEAMDTDNSGAITFDELKAGLRRYG-STLKDTEIRDLMEAADVDN 438
CPK05 RHPWICENGVAPDRALDPAVLSRLKQFSAMNKLKKMALKVIAESLSEEEIAGLREMFQAMDTDNSGAITFDELKAGLRKYG-STLKDTEIHDLMDAADVDN 450
CPK01 CHPWVQVDGVAPDKPLDSAVLSRMKQFSAMNKFKKMALRVIAESLSEEEIAGLKEMFNMIDADKSGQITFEELKAGLKRVG-ANLKESEILDLMQAADVDN 503
510 520 530 540 550 560 570 580 590 600
CPK11 SGTIDYGEFLAATLHMNKMEREENLVAAFSYFDKDGSGYITIDELQSACTE-FG-LCDTP-LDDMIKEIDLDNDGKIDFSEFTAMMRKGD---GVGRSRTM 474
CPK12 SGTIDYGEFLAATIHLNKLEREENLVAAFSFFDKDASGYITIEELQQAWKE-FG-INDSN-LDEMIKDIDQDNDGQIDYGEFVAMMRKGNGTGGGIGRRTM 473

```
CPK26 SGTIDYGEFIAATIHLNKLEREEHLLSAFRYFDKDGSGYITIDELQHACAE-QG-MSDVF-LEDVIKEVDQDNDGRIDYGEFVAMMQKGIVGRTMRKSINM 331
CPK33 NGSIDYIEFITATMHRHRLESNENVYKAFQHFDKDGSGYITTDELEAALKE-YG-MGDDATIKEILSDVDADNDGRINYDEFCAMMRSGN--PQQPRLF-- 521
CPK21 NGTIDYYEFISATMHRYKLDRDEHVYKAFQHFDKDNSGHITRDELESAMKE-YG-MGDEASIKEVISEVDTDNDGRINFEEFCAMMRSGSTQPQG-KLLPF 530
CPK32 DGYLDCDEFIAISVHLRKMGNDEHLKKAFAFFDQNNNGYIEIEELREALSDELG--TSEEVVDAIIRDVDTDKDGRISYEEFVTMMKTGTDWRKASRQYSR 515
CPK13 KGTLDYGEFVAVSLHLQKVANDEHLRKAFSYFDKDGNGYILPQELCDALKEDGG-DDCVDVANDIFQEVDTDKDGRISYEEFAAMMKTGTDWRKASRHYSR 507
CPK08 DGTLNYGEFVAVSVHLKKMANDEHLHKAFSFFDQNQSDYIEIEELREALNDEVD-TNSEEVVAAIMQDVDTDKDGRISYEEFAAMMKAGTDWRKASRQYSR 511
CPK07 DGTLNYSEFVAVSVHLKKMANDEHLHKAFNFFDQNQSGYIEIDELREALNDELDNTSSEEVIAAIMQDVDTDKDGRISYEEFVAMMKAGTDWRKASRQYSR 513
CPK09 NGSIDYIEFITATMHRHRLESNENLYKAFQHFDKDSSGYITIDELESALKE-YG-MGDDATIKEVLSDVDSDNDGRINYEEFCAMMRSGNPQQQQPRLF-- 541
CPK04 SGTIDYGEFLAATLHINKMEREENLVVAFSYFDKDGSGYITIDELQQACTE-FG-LCDTP-LDDMIKEIDLDNDGKIDFSEFTAMMKKGDGVGRSRTMRNN 476
CPK06 SGTIDYSEFIAATIHLNKLEREEHLVSAFQYFDKDGSGYITIDELQQSCIE-HG-MTDVF-LEDIIKEVDQDNDGRIDYEEFVAMMQKGNA---GVGRRTM 533
CPK05 SGTIDYSEFIAATIHLNKLEREEHLVAAFQYFDKDGSGFITIDELQQACVE-HG-MADVF-LEDIIKEVDQNNDGKIDYGEFVEMMQKGNA---GVGRRTM 545
CPK01 SGTIDYKEFIAATLHLNKIEREDHLFAAFTYFDKDGSGYITPDELQQACEE-FG-VEDVR-IEELMRDVDQDNDGRIDYNEFVAMMQKGSITGGPVK-MGL 600

         610       620       630
CPK11 MKNLNFNIAD-------AFGVDGEKSDD 495
CPK12 RNSLNFGTTLPDESMNV----------- 490
CPK26 SIRNNAVSQ------------------- 340
CPK33 ----------------------------
CPK21 H--------------------------- 531
CPK32 ERFNSISLKLMQDA---SLQVNGDTR-- 538
CPK13 GRFNSLSIKLMKDG---SLNLGNE---- 528
CPK08 ERFNSLSLKLMREG---SLQLEGEN--- 533
CPK07 ERFNSLSLKLMRDG---SLQLEGET--- 535
CPK09 ----------------------------
CPK04 LNFNIAEAFGVEDTSSTAKSDDSPK--- 501
CPK06 KNSLNISMRDV----------------- 544
CPK05 RNSLNISMRDA----------------- 556
CPK01 EKSFSIALKL------------------ 610
```

C

```
                10        20        30        40        50        60        70        80        90       100
BSK01 MGCCQSLFSGDNPLGKDGVQPQPLSQNNHGGA--------TTADNGGSGGASGVGGGGGGGGIP-SFSEFSFADLKAATNNFSSDNIVSESGEKAPNLVYK  92
BSK02 MGCLHS------KTANLPSSDDP----SAPNK--------PESVNGDQ----------VDQEIQ-NFKEFELNELRKATNGFSPSCIVSEGGEKAPNVVYR  72
BSK03 MGGQCS------SLSC-CRNTSH--KTAVLEA--------PDVDNGES---------SEITDVP-NFREYTLEQLKAATSGFAVEYIVSEHGEKAPNVVYK  74
BSK04 MGGQSS------KIGT-C--CSH--KTTALEA--------PDVENKEN---------GEVNGVH-SFREYSLEQLKIATSCFALENVVSEHGETAPNVVYQ  72
BSK05 MGPRCS------KLSL-CWWPTHL-KSTHNEA--------SDLDNGTD-------------DLP-SFTEFSFDQLRAATCGFSTDSIVSEHGVKAPNVVYK  71
BSK06 MGARCS------KFSF-CLFPSHFKSASVLES--------PDIENGGK-------------VWP-TFKEFKLEQLKSATGGFSSDNIVSEHGEKAPNVVYR  72
BSK07 MGCEVS------KLCAFCCVSDP--EGSNHGV--------TGLDEDRR---------GEGNDLP-QFREFSIETLRNATSGFATENIVSEHGEKAPNVVYK  75
BSK08 MGCEVS------KLSALCCVSES--GRSNPDV--------TGLDEEGR---------GESNDLP-QFREFSIETIRNATSGFAAENIVSEHGERAPNVVYK  75
BSK09 ------------MI-----------ERTNLIV--------LAADNKEE---------DEGSTCP-NFLEFSLEQLRVATDGFSADNIVSEHNERVPNIVYK  60
BSK10 MGCICF------KSWRRSSSSPSITSTVIDDL--------ENVREYDA--------DDDGGHYPLIFREFSLEQLRIATDGFSAGNIVSEHNDSVPNIVYK  79
BSK11 MGCCQSSFLKPSSLHDKKITSDDLSGRRGKGAKRGNRHRHANINEGRG---------WHFSDVP-DFSEFSASVLRDATNNFNKNAVVSVCSDQEPNLVYQ  91
  SSP MGCCYS----------LSSTVDPVQDHTTDAS--------SEPRNGGG-------------EDP-PLTKFSFSALKTATNHFSPENIVS---DQTSDVVFK  66

               110       120       130       140       150       160       170       180       190       200
BSK01 GRL---QNRRWIAVKKFTKMAWPEPKQFAEEAWGVGKLRHNRLANLIGYCCDGDERLLVAEFMPNDTLAKHLFHWENQTIEWAMRLRVGYYIAEALDYCST 190
BSK02 GKL---EGNHLVAIKRFSRQSWPDAQQFVVEATGVGKLRNKRIVSLIGCCAEGDERLLVAEYMPNDTLSKHLFHWEKQPLPWDMRVRIADYIAEALDYCNI 170
BSK03 GKL---ENQKKIAVKRFTRMAWPDSRQFLEEARSVGQLRSERMANLLGCCCEGDERLLVAEFMPNETLAKHLFHWETQPMKWTMRLRVVLYLAQALEYCTS 172
BSK04 GKL---ENHMKIAIKRFSGTAWPDPRQFLEEARLVGQLRSKRMANLLGYCCEGGERLLVAEFMPNETLAKHLFHWDTEPMKWAMRLRVALYISEALEYCSN 170
BSK05 GRL---EDDRWIAVKRFNRSAWPDTRQFLEEAKAVGQLRNERLANLIGFCCEGDERLLVAEFMPFETLSKHLFHWDSQPMKWSMRLRVALYLAQALEYCSS 169
BSK06 GRL---DDGRLIAVKRFNRLAWADHRQFLDEAKAVGSLRSDRLANLIGCCFEGEERLLVAEFMPHETLAKHLFHWENNPMKWAMRLRVALCLAQALEYCSN 170
BSK07 GKL---DNQRRIAVKRFNRKAWPDSRQFLEEAKAVGQLRNYRMANLLGCCYEGEERLLVAEFMPNETLAKHLFHWESQPMKWAMRLRVALHIAQALEYCTG 173
BSK08 GKL---ENQRRIAVKRFNRKSWPDSRQFLEEAKAVGQLRNHRMANLLGCCYEDEERLLIAEFMPNETLAKHLFHWESQPMKWAMRLRVALHIAQALEYCTS 173
BSK09 GQL---NDGRKIAVKRFQRLSWPDSLEFIEEAQAVGRCRSEHMANLIGCCSEGHERLLVAEYMPNETLAKHLFHWEKRPMKWEMRLRVALHTATALEYCND 158
BSK10 GKL---GDGRRIAVKRFQRLSWPDPFEFINEAQAVGRLRSEHMANLIGCCCDDNERLLVAEYMPNGTLAKHLFHWEKRPMKWEMRLKVALHTARALEYCND 177
BSK11 GCIRSDKDKRLIAVKKFSKTTWPDPKQFATEARAIGSLRHVRLVNLIGYCCEGDERLLVSEYMPNESLTKHLFHWEKQTMEWAMRLRVALYVAEALEYCRQ 192
  SSP GRL---QNGGFVAIKRFNNMAWSDPKLFLEEAQRVGKLRHKRLVNLIGYCCDGDKRFLVADFMANDTLAKRLFQRKYQTMDWSIRLRVAYFVAEALDYCNT 164

               210       220       230       240       250       260       270       280       290       300
BSK01 EGRPLYHDLNAYRVLFDEDGDPRLSCFGLMKNSRDGKSYSTNLAYTPPEYLRNGRVTPESVTYSFGTVLLDLLSGKHIPPSHALDMIRGKNIILLMDSHLE 291
BSK02 ENRKIYHDLNAYRILFDEEGDPRLSTFGLMKNSRDGKSYSTNLAYTPPEFLRTGRVIPESVIFSYGTILLDLLSGKHIPPSHALDIIRGKNALLLMDSSLE 271
BSK03 KGRTLYHDLNAYRVLFDEECNPRLSTFGLMKNSRDGKSYSTNLAFTPPEYLRTGRITPESVIYSFGTLLLDLLSGKHIPPSHALDLIRDRNLQTLTDSCLD 273
BSK04 NGHTLYHDLNAYRVLFDEECNPRLSTFGLMKNSRDGKSYSTNLAFTPPEYLRTGRITAESVIYSFGTLLLDLLTGKHIPPSHALDLIRDRNLQTLTDSCLE 271
BSK05 KGRALYHDLNAYRILFDQDGNPRLSCFGLMKNSRDGKSYSTNLAFTPPEYLRTGRVIPESVVYSFGTLLLDLLSGKHIPPSHALDLIRGKNFLMLMDSCLD 270
BSK06 KGRALYHDLNAYRVLFDKDGNPRLSCFGLMKNSRDGKSYSTNLAFTPPEYLRTGRVTPESVVFSFGTVLLDLMSGKHIPPSHALDLIRGKNCAMLMDSALE 271
BSK07 KGRALYHDLNAYRVLFDDDSNPRLSCFGLMKNSRDGKSYSTNLAFTPPEYLRTGRVTPESVMYSYGTLLLDLLSGKHIPPSHALDLIRDRNIQMLIDSCLE 274
BSK08 KGRALYHDLNAYRVLFDDDANPRLSCFGLMKNSRDGKSYSTNLAFTPPEYLRTGRVTPESVIYSFGTLLLDLLSGKHIPPSHALDLIRDRNIQMLMDSGLE 274
BSK09 WGIDLYHDLNTYRILFDKVGNPRLSCFGLMKCSREGKSYSTNLAFAPPEYLRLGTVIPESVTFSFGTLLLDLMSGRHIPPNHALDLFRGKNYLVLMDSALD 259
BSK10 KGIDLYHDLNPYRIMFDKTGIPKLSCFGLMKNSHEGKIYSTNLAFAPPEYLRLGTVIAESVTFSFGTLLLDLMSGRHIPPNHALDLFRGKNYLVLMDSALD 278
BSK11 SGLKLYHDLNTCRVLFDENGSPRLSCFGWMKNSKDGKNFSTNLAYTPPEYLRSGTLIPESVVFSFGTFLLDLLSGKHIPPSHAVGTIQKQNLNVLMDSHLE 293
  SSP AGFASYNNLSAYKVLFDEDGDACLSCFGLMKEINNDQ-------------ITTGSVNPENVIYRFGTVLVNLLSGKQIPPSHAPEMIHRKNVFKLMDPYLK 252

               310       320       330       340       350       360       370       380       390       400
BSK01 GKFSTEEATVVVELASQCLQYEPRERPNTKDLVATLAPLQTKSDVPSYVML--------GIKKQEEAPSTPQRPLSPLGEACSRMDLTAIHQILVMTHY-R 383
BSK02 GQYANDDATKLVDLASKCLQSEAKDRPDTKFLLSAVAPLQKQEEVASHVLM--------GLPKNTVILPT---MLSPLGKACAKMDLATFHDILLKTGY-R 360
BSK03 GQFSDSDGTELVRLASRCLQYEARERPNTKSLVTALTPLQKETEVLSHVLM--------GLPHSGSVS-----PLSPLGEACSRRDLTAMLEILEKLGY-K 360
BSK04 GQFSDSDGTELVRLTSCCLQYEARERPNIKSLVTALISLQKDTEVLSHVLM--------GLPQSGTFAS----PPSPFAEACSGKDLTSMVEILEKIGY-K 359
BSK05 GHFSNDDGTDLVRLASRCLQYEARERPNVKSLVSSLAPLQKETDIPSHVLM--------GIPHGAASPKETT-SLTPLGDACSRHDLTAIHEILEKVGY-K 361
BSK06 GHFSNEDGTELVRLATRCLQYEARERPNVKSLVTSLVTLQKESDVASYVLM--------GIPHETEAEEESPLSLTPFGDACLRVDLTAIQEILSKIGY-K 363
BSK07 GQFSSDDGTELIRLASRCLQYEPRERPNPKSLVTAMIPLQKDLETPSHQLM--------GIPSSASTT-----PLSPLGEACLRTDLTAIHEILEKLSY-K 361
BSK08 GQFSSDDGTELIRLASRCLQYEPRERPNPKSLVSAMIPLQKDLEIASHQLL--------GVPNSATTT-----ALSPLGEACLRSDLTAIHEIIEKLGY-K 361
BSK09 GQFSDEDRTELIHLASRCLRPEPDERPSIKFLMSALSRLEKRAELWPNVKEENI-----PTPSYTEPATKEPLPLTPFGEACWRVDLSGMHELLEKLGYGE 355
BSK10 GQFSDEDRTELIHVASRCFKTEPEERPSIKFLKATLSRLQKRAKLCPINVKRPMSPPSKNLPEKTKPAT-ESLKLTPFGDACSRADLSSIHELLEKLGY-E 377
BSK11 GNYPEEDAAMVFDLASKCLHNNPNERPEIGDIISVITTLQQKLDVPSYTML--------GISKLEKLEMEH--PKSLIYDACHQMDLAALHQILEAMEY-K 383
  SSP GKFSIDEANVVYKLASQCLKYEGQESPNTKEIVATLETLQTRTEAPSYEVV--------EMTNQEKDASSSS-NLSPLGEACLRMDLASIHSILVLAGY-D 343

               410       420       430       440       450       460       470       480       490       500
BSK01 DDEG-TNELSFQEWTQQMKDMLDARKRGDQSFREKDFKTAIDCYSQFIDVGTMVSPTVFGRRSLCYLLCDQPDAALRDAMQAQCVYPDWPTAFYMQSVALA 483
BSK02 DEEGAENELSFQEWTQQVQEMLNTKKFGDIAFRDKDFKNSIEYYSKLVGMMPVPSATVFARRAFSYLMTDQQELALRDAMQAQVCIPEWPTAFYLQALALS 461
BSK03 DDEGVTNELSFHMWTDQMQESLNSKKKGDVAFRQKDFREAIECYTQFIDGG-MISPTVCARRSLCYLMSDMPKEALDDAIQAQVISPVWHVASYLQSASLG 460
BSK04 DDE----DLSF-MWTEQMQEAINSKKKGDIAFRRKDFSEAIEFYTQFLDLG-MISATVLVRRSQSYLMSNMAKEALDDAMKAQGISPVWYVALYLQSAALS 454
BSK05 DDEGVANELSFQVWTDQIQETLNSKKQGDAAFKGKDFVTAVECYTQFIEDGTMVSPTVFARRCLCYLMSNMPQEALGDAMQAQVVSPEWPTAFYLQAAALF 462
BSK06 DDEGIANELSFQMWTNQMQESLNSKKQGDLAFRSKDFTTAVDCYTQFIDGGTMVSPTVHARRCLSYLMNDNAQEALTDALQAQVVSPDWPTALYLQAACLF 464
BSK07 DDEGAATELSFQMWTNQMQDSLNFKKKGDVAFRHKEFANAIDCYSQFIEGGTMVSPTVYARRSLCYLMNEMPQEALNDAMQAQVISPAWHIASYLQAVALS 462
BSK08 DDEGATTELSFQMWTDQMQDTLVFKKKGDSAFRHKDFAKAIECYSQFIEVGTMGSPTVHARQSLCYLMNDMPREALNNAMQAQVISPAWHIASYLQAVALS 462
BSK09 DDVVVTNEFSFQMWTGQMQENMDYKKHGDAAFRAKDFETAIEFYTEFMSGAPVVSPTVLARRCLCYLMSDMFREALSDAMQTQVASPEFSIALYLQAACLL 456
BSK10 EDNGVGNEFSFQMWTGEMQENMDYKKHGDAAFLAKDFDTAIEFYTEFMTGAPTVSPTVLARRCLCYLMTEMFSEALSDAMQAQVASPEWPIPLYLQAACLF 478
BSK11 EDE-VTCELSFQQWAQQIKDVCNTRQQGDSAFRNKHFESAIDKYTQFIEIGIMISPTVYARRSMCYLFCDQPDAALRDAMQAQCVYSDWPTAFYLQAVALS 483
  SSP DDKDII-ELSFEEWIQEVKELQDVRRNGDRAFVEQDFKTAIACYSQFVEERSLVYPSVYARRSLSYLFCDEPEKALLDGMHAQGVFPDWPTAFYLQSVALA 443

               510       520       530
BSK01 KLNMNTDAADMLNEAAQLEEKRQRGGRGS  512
BSK02 KLGMETDAQDMLNDGAAYDAKRQNSWRC-  489
BSK03 ILGMEKESQIALKEGSNLEAKMNGVPRVK  489
BSK04 VLGMEKESQIALTEGSILEARKISASTQN  483
BSK05 SLGMDKDACETLKDGTSLEAKKHNNRN--  489
BSK06 KLGMEADAQQALKDGTTLEAKKSNKR---  490
BSK07 ALGQENEAHAALKDGSMLESKRNRL----  487
BSK08 ALGQENEAHTALKDGAMLESKRNPL----  487
BSK09 KLGMEAEAKEALRHGSSLEAF--------  477
BSK10 KLEMEAEAKEALRHGSALEAY--------  499
BSK11 KLNMVEDSATMLKEALILEDKRGS-----  507
  SSP KLDMNTDSADTLKEAALLEVKK-------  465
```

D

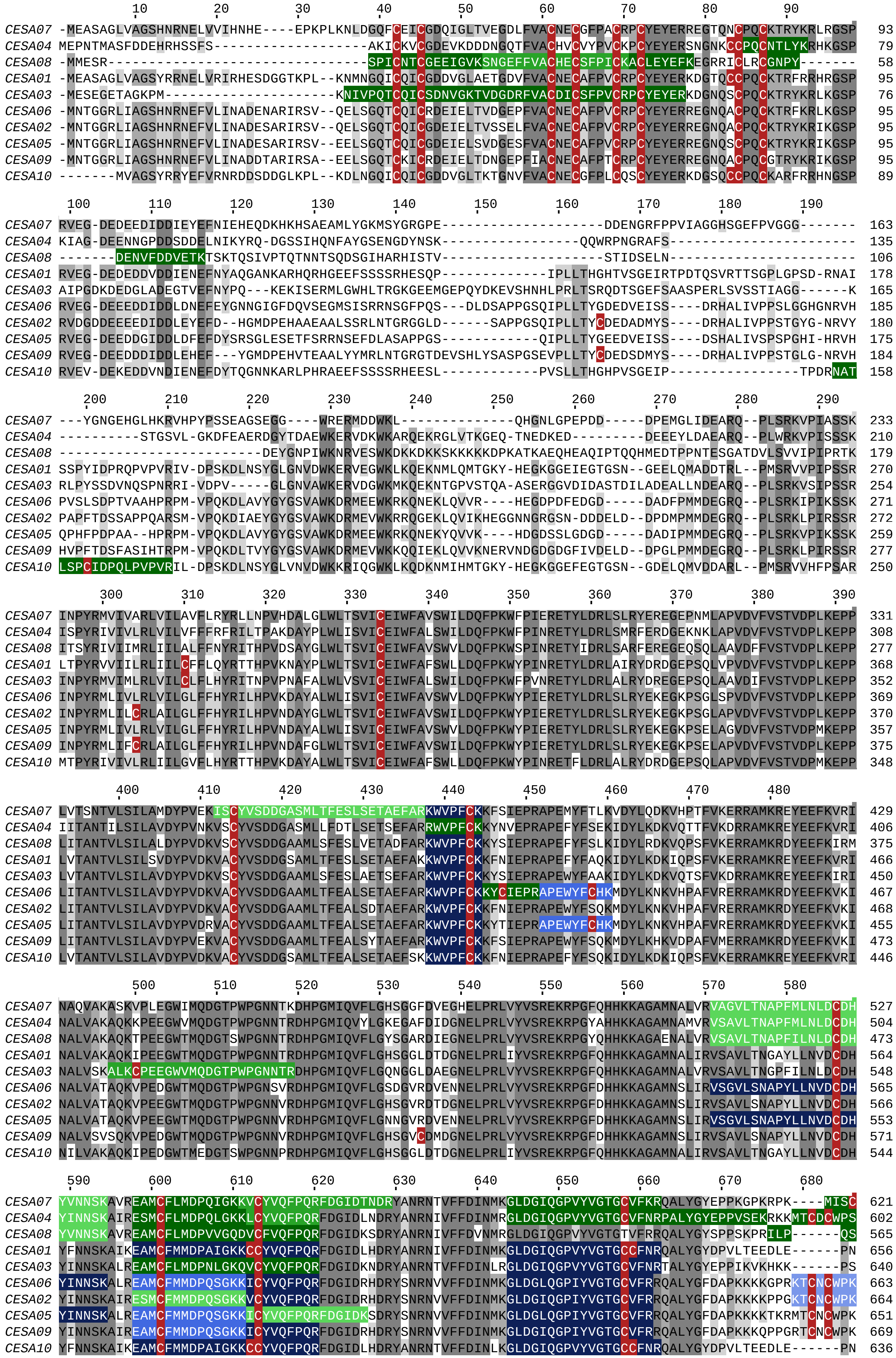
CESA07
CESA04
CESA08
CESA01
CESA03
CESA06
CESA02
CESA05
CESA09
CESA10

```
              690       700       710       720       730       740       750       760       770       780
CESA07 GCCPCFG------------------------RRRKNKKFS---------KNDMNGDVAALGGAEGD------------KEHLMSEMNFEKTFGQSSIF  674
CESA04 WICCCCGGGNRNHKSDSSKKKSGIKSLFSKLKKKTKKKSDDKTMSSYSRKRSSTEAIFDLEDIEEG--LEG--YDELEKSSLMSQKNFEKRFGMSPVF  696
CESA08 SSSSCCC------------------------LTKKKQPQDPSEIYKDAKREELDAAIFNLGDLDN--------YDEYDRSMLISQTSFEKTFGLSTVF  631
CESA01 IIVKSCC-GSR--------------------KKGKSSKKYNYEKRRGINRSDSNAPLFNMEDIDEG--FEG--YDD-ERSILMSQRSVEKRFGQSPVF  728
CESA03 LLSKLCG-GSR--------------------KKNSKAKKES-DKKKSGRHTDSTVPVFNLDDIEEG--VEGAGFDD-EKALLMSQMSLEKRFGQSAVF  713
CESA06 WCLLCFG------------------------SRKNRKAKTVAAD-KKKKNREASKQIHALENIEEGRVTKG---SNVEQSTEAMQMKLEKKFGQSPVF  733
CESA02 WCCLCCG------------------------LRKKSKTKA--KD-KKTNTKETSKQIHALENVDEGVIVP---VSNVEKRSEATQLKLEKKFGQSPVF  732
CESA05 WCLFCCG------------------------LRKNRKSKT--TD-KKKKNREASKQIHALENIEEG--TKG--TNDAAKSPEAAQLKLEKKFGQSPVF  718
CESA09 WCCLCCG------------------------MRKKKTGKV--KDNQRKKPKETSKQIHALEHIEEG--LQ---VTNAENNSETAQLKLEKKFGQSPVL  736
CESA10 IIVKSCF-GSR--------------------KKGKSRKIPNYEDNRSIKRSDSNVPLFNMEDIDED--VEG--YED-EMSLLVSQKRLEKRFGQSPVF  708

                790       800       810       820       830       840       850       860       870       880
CESA07 VTSTLMEEGGVPPSSSPAVLLKEAIHVISCGYEDKTEWGTELGWIYGSITEDILTGFKMHCRGWRSIYCMPKRPAFKGSAPINLSDRLNQVLRWALGS  772
CESA04 IASTLMENGGLPEATNTSSLIKEAIHVISCGYEEKTEWGKEIGWIYGSVTEDILTGFRMHCRGWKSVYCMPKRPAFKGSAPINLSDRLHQVLRWALGS  794
CESA08 IESTLMENGGVPDSVNPSTLIKEAIHVISCGYEEKTEWGKEIGWIYGSITEDILTGFKMHCRGWRSIYCMPLRPAFKGSAPINLSDRLHQVLRWALGS  729
CESA01 IAATFMEQGGIPPTTNPATLLKEAIHVISCGYEDKTEWGKEIGWIYGSVTEDILTGFKMHARGWISIYCNPPRPAFKGSAPINLSDRLNQVLRWALGS  826
CESA03 VASTLMENGGVPPSATPENLLKEAIHVISCGYEDKSDWGMEIGWIYGSVTEDILTGFKMHARGWRSIYCMPKLPAFKGSAPINLSDRLNQVLRWALGS  811
CESA06 VASARMENGGMARNASPACLLKEAIQVISCGYEDKTEWGKEIGWIYGSVTEDILTGFKMHSHGWRSVYCTPKLAAFKGSAPINLSDRLHQVLRWALGS  831
CESA02 VASAVLQNGGVPRNASPACLLREAIQVISCGYEDKTEWGKEIGWIYGSVTEDILTGFKMHCHGWRSVYCMPKRAAFKGSAPINLSDRLHQVLRWALGS  830
CESA05 VASAGMENGGLARNASPASLLREAIQVISCGYEDKTEWGKEIGWIYGSVTEDILTGFKMHSHGWRSVYCTPKIPAFKGSAPINLSDRLHQVLRWALGS  816
CESA09 VASTLLLNGGVPSNVNPASLLRESIQVISCGYEEKTEWGKEIGWIYGSVTEDILTGFKMHCHGWRSVYCMPKRAAFKGSAPINLSDRLHQVLRWALGS  834
CESA10 IAATFMEQGGLPSTTNPLTLLKEAIHVISCGYEAKTDWGKEIGWIYGSVTEDILTGFKMHARGWISIYCVPSRPAFKGSAPINLSDRLNQVLRWALGS  806

                  890       900       910       920       930       940       950       960       970
CESA07 VEIFFSRHSPLWYGYKGGKLKWLERFAYANTTIYPFTSIPLLAYCILPAICLLTDKFIMPPISTFASLFFISLFMSIIVTGILELRWSGVSIEEWWRN  870
CESA04 VEIFFSRHCPLWYAWG-GKLKILERLAYINTIVYPFTSIPLLAYCTIPAVCLLTGKFIIPTINNFASIWFLALFLSIIATAILELRWSGVSINDLWRN  891
CESA08 VEIFLSRHCPLWYGCSGGRLKLLQRLAYINTIVYPFTSLPLVAYCTLPAICLLTGKFIIPTLSNLASMLFLGLFISIILTSVLELRWSGVSIEDLWRN  827
CESA01 IEILLSRHCPIWYGYH-GRLRLLERIAYINTIVYPITSIPLIAYCILPAFCLITDRFIIPEISNYASIWFILLFISIAVTGILELRWSGVSIEDWWRN  923
CESA03 VEILFSRHCPIWYGYN-GRLKFLERFAYVNTTIYPITSIPLLMYCTLPAVCLFTNQFIIPQISNIASIWFLSLFLSIFATGILEMRWSGVGIDEWWRN  908
CESA06 VEIFLSRHCPIWYGYG-GGLKWLERLSYINSVVYPWTSLPLIVYCSLPAICLLTGKFIVPEISNYASILFMALFSSIAITGILEMQWGKVGIDDWWRN  928
CESA02 VEIFLSRHCPIWYGYG-GGLKWLERFSYINSVVYPWTSLPLIVYCSLPAVCLLTGKFIVPEISNYAGILFMLMFISIAVTGILEMQWGGVGIDDWWRN  927
CESA05 VEIFLSRHCPIWYGYG-GGLKWLERLSYINSVVYPWTSIPLLVYCSLPAICLLTGKFIVPEISNYASILFMALFGSIAVTGILEMQWGKVGIDDWWRN  913
CESA09 VEIFLSRHCPIWYGYG-GGLKWLERFSYINSVVYPWTSLPLLVYCSLPAICLLTGKFIVPEISNYAGILFLLMFMSIAVTGILEMQWGKIGIDDWWRN  931
CESA10 IEILLSRHCPIWYGYN-GRLKLLERIAYINTIVYPITSIPLLAYCMLPAFCLITNTFIIPEISNLASLCFMLLFASIYASAILELKWSDVALEDWWRN  903

                   990      1000      1010      1020      1030      1040      1050      1060      1070
CESA07 EQFWVIGGISAHLFAVVQGLLKILAGIDTNFTVTSK--ATDDDDFGELYAFKWTTLLIPPTTVLIINIVGVVAGISDAINNGYQSWGPLFGKLFFSFW  966
CESA04 EQFWVIGGVSAHLFAVFQGLLKVLFGVDTNFTVTSKGASDEADEFGDLYLFKWTTLLIPPTTLIILNMVGVVAGVSDAINNGYGSWGPLFGKLFFAFW  989
CESA08 EQFWVIGGVSAHLFAVFQGFLKMLAGLDTNFTVTSK--TADDLEFGELYIVKWTTLLIPPTSLLIINLVGVVAGFSDALNKGYEAWGPLFGKVFFAFW  923
CESA01 EQFWVIGGTSAHLFAVFQGLLKVLAGIDTNFTVTSK-ATDEDGDFAELYIFKWTALLIPPTTVLLVNLIGIVAGVSYAVNSGYQSWGPLFGKLFFALW 1020
CESA03 EQFWVIGGVSAHLFAVFQGILKVLAGIDTNFTVTSK-ASDEDGDFAELYLFKWTTLLIPPTTLLIVNLVGVVAGVSYAINSGYQSWGPLFGKLFFAFW 1005
CESA06 EQFWVIGGVSAHLFALFQGLLKVLAGVDTNFTVTSK--AADDGEFSDLYLFKWTSLLIPPMTLLIINVIGVIVGVSDAISNGYDSWGPLFGRLFFALW 1024
CESA02 EQFWVIGGASSHLFALFQGLLKVLAGVNTNFTVTSK--AADDGAFSELYIFKWTTLLIPPTTLLIINIIGVIVGVSDAISNGYDSWGPLFGRLFFALW 1023
CESA05 EQFWVIGGVSAHLFALFQGLLKVLAGVETNFTVTSK--AADDGEFSELYIFKWTSLLIPPTTLLIINVIGVIVGISDAISNGYDSWGPLFGRLFFAFW 1009
CESA09 EQFWVIGGVSSHLFALFQGLLKVLAGVSTNFTVTSK--AADDGEFSELYIFKWTSLLIPPTTLLIINIVGVIVGVSDAINNGYDSWGPLFGRLFFALW 1027
CESA10 EQFWVIGGTSAHLFAVFQGLLKVFAGIDTNFTVTSK-ASDEDGDFAELYVFKWTSLLIPPTTILLVNLVGIVAGVSYAINSGYQSWGPLMGKLLFAFW 1000

          1080      1090      1100      1110      1120      1130      1140
CESA07 VIVHLYPFLKGLMGRQNRTPTIVVIWSVLLASIFSLLWVRIDPFVLKTKGPDTSKCG--INC--                                   1026
CESA04 VIVHLYPFLKGLMGRQNRTPTIVVLWSILLASIFSLVWVRIDPFLPKQTGPLLKQCG--VDC--                                   1049
CESA08 VILHLYPFLKGLMGRQNRTPTIVILWSILLASVFSLVWVRINPFVSKTDTTSLSLNCLLIDC--                                    985
CESA01 VIAHLYPFLKGLLGRQNRTPTIVIVWSVLLASIFSLLWVRINPFVDANP-NANNFNG--KGGVF                                   1081
CESA03 VIVHLYPFLKGLMGRQNRTPTIVVVWSVLLASIFSLLWVRIDPFTSRVTGPDILECG--INC--                                   1065
CESA06 VIIHLYPFLKGLLGKQDRMPTIIVVWSILLASILTLLWVRVNPFVAKGG-PILEICG--LDCL-                                   1084
CESA02 VIVHLYPFLKGMLGKQDKMPTIIVVWSILLASILTLLWVRVNPFVAKGG-PVLEICG--LNCGN                                   1084
CESA05 VILHLYPFLKGLLGKQDRMPTIILVWSILLASILTLLWVRVNPFVAKGG-PILEICG--LDCL-                                   1069
CESA09 VIVHLYPFLKGLLGKQDRVPTIILVWSILLASILTLLWVRVNPFVSKDG-PVLEICG--LDCLK                                   1088
CESA10 VVAHLYPFLKGLLGRQNRTPTIVIVWSALLASIFSLLWVRINPFVSTTGVMSNSFMG--E----                                   1058
```

**Figure S3. S acylation sites in selected protein families**

Protein sequences for ROPs (A), CPKs (B), BSKs (C) and CESAs (D) were aligned using MUSCLE algorithm with default settings in Jalview. Peptides identified as being modified by S-acylation are highlighted. The exclusive peptide matches are indicated by three different shades of green while the ambiguous peptides are shown with shades of blue. The highest confidence groups are represented by the darkest shades. All cysteines residues are marked with red text. For CPKs and BSKs, the kinase domain is indicated by a red rectangle. For CPKs, only 14 CPKs that were identified in the data are shown. Shades of grey indicate sequence conservation.

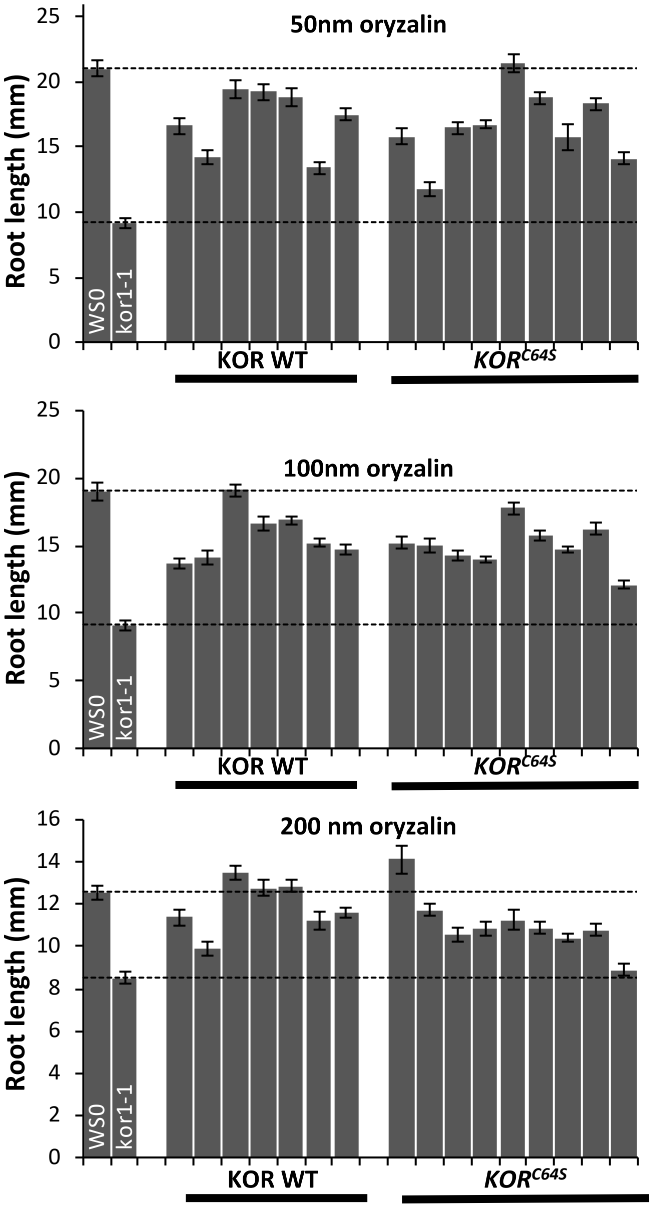

**Figure S4. S-Acylation of KOR1**

GFP fusions of WT or the *KOR*$^{C64S}$ mutant were expressed in *kor1-1* under control of the native promoter. Plants were grown on vertical plates containing the concentrations of oryzalin specified. At least 30 roots were measured for each line. Bars shown are standard error of mean. The dashed horizontal lines indicate root lengths of Ws and *kor1-1*.

| Group | Motif | Score | FG | BG | Fold Enrichment | Unadjusted p-value | Tests | Adjusted p-value |
|---|---|---|---|---|---|---|---|---|
| No TMH | xxVxQx_C_xxxxxx | 24.08 | 25/4235 | 2/4235 | 12.5 | 2.70E-06 | 446 | 0.001 |
| No TMH | xKxxQx_C_xxxxxx | 15.25 | 21/4235 | 5/4235 | 4.2 | 1.20E-03 | 437 | 0.420 |
| No TMH | Kxxxxx_C_xxxxxx | 7.8 | 375/4235 | 281/4235 | 1.3 | 7.70E-05 | 444 | 0.033 |
| No TMH | xxxCxx_C_xxxxxx | 6.32 | 166/4235 | 113/4235 | 1.5 | 7.50E-04 | 397 | 0.260 |
| TMH | xxxxxx_C_xxPxxx | 4.59 | 69/879 | 41/879 | 1.7 | 3.80E-03 | 200 | 0.530 |
| TMH | xxxxxx_C_xxxxxK | 4.74 | 73/879 | 46/879 | 1.6 | 6.60E-03 | 193 | 0.720 |
| TMH | xxxxxx_C_xxCxxx | 4.35 | 51/879 | 30/879 | 1.7 | 1.10E-02 | 191 | 0.880 |
| TMH | xxxxxF_C_xxxxxx | 4.3 | 55/879 | 32/879 | 1.7 | 7.60E-03 | 185 | 0.750 |

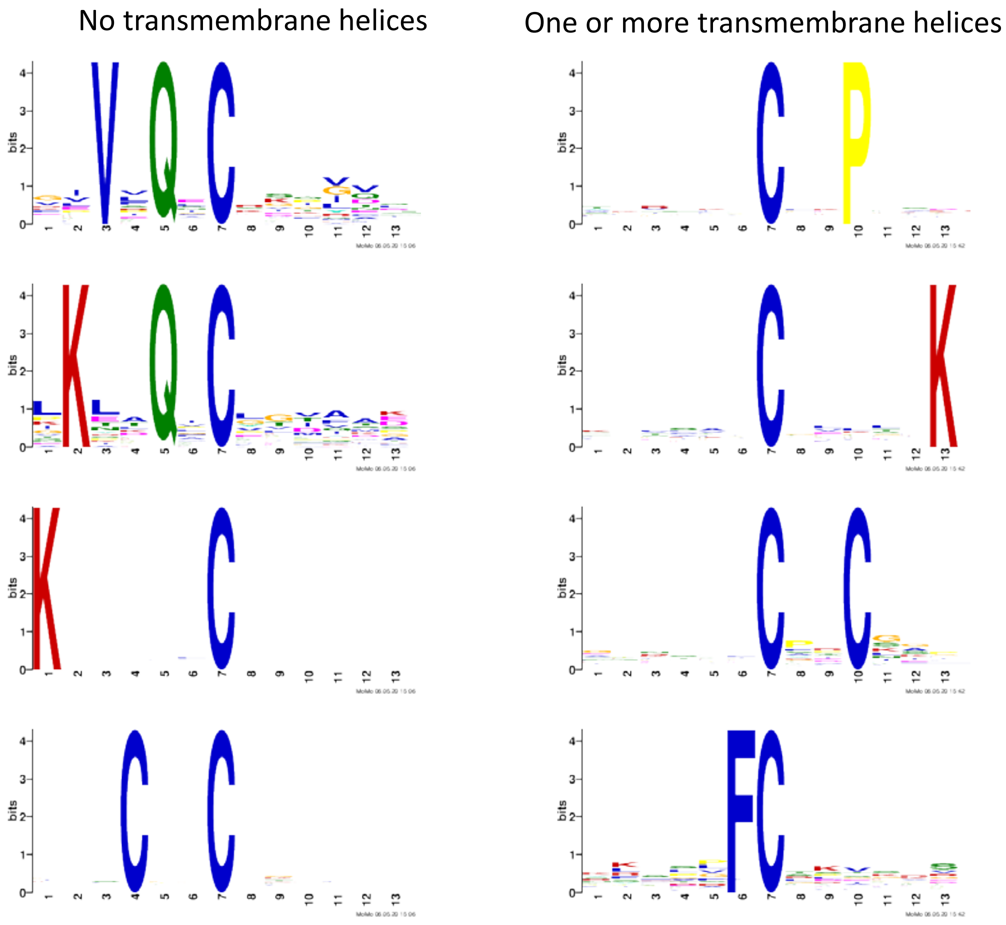

**Figure S5. Motif Analysis**

Motif enrichment analysis was performed using the motif-x algorithm using a 13 amino acid long motif containing the central cysteine and 6 residues on either side. Peptides generated from entire Araport proteome were used as background. Analysis was performed separately for high confidence cysteines from proteins that either lacked or contained one of more transmembrane domains. Both the scores (top) and Weblogo representations of the motif (bottom) are shown.

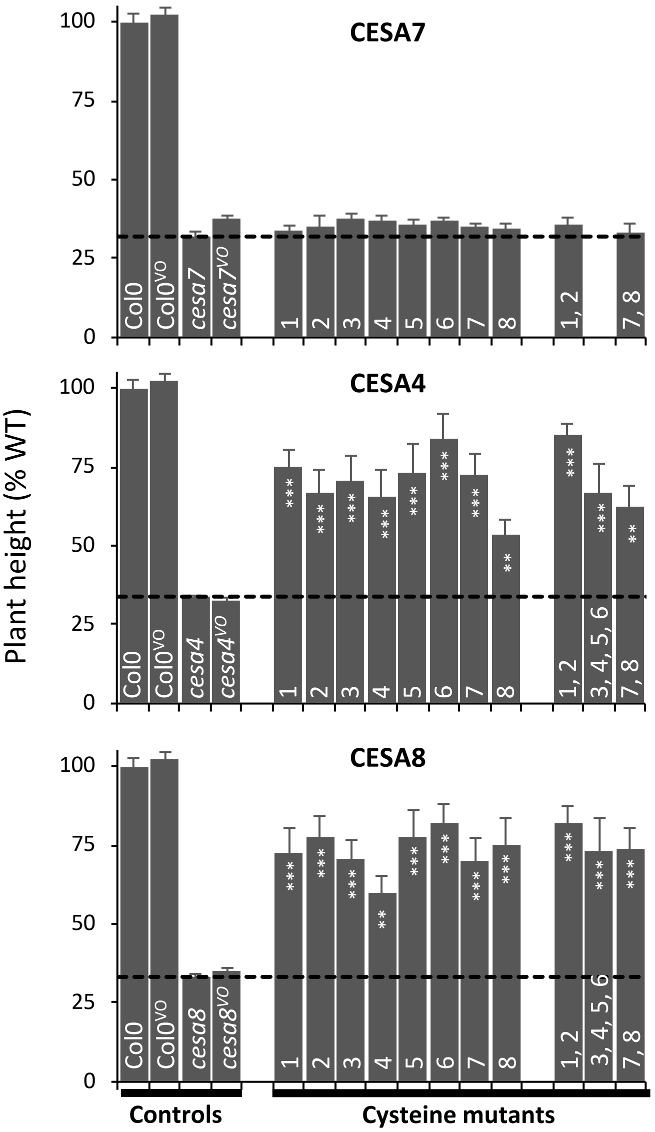

**Figure S6. Complementation analysis using CESA protein amino terminus cysteines mutants**

Constructs in which one or more of the 8 conserved cysteines from the amino terminus of Arabidopsis CESA7, CESA4and CESA8 have been mutated to serines were transformed into T-DNA null mutants: *cesa7*$^{irx3\text{-}7}$, *cesa4*$^{irx5\text{-}4}$ and *cesa8*$^{irx1\text{-}7}$ respectively and analysed for plant height. For each genotype at least 9 plants were analysed. Col0, *cesa7*$^{irx3\text{-}7}$, *cesa4*$^{irx5\text{-}4}$ and *cesa8*$^{irx1\text{-}7}$ and transformed with an empty vector (VO) are also included. The numbers of the cysteines mutated are indicated at the base of each bar and correspond to those indicated in Fig. 5A. Horizontal dashed line indicates plant height of background mutants. Error bars shown are standard error of mean. Significance levels at 0.001 (***), 0.01 (**) and 0.05 (*) are shown for comparison of cysteine mutants with the respective T-DNA mutants and were generated using univariate ANOVA.

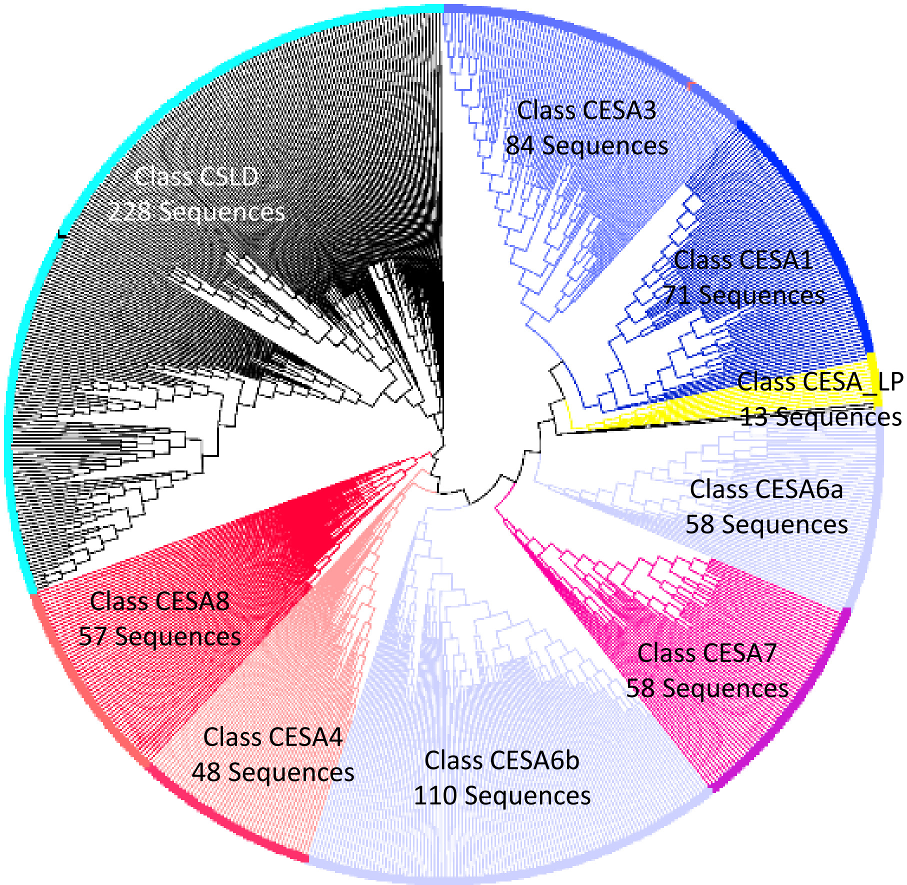

**Figure S7. CESA phylogeny based on amino terminal domain.**
Full length protein sequences of CESA and CSLD proteins from 43 fully sequenced plant genomes were aligned with MUSCLE and a ML tree produced. Using the clades of full length tree, CESA sequences were placed into different classes. The amino terminal domains (Pfam Zf-UDP) were extracted by taking the sequence between the first and the last of the 8 cysteine residues. These sequences were then aligned and used to generate a maximum likelihood tree. A subset of these, which included only those sequences that were constituted of exactly 46 amino acids (the number of residues for all 10 CESAs from Arabidopsis), were used to produce the weblogos shown in figure Fig 5c.

| LOCUS | NAME | Group | Tissue | | | | | | |
|---|---|---|---|---|---|---|---|---|---|
| | | | STEM | HYP | SLQ | MERIS | SDL | LEAF | Best |
| AT4G32410 | CESA01 | 1_CESA | B1 | B1 | B2 | B1 | B1 | B2 | B1 |
| AT4G39350 | CESA02 | 1_CESA | A1 | B1 | A3 | B1 | A3 | B2 | A1 |
| AT5G05170 | CESA03 | 1_CESA | A1 | A1 | A1 | A2 | A1 | A1 | A1 |
| AT5G44030 | CESA04 | 1_CESA | A1 | A1 | A1 | A3 | B2 | B2 | A1 |
| AT5G09870 | CESA05 | 1_CESA | B1 | B1 | B1 | A3 | B1 | B1 | A3 |
| AT5G64740 | CESA06 | 1_CESA | A3 | B1 | B1 | A1 | A3 | B1 | A1 |
| AT5G17420 | CESA07 | 1_CESA | A2 | A1 | A1 | A1 | B1 | B3 | A1 |
| AT4G18780 | CESA08 | 1_CESA | A1 | A1 | A2 | A1 | B1 | NA | A1 |
| AT2G21770 | CESA09 | 1_CESA | B1 | B1 | B1 | B2 | B1 | B1 | B1 |
| AT2G25540 | CESA10 | 1_CESA | B1 | B1 | A1 | B1 | B1 | B2 | A1 |
| AT5G49720 | KOR | 2_KOR | A1 | A1 | A1 | A1 | A1 | A2 | A1 |
| AT1G45688 | CC1 | 3_Cellulose-MT | NA | NA | NA | NA | NA | NA | NA |
| AT5G42860 | CC2 | 3_Cellulose-MT | NA | NA | NA | NA | NA | NA | NA |
| AT2G41990 | CC3 | 3_Cellulose-MT | NA | NA | NA | NA | NA | NA | NA |
| AT4G35170 | CC4 | 3_Cellulose-MT | NA | NA | NA | NA | NA | NA | NA |
| AT4G10840 | CMU1 | 3_Cellulose-MT | A3 | B1 | A2 | A2 | A3 | NA | A2 |
| AT3G27960 | CMU2 | 3_Cellulose-MT | NA | NA | NA | NA | NA | NA | NA |
| AT1G27500 | CMU3 | 3_Cellulose-MT | A1 | B1 | A3 | A3 | NA | NA | A1 |
| AT2G22125 | CSI1 | 3_Cellulose-MT | A2 | A1 | A1 | A2 | A1 | A1 | A1 |
| AT1G44120 | CSI2 | 3_Cellulose-MT | NA | NA | NA | NA | NA | NA | NA |
| AT1G77460 | CSI3 | 3_Cellulose-MT | A3 | NA | NA | A3 | NA | NA | A3 |
| AT1G12840 | DET3 | 4_Trafficking1 | A1 | A1 | A2 | A1 | A1 | A1 | A1 |
| AT5G06970 | PATROL1 | 4_Trafficking1 | A2 | A3 | A3 | NA | NA | NA | A2 |
| AT2G41770 | STELLO1 | 4_Trafficking1 | NA | NA | NA | NA | NA | NA | NA |
| AT3G57420 | STELLO2 | 4_Trafficking1 | NA | NA | NA | NA | NA | NA | NA |
| AT5G46630 | AP2M | 4_Trafficking2_endo | NA | A1 | A1 | A2 | A3 | A3 | A1 |
| AT2G44610 | RabH1B | 4_Trafficking2_endo | NA | NA | NA | NA | NA | NA | NA |
| AT5G57460 | TML | 4_Trafficking2_endo | A2 | A1 | A3 | A1 | A2 | NA | A1 |
| AT3G01780 | TPLATE | 4_Trafficking2_endo | A1 | A1 | A1 | A2 | A1 | A1 | A1 |
| AT5G24710 | TWD40-2 | 4_Trafficking2_endo | A1 | A1 | A1 | A1 | A3 | NA | A1 |
| AT5G03540 | Exo70A1 | 4_Trafficking3_exo | A1 | A3 | NA | NA | A2 | NA | A1 |
| AT5G58430 | Exo70B1 | 4_Trafficking3_exo | NA | NA | NA | NA | NA | NA | NA |
| AT5G49830 | Exo84B | 4_Trafficking3_exo | A2 | A1 | A1 | A1 | A1 | NA | A1 |
| AT1G21170 | Sec5B | 4_Trafficking3_exo | B1 | NA | NA | NA | B3 | NA | B1 |
| AT1G71820 | Sec6 | 4_Trafficking3_exo | NA | NA | A1 | A1 | NA | NA | A1 |
| AT1G78880 | SHOU4 | 4_Trafficking3_exo | A2 | A3 | B1 | B1 | B1 | B1 | A2 |
| AT1G16860 | SHOU4L | 4_Trafficking3_exo | A3 | B1 | B1 | B1 | B1 | B1 | A3 |
| AT4G22290 | SHOU4L2 | 4_Trafficking3_exo | NA | NA | NA | NA | NA | NA | NA |
| AT2G33100 | CslD1 | 5_CSLD | NA | NA | NA | NA | NA | NA | NA |
| AT5G16910 | CslD2 | 5_CSLD | B1 | A2 | A2 | B4 | B2 | B2 | A2 |
| AT3G03050 | CslD3 | 5_CSLD | B1 | B1 | A3 | B4 | A1 | B2 | A1 |
| AT4G38190 | CslD4 | 5_CSLD | NA | NA | NA | NA | NA | NA | NA |
| AT1G02730 | CslD5 | 5_CSLD | B1 | B1 | B1 | A3 | B1 | B1 | A3 |

**Table S2. List of proteins associated with cellulose biosynthesis**

The list of cellulose synthesis realted proteins includes CESAs, CSLDs and proteins listed by Anderson and Kieber, 2020. Protein class of each protein in various tissues is shown. A1, A2, A3 and B1, B2, B3 refer to high, medium and low confidence groups based on exclusive and ambiguous peptides respectively. NA means not detected.

| Level 1 code | Level 2 Code | Level 2 description | REF | Tissue | | | | | | |
|---|---|---|---|---|---|---|---|---|---|---|
| | | | | All | ST | HC | SQ | MS | SL | LF |
| 1 | 1.1 | Photosynthesis.photophosphorylation | 212 | 46 | 30 | 5 | 28 | 17 | 24 | 25 |
| 1 | 1.2 | Photosynthesis.calvin cycle | 40 | 21 | 16 | 4 | 15 | 7 | 12 | 13 |
| 1 | 1.3 | Photosynthesis.photorespiration | 27 | 8 | 5 | 2 | 3 | 2 | 3 | 1 |
| 1 | 1.4 | Photosynthesis.CAM/C4 photosynthesis | 9 | 3 | 2 | 1 | 3 | 0 | 1 | 1 |
| 2 | 2.1 | Cellular respiration.glycolysis | 54 | 29 | 19 | 12 | 20 | 7 | 17 | 5 |
| 2 | 2.2 | Cellular respiration.pyruvate oxidation | 9 | 2 | 1 | 1 | 2 | 1 | 2 | 1 |
| 2 | 2.3 | Cellular respiration.tricarboxylic acid cycle | 38 | 11 | 6 | 6 | 6 | 5 | 6 | 7 |
| 2 | 2.4 | Cellular respiration.oxidative phosphorylation | 141 | 19 | 9 | 9 | 9 | 10 | 10 | 7 |
| 3 | 3.1 | Carbohydrate metabolism.sucrose metabolism | 51 | 18 | 10 | 6 | 7 | 5 | 11 | 5 |
| 3 | 3.2 | Carbohydrate metabolism.starch metabolism | 45 | 21 | 12 | 11 | 12 | 8 | 9 | 7 |
| 3 | 3.3 | Carbohydrate metabolism.trehalose metabolism | 15 | 2 | 0 | 1 | 1 | 1 | 1 | 0 |
| 3 | 3.4 | Carbohydrate metabolism.raffinose family oligosaccharide biosynthesis | 10 | 1 | 1 | 0 | 0 | 0 | 1 | 0 |
| 3 | 3.5 | Carbohydrate metabolism.sorbitol metabolism | 1 | 1 | 1 | 1 | 1 | 1 | 1 | 1 |
| 3 | 3.6 | Carbohydrate metabolism.mannose metabolism | 7 | 1 | 0 | 0 | 0 | 1 | 0 | 0 |
| 3 | 3.7 | Carbohydrate metabolism.galactose metabolism | 1 | 1 | 1 | 1 | 1 | 1 | 1 | 1 |
| 3 | 3.8 | Carbohydrate metabolism.nucleotide sugar biosynthesis | 65 | 20 | 12 | 10 | 14 | 7 | 8 | 5 |
| 3 | 3.9 | Carbohydrate metabolism.fermentation | 6 | 0 | 0 | 0 | 0 | 0 | 0 | 0 |
| 3 | 3.10 | Carbohydrate metabolism.oxidative pentose phosphate pathway | 22 | 7 | 2 | 1 | 5 | 3 | 1 | 2 |
| 3 | 3.11 | Carbohydrate metabolism.gluconeogenesis | 6 | 2 | 1 | 1 | 1 | 0 | 0 | 0 |
| 4 | 4.1 | Amino acid metabolism.biosynthesis | 166 | 57 | 28 | 27 | 28 | 30 | 28 | 16 |
| 4 | 4.2 | Amino acid metabolism.degradation | 54 | 16 | 9 | 7 | 5 | 10 | 5 | 4 |
| 5 | 5.1 | Lipid metabolism.fatty acid synthesis | 116 | 33 | 12 | 10 | 21 | 18 | 16 | 6 |
| 5 | 5.2 | Lipid metabolism.glycerolipid synthesis | 54 | 10 | 5 | 2 | 5 | 7 | 4 | 3 |
| 5 | 5.3 | Lipid metabolism.galactolipid and sulfolipid synthesis | 9 | 2 | 0 | 0 | 1 | 2 | 1 | 1 |
| 5 | 5.4 | Lipid metabolism.sphingolipid metabolism | 38 | 3 | 0 | 2 | 1 | 2 | 0 | 1 |
| 5 | 5.5 | Lipid metabolism.phytosterols | 33 | 7 | 6 | 4 | 5 | 4 | 4 | 2 |
| 5 | 5.6 | Lipid metabolism.lipid A synthesis | 10 | 0 | 0 | 0 | 0 | 0 | 0 | 0 |
| 5 | 5.7 | Lipid metabolism.lipid degradation | 112 | 22 | 10 | 13 | 10 | 8 | 13 | 7 |
| 5 | 5.8 | Lipid metabolism.lipid transport | 31 | 5 | 2 | 2 | 3 | 0 | 2 | 1 |
| 5 | 5.9 | Lipid metabolism.lipid bodies-associated activities | 40 | 2 | 0 | 0 | 1 | 2 | 0 | 0 |
| 6 | 6.1 | Nucleotide metabolism.purines | 51 | 17 | 9 | 3 | 12 | 9 | 8 | 5 |
| 6 | 6.2 | Nucleotide metabolism.pyrimidines | 36 | 13 | 7 | 5 | 8 | 8 | 5 | 2 |
| 6 | 6.3 | Nucleotide metabolism.deoxynucleotide metabolism | 16 | 2 | 0 | 0 | 1 | 1 | 1 | 1 |
| 7 | 7.1 | Coenzyme metabolism.molybdenum cofactor synthesis | 7 | 1 | 0 | 0 | 0 | 1 | 1 | 1 |
| 7 | 7.2 | Coenzyme metabolism.thiamine pyrophosphate synthesis | 7 | 5 | 2 | 0 | 3 | 2 | 3 | 3 |
| 7 | 7.3 | Coenzyme metabolism.S-adenosyl methionine (SAM) cycle | 6 | 4 | 3 | 3 | 2 | 2 | 4 | 2 |
| 7 | 7.4 | Coenzyme metabolism.coenzyme A synthesis | 11 | 1 | 0 | 1 | 0 | 0 | 0 | 0 |
| 7 | 7.5 | Coenzyme metabolism.tetrahydrofolate synthesis | 25 | 8 | 7 | 6 | 6 | 2 | 4 | 4 |
| 7 | 7.6 | Coenzyme metabolism.biotin synthesis | 5 | 2 | 0 | 0 | 1 | 2 | 0 | 0 |
| 7 | 7.7 | Coenzyme metabolism.pyridoxalphosphate synthesis | 5 | 3 | 2 | 3 | 2 | 2 | 2 | 0 |
| 7 | 7.8 | Coenzyme metabolism.prenylquinone synthesis | 16 | 3 | 1 | 0 | 2 | 1 | 2 | 0 |
| 7 | 7.9 | Coenzyme metabolism.NAD/NADP biosynthesis | 11 | 4 | 2 | 1 | 3 | 2 | 2 | 2 |
| 7 | 7.10 | Coenzyme metabolism.FMN/FAD biosynthesis | 12 | 4 | 3 | 0 | 0 | 2 | 1 | 0 |
| 7 | 7.11 | Coenzyme metabolism.iron-sulfur cluster assembly machineries | 45 | 13 | 4 | 1 | 5 | 8 | 4 | 1 |
| 7 | 7.12 | Coenzyme metabolism.tetrapyrrol biosynthesis | 57 | 17 | 10 | 2 | 11 | 8 | 12 | 7 |
| 7 | 7.13 | Coenzyme metabolism.phylloquinone synthesis | 10 | 1 | 0 | 1 | 1 | 1 | 1 | 0 |
| 7 | 7.14 | Coenzyme metabolism.lipoic acid synthesis | 4 | 0 | 0 | 0 | 0 | 0 | 0 | 0 |
| 8 | 8.1 | Polyamine metabolism.putrescine | 12 | 0 | 0 | 0 | 0 | 0 | 0 | 0 |
| 8 | 8.2 | Polyamine metabolism.spermidine/spermine | 13 | 2 | 1 | 1 | 1 | 0 | 0 | 0 |
| 9 | 9.1 | Secondary metabolism.terpenoids | 111 | 19 | 11 | 4 | 12 | 10 | 9 | 3 |
| 9 | 9.2 | Secondary metabolism.phenolics | 45 | 14 | 4 | 0 | 11 | 4 | 7 | 1 |
| 9 | 9.3 | Secondary metabolism.nitrogen-containing secondary compounds | 66 | 20 | 11 | 7 | 9 | 6 | 11 | 10 |
| 10 | 10.1 | Redox homeostasis.reactive oxygen generation | 12 | 1 | 1 | 0 | 0 | 0 | 0 | 0 |
| 10 | 10.2 | Redox homeostasis.enzymatic reactive oxygen species scavengers | 11 | 3 | 0 | 0 | 1 | 2 | 1 | 1 |
| 10 | 10.3 | Redox homeostasis.low-molecular-weight scavengers | 27 | 10 | 8 | 3 | 8 | 3 | 5 | 4 |
| 10 | 10.4 | Redox homeostasis.hydrogen peroxide removal | 34 | 13 | 7 | 4 | 5 | 5 | 3 | 7 |
| 10 | 10.5 | Redox homeostasis.chloroplast redox homeostasis | 24 | 7 | 5 | 2 | 6 | 4 | 5 | 5 |

| Level 1 code | Level 2 Code | Level 2 description | REF | Tissue | | | | | | |
|---|---|---|---|---|---|---|---|---|---|---|
| | | | | All | ST | HC | SQ | MS | SL | LF |
| 10 | 10.6 | Redox homeostasis.cytosol/mitochondrion/nucleus redox homeostasis | 16 | 0 | 0 | 0 | 0 | 0 | 0 | 0 |
| 11 | 11.1 | Phytohormones.abscisic acid | 72 | 8 | 2 | 3 | 3 | 7 | 5 | 1 |
| 11 | 11.2 | Phytohormones.auxin | 58 | 7 | 3 | 5 | 3 | 3 | 4 | 3 |
| 11 | 11.3 | Phytohormones.brassinosteroid | 57 | 9 | 5 | 7 | 4 | 8 | 6 | 3 |
| 11 | 11.4 | Phytohormones.cytokinin | 56 | 0 | 0 | 0 | 0 | 0 | 0 | 0 |
| 11 | 11.5 | Phytohormones.ethylene | 34 | 3 | 1 | 2 | 1 | 1 | 2 | 1 |
| 11 | 11.6 | Phytohormones.gibberellin | 34 | 1 | 0 | 1 | 1 | 1 | 1 | 1 |
| 11 | 11.7 | Phytohormones.jasmonic acid | 28 | 9 | 3 | 3 | 4 | 6 | 2 | 3 |
| 11 | 11.8 | Phytohormones.salicylic acid | 12 | 0 | 0 | 0 | 0 | 0 | 0 | 0 |
| 11 | 11.9 | Phytohormones.strigolactone | 17 | 0 | 0 | 0 | 0 | 0 | 0 | 0 |
| 11 | 11.10 | Phytohormones.signalling peptides | 215 | 9 | 1 | 3 | 3 | 7 | 0 | 1 |
| 12 | 12.1 | Chromatin organisation.histones | 48 | 1 | 0 | 0 | 1 | 0 | 0 | 0 |
| 12 | 12.2 | Chromatin organisation.histone chaperone activities | 20 | 6 | 1 | 1 | 2 | 6 | 2 | 1 |
| 12 | 12.3 | Chromatin organisation.histone modifications | 138 | 15 | 2 | 3 | 5 | 10 | 1 | 1 |
| 12 | 12.4 | Chromatin organisation.chromatin remodeling complexes | 51 | 5 | 1 | 2 | 1 | 4 | 1 | 0 |
| 12 | 12.5 | Chromatin organisation.DNA methylation | 51 | 5 | 1 | 1 | 1 | 4 | 1 | 0 |
| 13 | 13.1 | Cell cycle.regulation | 82 | 1 | 0 | 0 | 0 | 1 | 0 | 0 |
| 13 | 13.2 | Cell cycle.interphase | 84 | 12 | 3 | 1 | 2 | 9 | 1 | 0 |
| 13 | 13.3 | Cell cycle.mitosis and meiosis | 160 | 10 | 1 | 0 | 0 | 9 | 1 | 0 |
| 13 | 13.4 | Cell cycle.cytokinesis | 72 | 12 | 3 | 7 | 6 | 7 | 5 | 1 |
| 13 | 13.5 | Cell cycle.organelle machineries | 50 | 7 | 0 | 1 | 1 | 6 | 2 | 0 |
| 14 | 14.1 | DNA damage response.DNA damage sensing and signalling | 4 | 0 | 0 | 0 | 0 | 0 | 0 | 0 |
| 14 | 14.2 | DNA damage response.BRCA1–BARD1 DNA-damage response heterodimer | 3 | 0 | 0 | 0 | 0 | 0 | 0 | 0 |
| 14 | 14.3 | DNA damage response.BRCC DNA-damage response complex | 8 | 0 | 0 | 0 | 0 | 0 | 0 | 0 |
| 14 | 14.4 | DNA damage response.DNA repair polymerase activities | 7 | 0 | 0 | 0 | 0 | 0 | 0 | 0 |
| 14 | 14.5 | DNA damage response.DNA repair mechanisms | 62 | 7 | 1 | 2 | 2 | 7 | 2 | 0 |
| 15 | 15.1 | RNA biosynthesis.DNA-dependent RNA polymerase (Pol) complexes | 40 | 6 | 2 | 4 | 2 | 6 | 2 | 1 |
| 15 | 15.2 | RNA biosynthesis.RNA polymerase I-dependent transcription | 10 | 2 | 0 | 0 | 0 | 2 | 1 | 0 |
| 15 | 15.3 | RNA biosynthesis.RNA polymerase II-dependent transcription | 156 | 11 | 2 | 3 | 1 | 9 | 4 | 0 |
| 15 | 15.4 | RNA biosynthesis.RNA polymerase III-dependent transcription | 26 | 0 | 0 | 0 | 0 | 0 | 0 | 0 |
| 15 | 15.5 | RNA biosynthesis.siRNA biogenesis | 18 | 1 | 0 | 0 | 1 | 1 | 1 | 0 |
| 15 | 15.6 | RNA biosynthesis.rRNA biogenesis | 5 | 0 | 0 | 0 | 0 | 0 | 0 | 0 |
| 15 | 15.7 | RNA biosynthesis.transcriptional activation | 1926 | 25 | 2 | 5 | 5 | 17 | 6 | 1 |
| 15 | 15.8 | RNA biosynthesis.transcriptional repression | 18 | 2 | 1 | 2 | 1 | 2 | 1 | 0 |
| 15 | 15.9 | RNA biosynthesis.organelle machineries | 73 | 9 | 6 | 1 | 3 | 6 | 6 | 3 |
| 16 | 16.1 | RNA processing.RNA 5'-end cap adding | 0 | 0 | 0 | 0 | 0 | 0 | 0 | 0 |
| 16 | 16.2 | RNA processing.RNA 3'-end polyadenylation | 0 | 0 | 0 | 0 | 0 | 0 | 0 | 0 |
| 16 | 16.3 | RNA processing.RNA quality control Exon Junction complex (EJC) | 12 | 3 | 1 | 1 | 1 | 2 | 1 | 1 |
| 16 | 16.4 | RNA processing.RNA splicing | 153 | 17 | 4 | 6 | 7 | 9 | 9 | 3 |
| 16 | 16.5 | RNA processing.ribonuclease activities | 17 | 0 | 0 | 0 | 0 | 0 | 0 | 0 |
| 16 | 16.6 | RNA processing.RNA editing | 2 | 0 | 0 | 0 | 0 | 0 | 0 | 0 |
| 16 | 16.7 | RNA processing.RNA modification | 93 | 19 | 0 | 3 | 4 | 13 | 7 | 0 |
| 16 | 16.8 | RNA processing.RNA decay | 45 | 12 | 3 | 5 | 1 | 9 | 4 | 1 |
| 16 | 16.9 | RNA processing.messenger ribonucleoprotein particle (mRNP) | 28 | 3 | 2 | 1 | 3 | 1 | 3 | 1 |
| 16 | 16.10 | RNA processing.organelle machineries | 103 | 5 | 1 | 0 | 2 | 3 | 3 | 1 |
| 17 | 17.1 | Protein biosynthesis.cytosolic ribosome | 294 | 36 | 11 | 12 | 19 | 13 | 17 | 7 |
| 17 | 17.2 | Protein biosynthesis.aminoacyl-tRNA synthetase activities | 56 | 23 | 16 | 14 | 20 | 15 | 19 | 10 |
| 17 | 17.3 | Protein biosynthesis.translation initiation | 81 | 19 | 8 | 9 | 14 | 12 | 11 | 4 |
| 17 | 17.4 | Protein biosynthesis.translation elongation | 27 | 6 | 4 | 2 | 5 | 4 | 3 | 5 |
| 17 | 17.5 | Protein biosynthesis.translation termination | 6 | 1 | 0 | 0 | 1 | 1 | 0 | 1 |
| 17 | 17.6 | Protein biosynthesis.organelle translation machineries | 157 | 27 | 11 | 2 | 12 | 17 | 12 | 10 |
| 18 | 18.1 | Protein modification.N-linked glycosylation | 56 | 7 | 3 | 5 | 2 | 3 | 4 | 1 |
| 18 | 18.2 | Protein modification.O-linked glycosylation | 17 | 3 | 0 | 0 | 2 | 1 | 1 | 0 |

| Level 1 code | Level 2 Code | Level 2 description | REF | Tissue | | | | | | |
|---|---|---|---|---|---|---|---|---|---|---|
| | | | | All | ST | HC | SQ | MS | SL | LF |
| 18 | 18.3 | Protein modification.hydroxylation | 13 | 0 | 0 | 0 | 0 | 0 | 0 | 0 |
| 18 | 18.4 | Protein modification.disulfide bond formation | 12 | 3 | 0 | 1 | 2 | 2 | 1 | 1 |
| 18 | 18.5 | Protein modification.ADP-ribosylation | 5 | 0 | 0 | 0 | 0 | 0 | 0 | 0 |
| 18 | 18.6 | Protein modification.acetylation | 10 | 1 | 1 | 0 | 0 | 1 | 1 | 0 |
| 18 | 18.7 | Protein modification.lipidation | 26 | 3 | 1 | 3 | 0 | 1 | 1 | 1 |
| 18 | 18.8 | Protein modification.phosphorylation | 959 | 75 | 27 | 35 | 20 | 45 | 21 | 10 |
| 18 | 18.9 | Protein modification.tyrosine sulfation | 1 | 0 | 0 | 0 | 0 | 0 | 0 | 0 |
| 18 | 18.10 | Protein modification.dephosphorylation | 150 | 20 | 4 | 11 | 9 | 12 | 7 | 1 |
| 18 | 18.11 | Protein modification.S-nitrosylation and denitrosylation | 5 | 0 | 0 | 0 | 0 | 0 | 0 | 0 |
| 18 | 18.12 | Protein modification.S-glutathionylation and deglutathionylation | 75 | 9 | 2 | 2 | 4 | 2 | 3 | 0 |
| 18 | 18.13 | Protein modification.protein folding and quality control | 78 | 16 | 7 | 4 | 6 | 6 | 8 | 6 |
| 18 | 18.14 | Protein modification.peptide maturation | 45 | 5 | 2 | 3 | 2 | 3 | 3 | 2 |
| 18 | 18.15 | Protein modification.protein repair | 2 | 0 | 0 | 0 | 0 | 0 | 0 | 0 |
| 19 | 19.1 | Protein degradation.ER-associated protein degradation (ERAD) machinery | 17 | 0 | 0 | 0 | 0 | 0 | 0 | 0 |
| 19 | 19.2 | Protein degradation.26S proteasome | 57 | 15 | 6 | 8 | 10 | 6 | 9 | 5 |
| 19 | 19.3 | Protein degradation.N-end rule pathway of targeted proteolysis | 6 | 1 | 1 | 1 | 1 | 1 | 1 | 1 |
| 19 | 19.4 | Protein degradation.peptide tagging | 567 | 32 | 12 | 17 | 21 | 21 | 18 | 5 |
| 19 | 19.5 | Protein degradation.peptidase families | 387 | 52 | 23 | 25 | 24 | 25 | 23 | 15 |
| 20 | 20.1 | Cytoskeleton.microtubular network | 116 | 11 | 5 | 3 | 7 | 4 | 5 | 3 |
| 20 | 20.2 | Cytoskeleton.microfilament network | 130 | 13 | 6 | 4 | 5 | 6 | 3 | 2 |
| 20 | 20.3 | Cytoskeleton.actin and tubulin folding | 21 | 10 | 7 | 6 | 9 | 3 | 9 | 2 |
| 20 | 20.4 | Cytoskeleton.cytoskeleton-nucleoskeleton linking | 13 | 0 | 0 | 0 | 0 | 0 | 0 | 0 |
| 20 | 20.5 | Cytoskeleton.cp-actin-dependent plastid movement | 14 | 2 | 1 | 0 | 0 | 0 | 1 | 0 |
| 20 | 20.6 | Cytoskeleton.cytoskeleton-plasma membrane-cell wall interface | 13 | 1 | 0 | 0 | 0 | 1 | 0 | 0 |
| 21 | 21.1 | Cell wall.cellulose | 37 | 10 | 5 | 6 | 5 | 5 | 3 | 2 |
| 21 | 21.2 | Cell wall.hemicellulose | 89 | 7 | 2 | 3 | 3 | 1 | 3 | 0 |
| 21 | 21.3 | Cell wall.pectin | 180 | 32 | 14 | 8 | 16 | 10 | 7 | 6 |
| 21 | 21.4 | Cell wall.cell wall proteins | 140 | 4 | 0 | 3 | 1 | 0 | 1 | 0 |
| 21 | 21.5 | Cell wall.cell wall-bound hydroxycinnamic acids | 1 | 1 | 0 | 0 | 1 | 0 | 1 | 1 |
| 21 | 21.6 | Cell wall.lignin | 37 | 10 | 4 | 4 | 3 | 2 | 4 | 1 |
| 21 | 21.7 | Cell wall.callose | 12 | 2 | 2 | 1 | 1 | 2 | 1 | 0 |
| 21 | 21.8 | Cell wall.sporopollenin | 14 | 3 | 0 | 0 | 0 | 3 | 0 | 0 |
| 21 | 21.9 | Cell wall.cutin and suberin | 75 | 12 | 3 | 1 | 5 | 4 | 3 | 0 |
| 22 | 22.1 | Vesicle trafficking.clathrin coated vesicle (CCV) machinery | 55 | 19 | 10 | 12 | 7 | 14 | 5 | 5 |
| 22 | 22.2 | Vesicle trafficking.clathrin-independent machinery | 3 | 0 | 0 | 0 | 0 | 0 | 0 | 0 |
| 22 | 22.3 | Vesicle trafficking.Coat protein I (COPI) coatomer machinery | 65 | 10 | 5 | 4 | 4 | 2 | 3 | 4 |
| 22 | 22.4 | Vesicle trafficking.Coat protein II (COPII) coatomer machinery | 22 | 13 | 9 | 5 | 7 | 6 | 6 | 4 |
| 22 | 22.5 | Vesicle trafficking.autophagosome formation | 44 | 2 | 0 | 1 | 0 | 1 | 1 | 0 |
| 22 | 22.6 | Vesicle trafficking.endomembrane trafficking | 90 | 14 | 6 | 5 | 6 | 7 | 2 | 1 |
| 22 | 22.7 | Vesicle trafficking.target membrane tethering | 85 | 21 | 17 | 13 | 9 | 14 | 11 | 6 |
| 22 | 22.8 | Vesicle trafficking.SNARE target membrane recognition and fusion complexes | 67 | 4 | 3 | 3 | 3 | 3 | 3 | 1 |
| 22 | 22.9 | Vesicle trafficking.regulation of membrane tethering and fusion | 117 | 12 | 5 | 6 | 9 | 5 | 5 | 3 |
| 23 | 23.1 | Protein translocation.chloroplast | 44 | 14 | 5 | 1 | 6 | 8 | 9 | 9 |
| 23 | 23.2 | Protein translocation.mitochondrion | 40 | 2 | 0 | 0 | 0 | 2 | 0 | 0 |
| 23 | 23.3 | Protein translocation.endoplasmic reticulum | 28 | 4 | 2 | 3 | 4 | 1 | 3 | 1 |
| 23 | 23.4 | Protein translocation.peroxisome | 16 | 3 | 2 | 1 | 1 | 1 | 2 | 0 |
| 23 | 23.5 | Protein translocation.nucleus | 70 | 23 | 13 | 8 | 12 | 15 | 10 | 4 |
| 23 | 23.6 | Protein translocation.plasmodesmata intercellular trafficking | 2 | 0 | 0 | 0 | 0 | 0 | 0 | 0 |
| 24 | 24.1 | Solute transport.primary active transport | 185 | 38 | 20 | 24 | 19 | 16 | 14 | 9 |
| 24 | 24.2 | Solute transport.carrier-mediated transport | 763 | 21 | 14 | 4 | 9 | 8 | 6 | 8 |
| 24 | 24.3 | Solute transport.channels | 178 | 9 | 5 | 1 | 1 | 4 | 3 | 3 |
| 24 | 24.4 | Solute transport.porins | 13 | 4 | 0 | 0 | 2 | 3 | 1 | 0 |
| 25 | 25.1 | Nutrient uptake.nitrogen assimilation | 48 | 12 | 9 | 8 | 9 | 4 | 11 | 6 |
| 25 | 25.2 | Nutrient uptake.sulfur assimilation | 13 | 3 | 0 | 0 | 1 | 1 | 2 | 1 |
| 25 | 25.3 | Nutrient uptake.phosphorus assimilation | 26 | 0 | 0 | 0 | 0 | 0 | 0 | 0 |

| Level 1 code | Level 2 Code | Level 2 description | REF | Tissue | | | | | | |
|---|---|---|---|---|---|---|---|---|---|---|
| | | | | All | ST | HC | SQ | MS | SL | LF |
| 25 | 25.4 | Nutrient uptake.iron uptake | 49 | 3 | 1 | 0 | 1 | 2 | 2 | 1 |
| 25 | 25.5 | Nutrient uptake.copper uptake | 23 | 3 | 1 | 2 | 0 | 0 | 2 | 1 |
| 26 | 26.1 | External stimuli response.light | 52 | 3 | 1 | 2 | 2 | 1 | 2 | 1 |
| 26 | 26.2 | External stimuli response.gravity | 11 | 2 | 0 | 1 | 1 | 1 | 2 | 0 |
| 26 | 26.3 | External stimuli response.temperature | 85 | 20 | 9 | 8 | 13 | 12 | 14 | 5 |
| 26 | 26.4 | External stimuli response.drought | 2 | 1 | 1 | 0 | 0 | 0 | 0 | 0 |
| 26 | 26.5 | External stimuli response.salinity | 8 | 2 | 2 | 2 | 0 | 1 | 1 | 0 |
| 26 | 26.6 | External stimuli response.biotic stress | 203 | 12 | 5 | 5 | 3 | 5 | 3 | 5 |
| 27 | 27.1 | Multi-process regulation.circadian clock | 33 | 0 | 0 | 0 | 0 | 0 | 0 | 0 |
| 27 | 27.2 | Multi-process regulation.TOR signalling pathway | 9 | 3 | 2 | 3 | 1 | 1 | 1 | 0 |
| 27 | 27.3 | Multi-process regulation.SnRK1 metabolic regulator system | 29 | 2 | 1 | 2 | 0 | 2 | 2 | 0 |
| 27 | 27.4 | Multi-process regulation.Rop GTPase regulatory system | 51 | 6 | 2 | 5 | 5 | 2 | 3 | 2 |
| 27 | 27.5 | Multi-process regulation.programmed cell death | 17 | 3 | 2 | 2 | 2 | 1 | 1 | 1 |
| 35 | 35.1 | not assigned.annotated | 6705 | 448 | 160 | 172 | 157 | 252 | 143 | 81 |
| 35 | 35.2 | not assigned.not annotated | 7809 | 267 | 75 | 106 | 85 | 145 | 83 | 38 |
| 50 | 50.1 | Enzyme classification.EC_1 oxidoreductases | 424 | 40 | 14 | 14 | 22 | 17 | 18 | 13 |
| 50 | 50.2 | Enzyme classification.EC_2 transferases | 421 | 34 | 12 | 11 | 8 | 14 | 21 | 8 |
| 50 | 50.3 | Enzyme classification.EC_3 hydrolases | 274 | 30 | 13 | 13 | 12 | 15 | 11 | 8 |
| 50 | 50.4 | Enzyme classification.EC_4 lyases | 31 | 4 | 0 | 1 | 1 | 2 | 1 | 0 |
| 50 | 50.5 | Enzyme classification.EC_5 isomerases | 13 | 2 | 0 | 1 | 2 | 2 | 1 | 0 |
| 50 | 50.6 | Enzyme classification.EC_6 ligases | 9 | 5 | 4 | 2 | 4 | 3 | 4 | 2 |

**Table S3. Functional classification of high confidence proteins**

Proteins identified with high confidence in at least one of the 6 tissues (All) or each individual tissue (Stem(ST), Hypocotyl (HC), Silique (SQ), Meristem (MS), Seedling(SL) and Leaf (LF) were subjected to over-representation analysis using Mapman categories. The total number of proteins for each category in whole proteome (Araport11) and the acylomes from all or individual tissues are shown. Green shading indicates over representation with a P value less than 0.05 while blue shading indicates under-representation.

| Peptide Type | Confidence Group | Prediction cutoff | | | | % predited |
| --- | --- | --- | --- | --- | --- | --- |
| | | 1_High | 2_Med | 3_low | NA | |
| **Exclusive** | **High** | 582 | 444 | 261 | 3992 | 24.4 |
| **Exclusive** | **Medium** | 222 | 150 | 103 | 1460 | 24.5 |
| **Exclusive** | **Low** | 185 | 164 | 95 | 1563 | 22.1 |
| **Ambiguous** | **High** | 189 | 158 | 117 | 1624 | 22.2 |
| **Ambiguous** | **Medium** | 72 | 45 | 27 | 456 | 24.0 |
| **Ambiguous** | **Low** | 28 | 36 | 25 | 381 | 18.9 |
| **Total** | | 1278 | 997 | 628 | 9476 | **23.5** |

**Table S4. Comparison of S-acylated cysteines identified in the data with in silico predictions** Peptides from the different confidence groups identified in this study were compared to those predicted by CSS-Palm. The final column shows the overlap between the experimental defined

| Kinase group | Number of kinases in group | Number of kinases in acylome data | | | | | | |
|---|---|---|---|---|---|---|---|---|
| | | Total | A1 | A2 | A3 | B1 | B2 | B3 |
| MAP3K | 85 | 24 | 10 | 4 | 3 | 6 | 0 | 1 |
| RLCK_07 | 50 | 19 | 9 | 6 | 2 | 1 | 1 | 0 |
| soluble | 61 | 13 | 8 | 1 | 2 | 2 | 0 | 0 |
| CDPK | 34 | 14 | 7 | 3 | 1 | 2 | 0 | 1 |
| RLCK_02 | 12 | 11 | 6 | 0 | 1 | 4 | 0 | 0 |
| LRR_03 | 44 | 9 | 3 | 3 | 1 | 2 | 0 | 0 |
| MAPK | 20 | 17 | 3 | 0 | 1 | 12 | 0 | 1 |
| LRR_08B | 54 | 6 | 2 | 1 | 3 | 0 | 0 | 0 |
| AGC | 39 | 13 | 2 | 1 | 1 | 5 | 0 | 4 |
| CDK | 30 | 5 | 2 | 1 | 1 | 1 | 0 | 0 |
| RLCK_08 | 10 | 5 | 2 | 1 | 0 | 1 | 1 | 0 |
| LRR_10 | 16 | 8 | 2 | 0 | 2 | 4 | 0 | 0 |
| LRR_06B | 15 | 7 | 2 | 0 | 0 | 5 | 0 | 0 |
| SnRK3 | 27 | 5 | 2 | 0 | 0 | 3 | 0 | 0 |
| LRR_06A | 5 | 2 | 2 | 0 | 0 | 0 | 0 | 0 |
| LRR_09A | 8 | 2 | 2 | 0 | 0 | 0 | 0 | 0 |
| LRR_11 | 38 | 2 | 2 | 0 | 0 | 0 | 0 | 0 |
| RK_01 | 20 | 2 | 2 | 0 | 0 | 0 | 0 | 0 |
| SnAK | 2 | 2 | 2 | 0 | 0 | 0 | 0 | 0 |
| SnRK1 | 3 | 3 | 1 | 1 | 0 | 1 | 0 | 0 |
| CKL | 13 | 8 | 1 | 0 | 1 | 4 | 0 | 2 |
| LRR_01 | 51 | 2 | 1 | 0 | 1 | 0 | 0 | 0 |
| RLCK_10 | 10 | 2 | 1 | 0 | 1 | 0 | 0 | 0 |
| RLCK_10A | 3 | 2 | 1 | 0 | 1 | 0 | 0 | 0 |
| LRR_08C | 22 | 3 | 1 | 0 | 0 | 2 | 0 | 0 |
| MAP2K | 10 | 3 | 1 | 0 | 0 | 2 | 0 | 0 |
| LRR_12 | 8 | 1 | 1 | 0 | 0 | 0 | 0 | 0 |
| RLCK_06 | 16 | 1 | 1 | 0 | 0 | 0 | 0 | 0 |
| LRR_02 | 14 | 8 | 0 | 1 | 0 | 2 | 5 | 0 |
| LRR_09 | 4 | 1 | 0 | 1 | 0 | 0 | 0 | 0 |
| NEK | 7 | 1 | 0 | 1 | 0 | 0 | 0 | 0 |
| L-LPK | 38 | 9 | 0 | 0 | 2 | 7 | 0 | 0 |
| SnRK2 | 10 | 9 | 0 | 0 | 2 | 7 | 0 | 0 |
| WNK | 11 | 2 | 0 | 0 | 2 | 0 | 0 | 0 |
| CK_II | 4 | 4 | 0 | 0 | 1 | 3 | 0 | 0 |
| RLCK_05 | 11 | 3 | 0 | 0 | 1 | 2 | 0 | 0 |
| PERK | 13 | 2 | 0 | 0 | 1 | 1 | 0 | 0 |
| LRR_07 | 5 | 1 | 0 | 0 | 1 | 0 | 0 | 0 |
| WAK | 26 | 1 | 0 | 0 | 1 | 0 | 0 | 0 |
| SLK | 10 | 8 | 0 | 0 | 0 | 0 | 6 | 2 |

**Table S5. Classification of protein kinases identified in the acylome**
Classification of protein kinases as described by Zulawski et al, 2014. Number of kinases identified in the acylome data within each confidence class are indicated. A1, A2, A3 and B1, B2, B3 refer to high, medium and low confidence using the exclusive and ambiguous peptide data respectively.

| Primer use | Cloning target | Forward primer sequence | Reverse primer sequence |
|---|---|---|---|
| CDS | CDS_KOR | ggggacaagtttgtacaaaaaagcaggctggATGGCTAGCTACGGAAGAG | ggggaccactttgtacaagaaagctgggtaAGTAGACTACAGTTGTTATT |
| CDS | CDS_CC1 | ggggaccactttgtacaaaaaagcaggctggATGCACGCCAAAACC | ggggaccactttgtacaagaaagctgggtaTCAAACTGTAGTGAC |
| CDS | CDS_CC2 | ggggaccactttgtacaaaaaagcaggctggATGCACGCGAAGACC | ggggaccactttgtacaagaaagctgggtaTTAAATAGATGTGAC |
| CDS | CDS_CMU1 | ggggaccactttgtacaaaaaagcaggctggATGCCAGCAATGCCA | ggggaccactttgtacaagaaagctgggtaTCAGAACTTGAAACC |
| CDS | CDS_CMU2 | ggggaccactttgtacaaaaaagcaggctggATGGACGTAGGAGAG | ggggaccactttgtacaagaaagctgggtaTCAATAAACCGGTCT |
| CDS | CDS_CMU3 | ggggaccactttgtacaaaaaagcaggctggATGGAAGGAGGGTCT | ggggaccactttgtacaagaaagctgggtaTTAACGAAGAGCTGA |
| CDS | CDS_CSI1 | ggggaccactttgtacaaaaaagcaggctggATGACAAGTGCTCTT | ggggaccactttgtacaagaaagctgggtaTTACTTGTTAGACCA |
| CDS | CDS_TUA1 | ggggaccactttgtaaaaaaagcaggctggATGAGGGAGATCATTAGCAT | ggggaccactttgtacaagaaagctgggtaCTAATACTCATCGCCTTCTT |
| CDS | CDS_TUA4 | ggggacaagtttgtacaaaaaagcaggctggATGAGAGAGTGCATTTCGATCC AC | ggggaccactttgtacaagaaagctgggtaTTAGTATTCCTCTCCTTCATCATCT |
| CDS | CDS_TUB6 | ggggacaagtttgtacaaaaaagcaggctggATGAGAGAAATCCTTCACATTC AA | ggggaccactttgtacaagaaagctgggtaTCACTCATGATCCAATATCTCTTCT |
| CDS | CDS_TUB8 | ggggaccactttgtaaaaaaagcaggctggATGCGAGAGATTCTTCACAT | ggggaccactttgtacaagaaagctgggtaTTATTGCTCCTCCTGCACTT |
| CDS | CDS_SHOU4 | ggggaccactttgtaaaaaaagcaggctggATGGGTTCGAGATACCCATC | ggggaccactttgtacaagaaagctgggtaTCAAACCGGTATGGCATCAA |
| CDS | CDS_SHOU4L | ggggaccactttgtaaaaaaagcaggctggATGGGTTCGAGATACGCATC | ggggaccactttgtacaagaaagctgggtaTCAGACAGGAATTGCATCTA |
| C>S mutant | KOR_C64S | aagtacgtcgatctcggtTCTattatcgttagccgc | complement of forward primer |
| C>S mutant | CESA4_ZNc/s_C1 | cgtcattcctcattctcggctaagattAGCaaagtc | complement of forward primer |
| C>S mutant | CESA4_ZNc/s_C2 | aaagtcTCTggcgatgaggtcaaagacgatgacaat | complement of forward primer |
| C>S mutant | CESA4_ZNc/s_C3 | gatgacaatggtcagacttttgtggcgTCTcacgtg | complement of forward primer |
| C>S mutant | CESA4_ZNc/s_C4 | acttttgtggcgtgtcacgtgAGCgtttacccggtt | complement of forward primer |
| C>S mutant | CESA4_ZNc/s_C5 | gtttacccggttAGCaaaccttgctatgaatatgag | complement of forward primer |
| C>S mutant | CESA4_ZNc/s_C6 | gtttacccggtttgcaaacctAGCtatgaatatgag | complement of forward primer |
| C>S mutant | CESA4_ZNc/s_C7 | tatgagcgtagcaacggtaacaaatgtAGCcctcaa | complement of forward primer |
| C>S mutant | CESA4_ZNc/s_C8 | cctcaaAGCaacactctttacaaacgccacaaaggc | complement of forward primer |
| C>S mutant | CESA4_ZNc/s_C1,2 | gctaagattAGCaaagTCTctggcgatgaggtcaaa | complement of forward primer |
| C>S mutant | CESA4_ZNc/s_C3,4,5,6 | gcgTCTcacgtgAGCgtttacccggttAGCaaacctAGCtat | complement of forward primer |
| C>S mutant | CESA4_ZNc/s_C7,8 | aacggtaacaaatgtAGCcctcaaAGCaacactctt | complement of forward primer |
| C>S mutant | CESA8_ZNc/s_C1 | atcAGCaacacttgtggtgaagagattggtgtaaaa | complement of forward primer |
| C>S mutant | CESA8_ZNc/s_C2 | atctgcaacactTCTggtgaagagattggtgtaaaa | complement of forward primer |
| C>S mutant | CESA8_ZNc/s_C3 | aaatcaaacggagagttctttgtggctTCTcatgag | complement of forward primer |
| C>S mutant | CESA8_ZNc/s_C4 | ttctttgtggcttgtcatgagTCTagtttcccgatc | complement of forward primer |
| C>S mutant | CESA8_ZNc/s_C5 | agtttcccgatcAGCaaagcttgtcttgagtatgaa | complement of forward primer |
| C>S mutant | CESA8_ZNc/s_C6 | agtttcccgatctgcaaagctTCTcttgagtatgaa | complement of forward primer |
| C>S mutant | CESA8_ZNc/s_C7 | tatgaattcaaagaaggtcgaagaattAGCttgcgt | complement of forward primer |
| C>S mutant | CESA8_ZNc/s_C8 | ttgcgtAGCggcaatccttacgatgagaatgtgttt | complement of forward primer |
| C>S mutant | CESA8_ZNc/s_C1,2 | tctcccatcAGCaacactTCTggtgaagagattggt | complement of forward primer |
| C>S mutant | CESA8_ZNc/s_C3,4,5,6 | gctTCTcatgagTCTagtttcccgatcAGCaaagctTCTctt | complement of forward primer |
| C>S mutant | CESA8_ZNc/s_C7,8 | agaattAGCttgcgtAGCggcaatccttacgatgag | complement of forward primer |
| C>S mutant | CESA7_ZNc/s_C1 | ctagatggacaattcTCTgagatatgtggagatcag | complement of forward primer |
| C>S mutant | CESA7_ZNc/s_C2 | caattctgtgagataTCTggagatcagattggttta | complement of forward primer |
| C>S mutant | CESA7_ZNc/s_C3 | gacctcttcgtagctAGCaatgagtgtggttttccg | complement of forward primer |
| C>S mutant | CESA7_ZNc/s_C4 | gtagcttgcaatgagTCTggttttccggcgtgtaga | complement of forward primer |
| C>S mutant | CESA7_ZNc/s_C5 | tgtggttttccggcgTCTagaccttgctatgagtac | complement of forward primer |
| C>S mutant | CESA7_ZNc/s_C6 | ccggcgtgtagacctAGCtatgagtacgagagaaga | complement of forward primer |
| C>S mutant | CESA7_ZNc/s_C7 | gaaggaacacaaaacTCTcctcagtgtaagactcgt | complement of forward primer |
| C>S mutant | CESA7_ZNc/s_C8 | caaaactgtcctcagTCTaagactcgttacaagcgt | complement of forward primer |
| C>S mutant | CESA7_ZNc/s_C1,2 | gatggacaattcTCTgagataTCTggagatcagatt | complement of forward primer |
| C>S mutant | CESA7_ZNc/s_C7,8 | ggaacacaaaacTCTcctcagTCTaagactcgttac | complement of forward primer |
| C>S mutant | CESA7$_{VR2}$ | atgataagcTCTggtTCTTCTcctAGCtttgggcgc | complement of forward primer |
| C>S mutant | CESA7$_{CT}$ | acttccaagTCTggcatcaacAGCtgaagcaaaatc | complement of forward primer |
| C>S mutant | CESA4$_{VR2}$ | TCTgatTCTtggccgtcgtggatcTCCTCTTCTAGC | complement of forward primer |
| C>S mutant | CESA4$_{CT}$ | ggtccgttactgaagcaaTCTggcgtcgacTCTtaa | complement of forward primer |
| C>S mutant | CESA8$_{VR2}$ | cttcatcatcgtcgTCTAGCTCTctaaccaagaaga | complement of forward primer |
| C>S mutant | CESA8$_{CT}$ | tccctttctctgaacTCTcttttgatcgatAGCtaa | complement of forward primer |
| Vector | pCESA8 | atcggtaccgggcccAAACCCATAACTTTAGT | atcactagtCTTCGAATTCCCCTGT |
| Vector | pCESA4 | atcggtaccggcgcgcccCGGTTTTTGGTTTGATTT | atcactagtGGCGAGGTACACTGAGCTCTCGG |
| Vector | pCESA7 | atggtaccttaattaatcgAGAGCCCGAGTCTACCTATTG | gatcaattgAGGGACGGCCGGAGATTAGCAGCGATCTTGA |
| Vector | pKOR | gattggtaccgggcccACATAGCTGCCCATATATTT | gattctagaGATGATGCTCTCTGATA |
| Vector | pCC1 | gatggcgcgccCTTCCATGGAAACTCTCCAC | gattctagaTGTTTTCCGATTTGGGAAGGTG |
| Vector | pCC2 | gatggtaccggcgcgccTCTTGTCTAGCAAATTCCAG | gatactagtCTTGAGATTGGAATGGAAGAT |
| Vector | pCMU1 | gatggcgcgccTAGGTTAGACCAGTGGTGCC | gatactagtGAATGTGTCTCTCTGTGGGAAG |
| Vector | pCMU2 | gatggtaccggcgcgccAATATACAAAAATAAAAATCC | gatactagtGGCCTCCAAAACTCACAACTCAAT |
| Vector | pCSI1 | gatggtaccggcgcgccCCATTGTTATGGGCTAATTAA | gatactagtATCTTCACTTCACTTAAAAAAATTCTC |
| Vector | pTUB6 | gatggtaccggcgcgccTGGGAAAAAAATAAATTAAG | gattctagaCTTCTATTTTATCTGAAATCAAC |
| Vector | pTUB8 | gatggtaccggcgcgccGTATCTAAACACCAGCCTA | gatactagtCTTTGATTAGTAAGCTAGAGTATG |
| Vector | GW1 | atcggtaccactagtACAAGTTTGTACAAAAAAGC | atctctagaAAACCACTTTGTACAAGAAAGCTGAA |
| Vector | p35S | gatggtaccgggcccggcgcgccACTAGAGCCAAGCTGATCTCCT | gatactagtcaattgTCGACTAGAATAGTAAATTGTAATGTTGTT |
| Vector | tNOS | gattctagaGAATTTCCCCGATCGTTCAAACATTTGGCAA | gatcctgcaggttaattaaCGAATTCCCGATCTAGTAACA |
| Vector | RGS-6xHIS-FLAG-EGFP | gatactagtATGGAAGCTAGCAGGGGATCCCATCACCATCACCATCAC GACTACAAAGATGACGATGACAAGATGGTGAGCAAGGGCGAGGAG CTGT | gattctagaCTTGTACAGCTCGTCCATGCCGAGAGTGA |
| Vector | RGS-6xHIS-STREP-EYFP-GW1 | atccaattgactagtATGAGAGGATCCCATCACCATCA | atctctagaAAACCACTTTGTACAAGAAAGCTGAA |
| Vector | VND7-VP16-GR | ggggacaagtttgtacaaaaaagcaggctggATGGATAATATAATGCAATCGT CA | ggggaccactttgtacaagaaagctgggtaTCATTTTTGATGAAACAGAA |
| Vector | p35S-UTR | gatggtaccgggcccggcgcgccGGAAACCTCCTCGGA | gatactagtATCGAATTTGGGCAGAATATACAGAAGCT |
| Vector | UTR-tNOS | gattctagaTTAACTCTGGTTTCATTAAA | gatcctgcaggGATCTAGTAACATAGATG |
| Vector | p35S-P19-t35S | gatgggcccgaattcgagctcggtaccCCCCTACTCCAAA | gatggcgcgccATCTTTTATCTTTAGAGTTAAGAACTCTT |

**Table S6. List of all primers used in this study**

Primers used for cloning of CDS fragments, cysteine mutants (C>S mutants) and the components of the destination vectors are shown. For CDS primers, the sequence of gateway adapter sequences is shown in lower case, for cysteine mutant primers, the codons for serine are shown in upper case while for the vector component primers, restriction sites used are shown in lower case.
